# Engineering a *Sinorhizobium meliloti* chassis with monopartite, single replicon genome configuration

**DOI:** 10.1101/2022.05.23.493018

**Authors:** Marcel Wagner, Johannes Döhlemann, David Geisel, Patrick Sobetzko, Javier Serrania, Peter Lenz, Anke Becker

## Abstract

Multipartite bacterial genomes pose challenges for genome engineering and establishment of additional replicons. We simplified the tripartite genome structure (3.65 Mbp chromosome, 1.35 Mbp megaplasmid pSymA, 1.68 Mbp chromid pSymB) of *Sinorhizobium meliloti*. Strains with bi- and monopartite genome configurations were generated by targeted replicon fusions. Our design preserved key genomic features, such as replichore ratios, GC skew, and KOPS and coding sequence distribution. Under standard culture conditions, growth rates of these strains and the wild type were nearly comparable. Spatiotemporal replicon organization and segregation were maintained in the triple replicon fusion strain. Deletion of the replication initiator-encoding genes including the *oriVs* of pSymA and pSymB from this strain resulted in a monopartite genome with *oriC* as the sole origin of replication, a strongly unbalanced replichore ratio, slow growth and an aberrant cellular localization of *oriC*. Suppressor mutation R436H in the cell cycle histidine kinase CckA and a 3.2 Mbp inversion, both individually, largely restored growth. These strains will facilitate integration of secondary replicons in *S. meliloti,* and thus be useful for genome engineering applications, such as generating hybrid genomes.

**Graphical Abstract:** 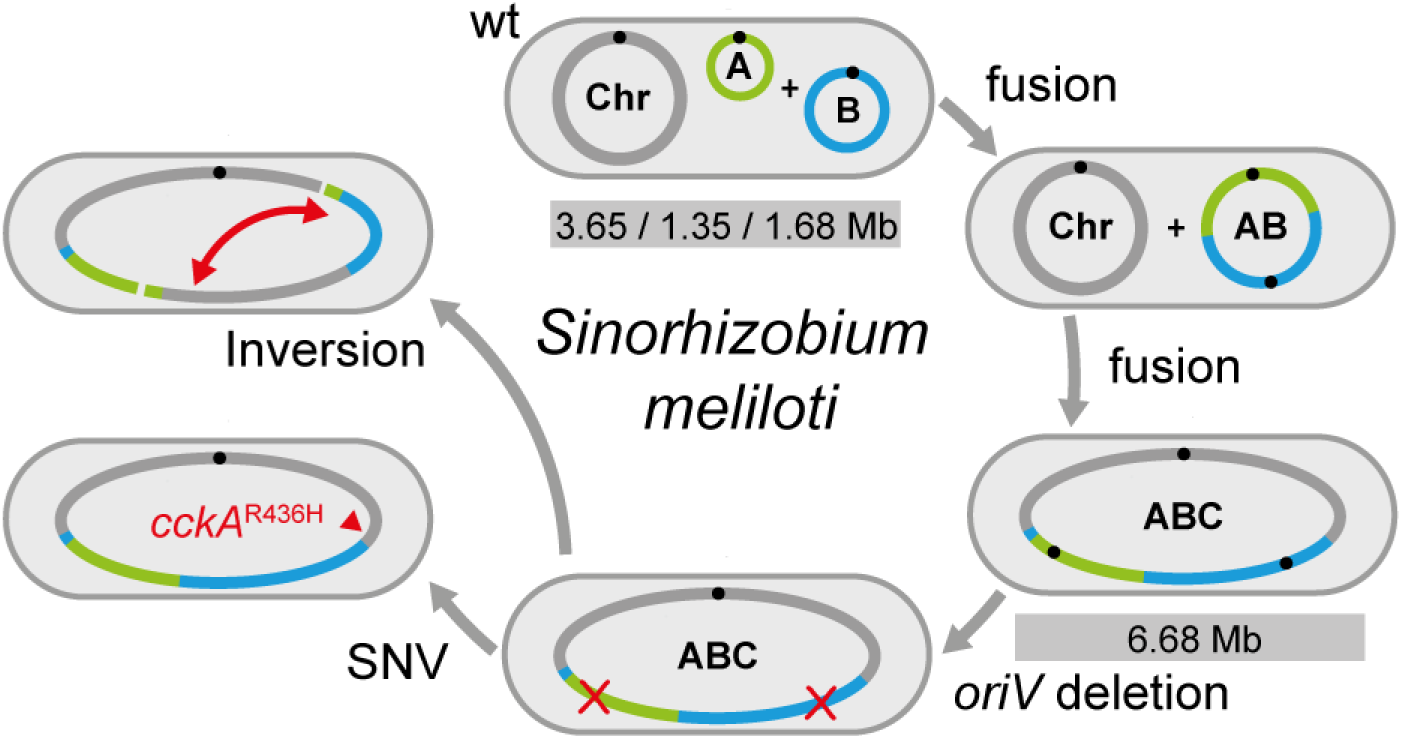

## INTRODUCTION

It is estimated that about 10 % of the bacterial species represented in genome databases carry their genomic information divided on several replicons, i.e. have multipartite genomes. *^(1)^* Their genome architectures are diverse in terms of number and size of DNA molecules. In addition to the main chromosome, these bacteria harbor one or more large secondary replicons classified as secondary chromosomes, chromids or megaplasmids according to their origin, presence of core essential genes, and size. *^(1,2)^* The presumed advantages of multipartite genomes include promotion of horizontal gene transfer, genome plasticity, and exploitation of new ecological niches. *^(2,3)^*

Several members of the Pseudomonadota with segmented genomes, such as *Agrobacterium tumefaciens* (*Rhizobium radiobacter, Agrobacterium fabrum*), *Methylorubrum extorquens*, *Sinorhizobium meliloti* (*Ensifer meliloti*), and *Vibrio natriegens*, serve as established chassis in synthetic biology for basic and applied research, and are already used in or of increasing interest to industrial biotechnology. *^(4–9)^* All these fields of research and application have in common that new or modified genetic material has to be established in the bacterial cell, often using extrachromosomal replicons (ECRs). However, this genome engineering is complicated by a multipartite genome configuration. Multiple replicons can, for instance, restrict the establishment of further ECRs through incompatibility, or lead to genomic reorganization and instability through recombination events. The advantages of a streamlined genomic architecture with fewer replicons became evident from comparing two natural *M. exorquens* isolates, leading to strain AM1 (chromosome, 4 ECRs) increasingly being replaced by strain PA1 (chromosome, no ECRs) in research and application. *^(10)^*

Fundamentally different approaches to the rational streamlining of multipartite genomes are the reduction of the number of replicons through replicon curing or the fusion of replicons, with both approaches also being combinable. The former approach is associated with the loss of genetic information, while the latter approach can retain the entire genetic information.

*S. meliloti*, a soil-dwelling diazotrophic representative of α-proteobacteria with multipartite genomes, exemplifies both approaches, either through targeted genetic interventions or genetic events in natural isolates. The *S. meliloti* type strain Rm1021 carries a main chromosome of 3.65 Mbp, megaplasmid pSymA of 1.35 Mbp, and chromid pSymB of 1.68 Mbp. *^(11)^* Vertical transmission of the secondary replicons is mediated by replicon-specific *repABC* loci *^(12)^* that provide a complete replication and segregation system. *^(13)^* The *repABC* operon codes for the DNA replication initiator protein RepC and the ParAB-type partitioning proteins RepA and RepB. Together with centromer-like sequences (*parS*) these partitioning proteins are crucial for segregation of the replicated DNA. *^(14)^* The replication origin of *repABC*-type secondary replicons (*oriV*) is located within an AT rich region of *repC*. *^(15)^* In contrast, the chromosomal origin of replication (*oriC*) is characterized by DnaA-boxes and DNA unwinding elements (DUE). *^(16)^*

Strains cured for pSymA, pSymB or both secondary replicons were generated, with the latter resulting in a genome reduction of 45 %. *^(17)^* However, the lack of pSymB is associated with considerable fitness loss. *^(17)^* In addition, the loss of secondary replicon-harbored genes can strongly limit the application range of replicon-cured strains, as these genes are e.g. important for symbiotic nitrogen fixation as well as for fitness and a broad substrate spectrum in the natural habitat. *^(18,19)^* Yet, such strains have already been used as a platform for the construction of hybrid genomes by complementation with secondary replicons from closely related strains (*in-cis* hybrids). *^(20)^* Cointegration events of two or three replicons, resulting in DNA molecules with two or three origins of replication, have been reported for natural strain isolates of the α-proteobacteria *S. meliloti, Sinorhizoium fredii* and *A*. *tumefaciens*, as well as for the *β*-proteobacterium *Cupriavidus necator* and the γ-proteobacterium *Vibrio cholerae*. *^(21–27)^*

In bacterial cells, the genomic DNA is highly organized, and replication and segregation are coordinated in space and time. *^(28)^* Tethering of the chromosomal origin and terminus regions to cell poles plays an important role for spatial chromosome organization and segregation. *^(29)^* Previously, a longitudinal *oriC* (old pole)-*terC* (new pole) configuration of the main chromosome and subpolar localization (old pole) of the *ori*s of the secondary replicons was reported for *S. meliloti* G_1_-phase cells with tripartite genome configuration. *^(13,30)^* Additionally, it was shown that segregation of the replicons follows a predominant order, with segregation of the *ori* of the main chromosome preceding those of the secondary replicons. *^(13)^* This raises the question whether replicon fusions impair the spatial organization and spatiotemporal segregation of genomic DNA.

In this study, we constructed *S. meliloti* strains with a bi-and monopartite genome configuration, either with the genomic DNA molecules including one or more origins of replication. We show that a cointegrate of the three genomic replicons that per design preserved typical features of bacterial genome organization, maintained growth properties and key features of spatial organization and segregation dynamics of each replicon under standard laboratory culture conditions. Furthermore, we show that an engineered strain with monopartite genome configuration, harboring *oriC* as sole origin of replication and characterized by a strongly unbalanced replichore ratio is viable, but impaired in growth. Suppressor mutants of this strain with a missense mutation in the cell cycle histidine kinase CckA or a partial genomic inversion showed significantly restored growth, making them chassis candidates for facilitated integration of ECRs and for future construction of hybrid genomes.

## RESULTS AND DISCUSSION

### Replicon fusions preserving strand asymmetry patterns

To study the properties of a triple-replication origin chromosome in *S. meliloti* in terms of spatial organization and spatiotemporal dynamics as well as propagation stability during the cell cycle, we merged the tripartite genome of this α-proteobacterium into a single DNA molecule. Targeted fusions of the three replicons were achieved by Cre/*lox*-mediated recombination in *S. meliloti* SmCreΔhsdR. *^(31)^* This strain, further referred to as wild type in this study, lacks *hsdR* encoding a restriction endonuclease and carries chromosomally integrated *cre* encoding the Cre recombinase. This recombinase catalyzes site-specific recombination of DNA between *loxP* sites. Replicon cointegrations sporadically occurring in *S. meliloti* by recombination between *nodPQ1* and *nodPQ2* as well as between the *algI* paralogues *SMb20843* and *SMc01551 ^(21)^* acted as blueprint for selection of the fusion sites, as these sites were compatible with a nearly symmetric structure of the fusion construct with respect to the distribution of three origins of replication.

Following integration of *lox* sites, in close proximity to *nodPQ1* and *nodPQ2*, megaplasmid pSymA and chromid pSymB were initially merged to generate pSymAB by induced Cre/*lox*-mediated site-specific recombination. Thus, the resulting strain SmAB harbored a bipartite genome composed of the main chromosome and pSymAB (Fig 1A, S1 Fig).

**Figure 1.**
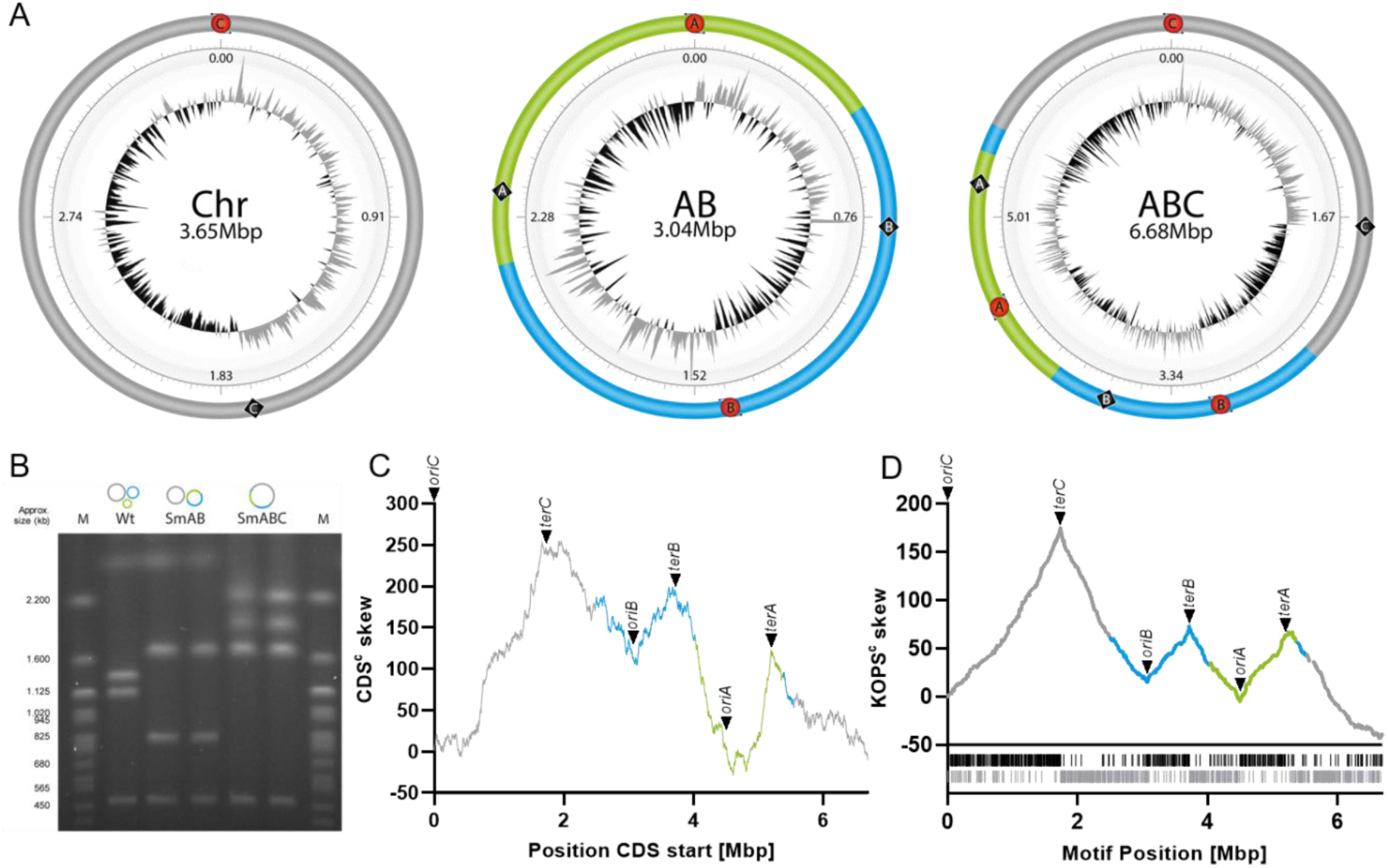
Design and compositional properties of the genomic content in *S. meliloti* replicon fusion strains SmAB and SmABC. (A) Configurations of replicon fusions. The inner concentric ring represents the deviation (positive: grey colored, negative: black colored) from the average GC-skew of the entire sequence calculated by a sliding window of (G-C)/(G+C). Replication origin of pSymA, pSymB and the chromosome (red circles including A/B/C); predicted terminus region of pSymA, pSymB and the chromosome (black diamonds including A/B/C); chromosome (Chr); pSymAB (AB); fusion product of pSymAB and the chromosome (ABC). **(B)** Pulsed-field gel electrophoresis of PacI digested gDNA from *S. meliloti* strains SmCreΔhsdR (Wt), SmAB (AB) and SmABC (ABC). Expected banding pattern for SmCreΔhsdR (wt): 3.65 Mbp (chromosome), 1.35 Mbp (pSymA), 1.15 Mbp (pSymB fragment 1), 0.53 Mbp (pSymB fragment 2). SmAB: 3.65 Mbp (chromosome), 1.67 Mbp, 0.83 Mbp, 0.53 Mbp. SmABC: 2.53 Mbp, 1.94 Mbp, 1.67 Mbp, 0.53 Mbp. M: PFGE marker yeast chromosomes, *Saccharomyces cerevisiae* (Strain YNN295). **(C)** Cumulated CDS skew (CDS^c^ skew) curve represents the coding sequence orientation bias in the SmABC fusion strain. **(D)** KOPS distribution in SmABC. For the analysis and representation of the replicon structure the *E. coli* KOPS consensus sequence (GGGNAGGG) was used. KOPS^c^ visualizes the accumulation of the KOPS motifs either located on the forward strand (black lines) or on the reverse strand (grey lines) of the sequence in SmABC. Color code: chromosome (grey), pSymA (green), pSymB (blue).

Following integration of a *lox* site in vicinity to both *algI* paralogues on chromosome and pSymAB, and induced Cre/*lox*-mediated site-specific recombination, these genomic elements were merged, giving rise to a single genomic DNA molecule in strain SmABC (Fig 1A, S1 Fig). To reduce the possibility of direct revertants after the A-B and AB-C fusion steps, we locked the genomic design by deletion of the remaining active *lox* site on both the AB and ABC fused replicons (S1 Fig). Thus, in SmAB and SmABC, the AB and ABC DNA molecules each contained only one remaining inactive *lox* site with downstream antibiotic resistance marker. Fusion sites, correct genomic arrangements, and genome sequence of SmAB and SmABC were validated using PCR, Pulsed-field gel electrophoresis (PFGE) (Fig 1B), and Illumina paired-end sequencing, respectively.

SmAB and SmABC genomic DNA configurations were designed to display wild type-like replichore ratios and distribution of GC content (Fig 1A, S2 Fig). Replichore orientation was reported to be defined by factors such as KOPS (FtsK orienting polar sequence) motifs and orientation of strongly expressed genes, both mostly situated on the leading DNA strand. *^(32–34)^* Each of the *S. meliloti* replichores contains a higher number of protein-coding genes on the leading than on the lagging strand (S3 Fig). This bias was preserved in SmAB and SmABC, even though two and three of the six replichores, respectively, were composed of segments originating from two or three different replicons of the wild type (Fig 1C). Also, the distribution of sequence motifs matching the *E. coli* KOPS consensus reflected the replicon structure and showed a strand asymmetry pattern with increasing motif density from *ori* towards *ter* in the replicons of wild type, SmAB and SmABC (Fig 1D, S4 Fig).

### Cell growth and morphology were not affected by the genome rearrangements

Like other α-proteobacteria, *S. meliloti* undergoes asymmetric cell division which leads to siblings of uneven cell size. *^(13,35,36)^* Cell shape and size of the *S. meliloti* replicon fusion strains and the wild type were indistinguishable by microscopic analysis (Fig 2A). A few cells with strongly asymmetrically placed constriction sites, probably resulting in minicells after cell division, were observed in cultures of the *S. meliloti* wild type, SmAB and SmABC strains in TY (complex) and high salt TY media (S5A Fig). A quantitative analysis identified a maximum of 0.3 % minicells in wild type and replicon fusion strain cultures (S5B Fig, S5C Fig).

**Figure 2.**
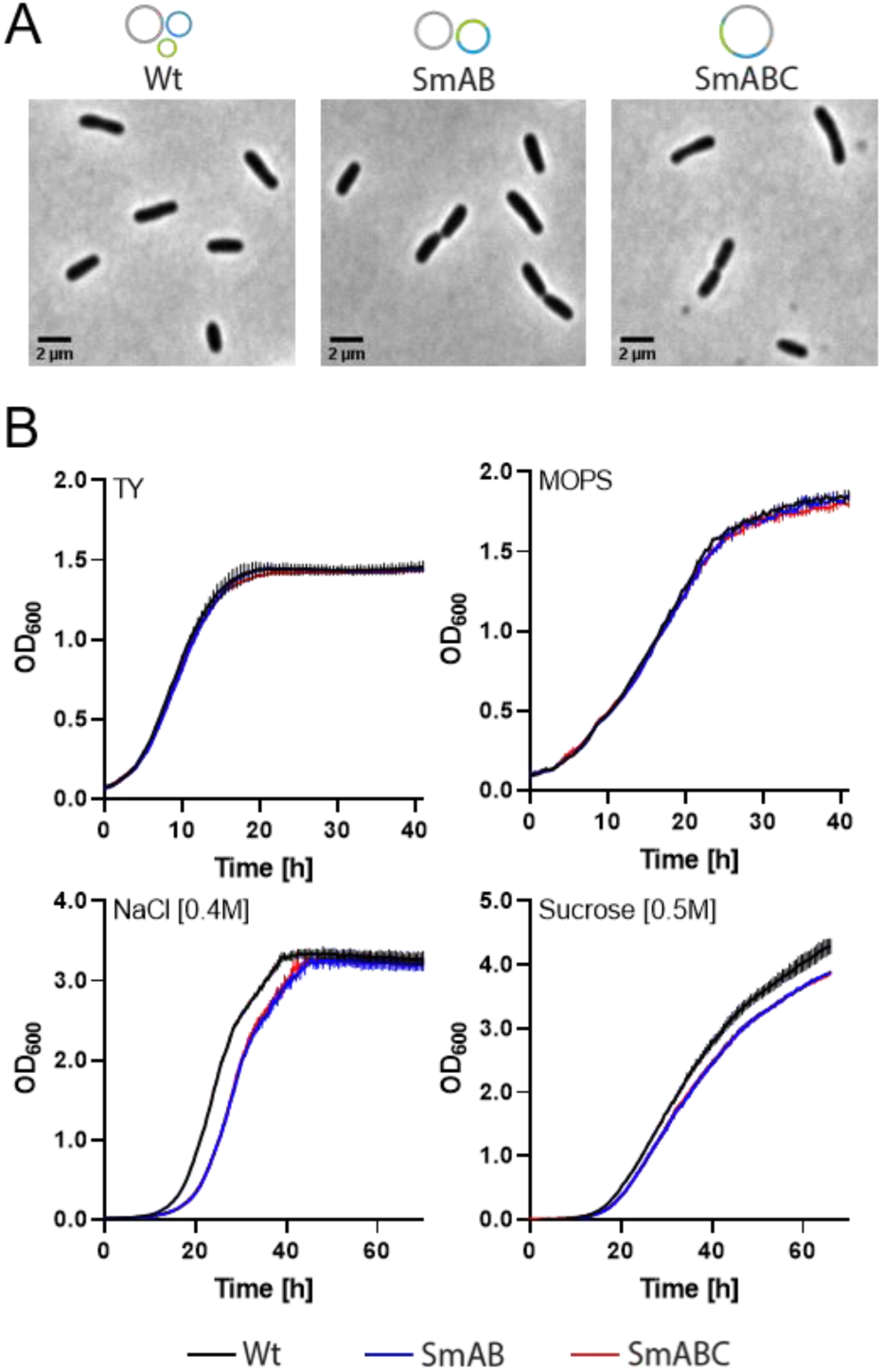
Basic characterization of *S. meliloti* replicon fusion strains SmAB and SmABC. (A) Phase contrast microscopy images of SmAB, SmABC and precursor strain SmCreΔhsdR (wt) representative cells at different stages of the cell cycle. Scale bar: 2 µm. **(B)** Growth of *S. meliloti* SmAB and SmABC compared to precursor strain SmCreΔhsdR (wt) in rich medium (TY), minimal medium (MOPS low Pi), high salt medium (TY + 0.4 M NaCl) and high sucrose medium (TY + 0.5 M sucrose). Data represent the mean ± standard deviation of three technical replicates. Growth curves of biological replicates are shown in S6A Fig.

When cultivated in TY or low phosphate minimal MOPS (defined) media, cell growth of both replicon fusion strains was similar to that of the wild type (Fig 2B) indicating that the new genome configuration does not significantly interfere with cell cycle processes. To analyze the effect of high-salinity and hyperosmotic stress on these strains, TY medium was supplemented with 0.4 M NaCl and 0.5 M sucrose, respectively. Compared to the wild type, SmAB and SmABC appeared slightly impaired in growth in these media (Fig 2B). Supplementing TY medium with 0.6 M NaCl or 0.7 M sucrose enhanced the difference in growth between wild type and replicon fusion strains (S6B Fig). High salt conditions are known to influence the supercoiling state of genomic DNA, *^(37)^* thus, the triple replicon fusion strain may be less robust to conditions affecting DNA condensation.

### The secondary replicon *oriV*s were dispensable in the strain with monopartite genome configuration

We asked if a single replication origin would be sufficient for proper vertical transmission of the AB and ABC DNA molecules in SmAB or SmABC. To this end, the *repB*-*repC* intergenic region and the *repC* coding region (*repC2* of pSymA and *repC1* of pSymB) were targeted for deletion (S7A Fig). In the wild type, deletion of these *repC* regions was not achieved. However, SmAB or SmABC lacking either of these regions (SmABΔrepC1, SmABΔrepC2 or SmABCΔrepC1, SmABCΔrepC2) or SmABC lacking both these regions (SmABCΔoriV) were obtained (S7B Fig). The configuration of the genomic DNA in all these strains was validated by PFGE (S7C Fig). Illumina sequencing verified the intended genetic changes in SmAB, SmABC and SmABCΔoriV, and revealed the modifications resulting from the construction process (S4 Table, S5 Table). Apart from these modifications, we identified only up to three single nucleotide variations (SNVs) per engineered genome compared to the wild type genome (S6 Table). Considering the nature of these changes, we suppose they are unlikely to be relevant to bacterial strain survival or fitness.

The sequential inactivation of the secondary replicon *oriV*s was associated with a gradually increasing growth defect, which appeared to correlate with the number of *oriV*s per DNA molecule and are possibly due to replication obstacles arising from the new replichore structures. Deletion of a single *repC* copy in SmAB resulted in a more severe growth defect than in SmABC, whose monopartite genomic DNA still carried two *ori*s (*repC* and *oriC*) after deletion of one of the *repCs*. However, deletion of both *repC* copies in SmABC caused the most severe growth defect (Fig 3A), even though cell morphology was inconspicuous in a microscopic snap shot image analysis (Fig 3B).

**Figure 3.**
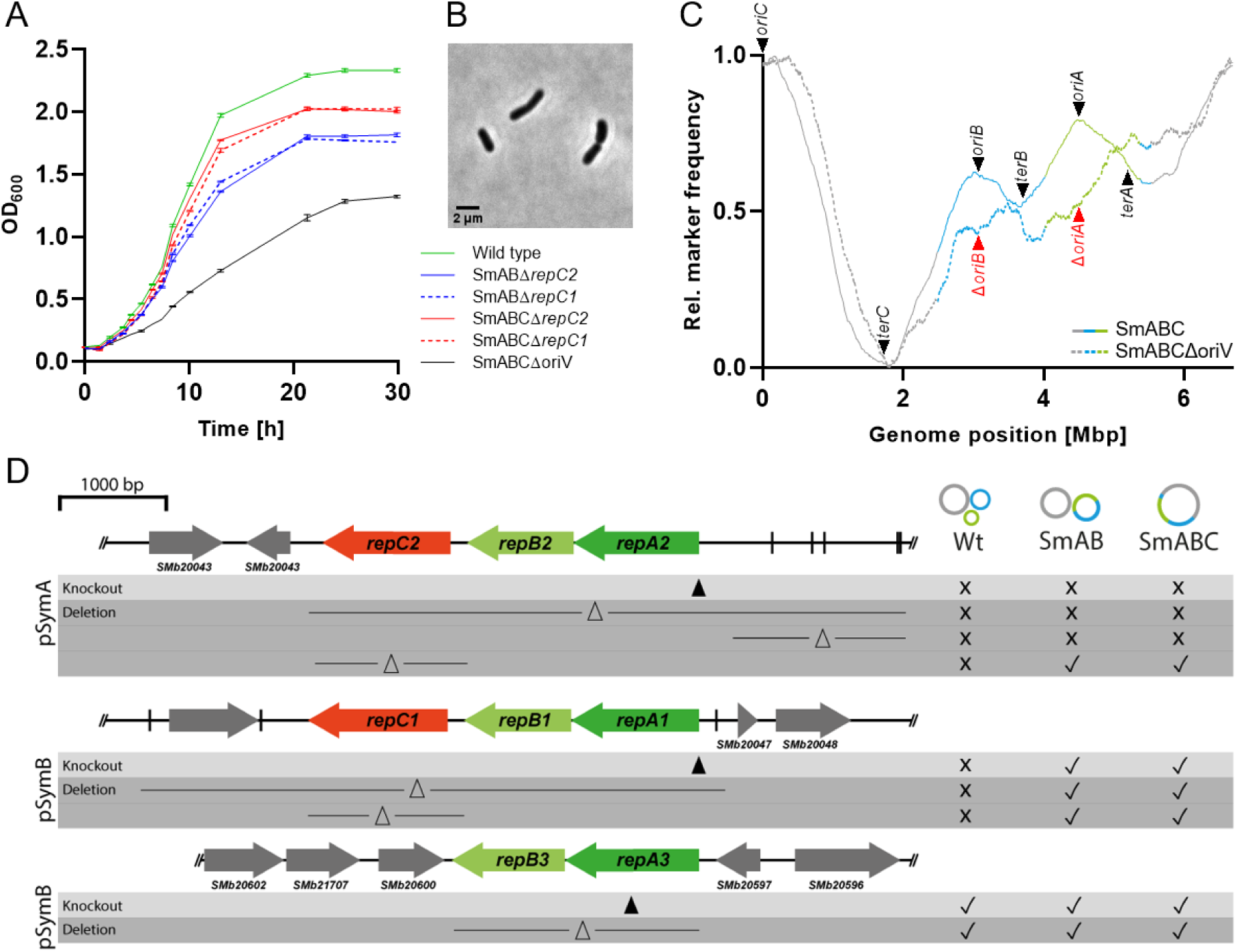
**Deletion and mutational studies in *S. meliloti* SmCreΔhsdR (wt), SmAB and SmABC fusion strains**. **(A)** Growth curves of *S. meliloti* SmAB and SmABC *repC* deletion strains. Prior to inoculation, overnight cultures were washed with 0.9 % NaCl and adjusted to an OD600 ∼ 0.15 in TY medium. Mean and standard deviation was calculated from three technical replicates. **(B)** Morphology of the double *repC* deletion mutant strain SmABCΔoriV. **(C)** Marker frequency analysis of SmABC and SmABCΔoriV of logarithmic (OD600 of 0.6) vs. stationary (OD600 of 2.6) cultures. Trimmed and normalized marker frequencies are depicted in log2 as a function of the genome position in Mbp. Arrowheads indicate the position of origin and predicted terminus regions. *^(38)^* Note that SmABCΔoriV lacks *oriA* and *oriB* due to *repC* deletions (Δ*oriA* and Δ*oriB*). **(D)** True to scale scheme of the secondary replicon derived replication and partitioning loci in *S. meliloti*. Short lines perpendicular to the sequence line depict the position of palindromic sequences (*parS*) involved in the process of DNA partitioning. Black arrowheads point to the area of gene disruption. The lines with a capital delta in between represent the regions that were targeted for deletion.

To test if replication is initiated at all three *ori*s and to determine the replication termination regions of the triple replicon fusion molecule in SmABC a Marker Frequency Analysis (MFA) was performed. The MFA data indicate that all three *ori*s mediated bidirectional replication (Fig 3C, S8B Fig). Furthermore, this data suggests that replication terminated in regions close to or overlapping the MFA minima for chromosome, pSymA and pSymB identified in the parental strain with tripartite genome configuration (S8A Fig), which roughly match with predicted *ter* regions. *^(38)^* MFA analysis of SmABCΔoriV shows clear sequence read enrichment with a maximum at *oriC* but no additional enrichment maxima at the mutated *repABΔC* (Δ*oriA*/Δ*oriB*) regions. The latter suggests loss of function of *oriA* and *oriB* in SmABCΔoriV (Fig 3C). This analysis also suggests replication termination in a region close to or overlapping *terC*, which separates two replichores largely differing in size with a ratio of approximately 3:1 (S8C Fig). The conservation of *terC* in the single replicon strain SmABCΔoriV, with even extremely skewed replichores, provides evidence for mechanisms defining this region for termination in the replication process. Differences in replication fork progression are maintained, e.g., by replication fork trapping sequences that slow down or stall the replisome. *^(39)^* In contrast, blocking of replications fork progression was not found for the terminus regions of either of the two chromosomes of *V. cholerae*. *^(40)^*

Plateaus or decrease in locus frequencies, including Δ*oriB*, *terB* and *terA* regions, might indicate replication obstacles in the single-replicon monopartite genomic DNA. Nevertheless, replisomes that had initiated replication at *oriC* do not appear to have been ultimately stalled in the original pSymA and pSymB terminus regions.

In addition, fluorescence microscopy-based analysis of mCherry-fused DNA polymerase III beta subunit DnaN revealed a maximum of two fluorescent foci in SmABCΔoriV (only *oriC* present) in constrast to four to five foci observed in parallel over the cell cycle in SmABC (*oriC* and both *oriV*s present) (S9 Fig). This is indicative of a reduced number of replisome formations in SmABCΔoriV and further confirms inactivation of both *oriV*s in this strain. Collectively, MFA and this microscopy data indicate that SmABCΔoriV contains a single 6.7 Mbp replicon and that the initiator protein-encoding *repC* genes including the *oriVs* of the secondary replicons are dispensable in this strain.

### The partitioning system of pSymB was dispensable in the replicon fusion strains

It is thought that the RepAB proteins are specific to partitioning of the cognate *repABC*-type replicons. *^(41)^* To study if the gene products of the *repA* and *repB* genes play a role for partitioning of the AB and ABC fusion replicons, we aimed to delete or inactivate the *repAB* gene regions individually and together in SmAB and SmABC (Fig. 3D). pSymA contains the single *rep* locus *repA2B2C2*, whereas pSymB harbors the complete *repA1B1C1* and the truncated *repA3B3* locus.

We found striking differences regarding the necessity of these elements in wild type, SmAB and SmABC strains. Deletion of *repA1B1C1* with adjacent partitioning sites and of *repA3B3* was possible in SmAB and SmABC (Fig. 3D). In the wild type, deletion or inactivation was achieved for *repA3B3* only (Fig.3D). Also, we obtained double mutants by deletion of *repA1B1C1* and inactivation of *repA3B3* in SmAB and SmABC but not in the wild type. Thus, the pSymB partitioning system appears to be dispensable for *S. meliloti* harbouring AB or ABC replicon fusions, in contrast to the results reported for *Agrobacterium tumefaciens*. *^(23)^* Deletion of pSymA-associated *repA2B2C2* with adjacent partitioning sites and of the partitioning sites alone was not successful in wild type and both replicon fusion strains (Fig. 3D). We also did not achieve inactivation of this *rep* locus by plasmid integrations in these strains. This suggests that the pSymA partitioning genes are required in *S. meliloti*, even if pSymA and pSymB or all three replicons are fused. An important role for the pSymA-derived partitioning system-encoding genes in the triple replicon fusion strain was unexpected since *S. meliloti* can be cured from pSymA. *^(17,42)^*

### A triple color fluorescent labeling system for simultaneous microscopic monitoring of *oriC* and two further freely selectable genomic loci

To enable live cell imaging analyses of spatial organization and spatiotemporal dynamics of the rearranged genomic DNA in the replicon fusion strains SmAB, SmABC and SmABCΔoriV, we established a triple color fluorescent labeling system. As this required using several antibiotic resistance markers, we eliminated by homologous recombination the inactive *lox* sites together with the downstream resistance markers that remained from the replicon fusion procedure in the AB and ABC DNA molecules (S1 Fig, S1A Data). This gave rise to derivatives of SmAB, SmABC and SmABCΔoriV termed SmABΔR, SmABCΔR and SmABCΔoriVΔR, respectively. The genome architecture of these strains was validated by PFGE (S1B Data) and Illumina genome sequencing.

ParB is known to specifically recognize and bind cognate *parS* sequences typically located around the chromosomal origin of replication in various bacterial species. *^(43)^* This protein was previously reported to be essential in *S. meliloti ^(44)^* and used as marker for *oriC* in this bacterium. *^(13,45)^* Indeed, we found good evidence for *Caulobacter crescentus-*like *parS* sites localized close to *oriC* in *S. meliloti* (S10C Fig). In our study, we employed a ParB-cerulean fusion to label *oriC*. To this end, *parB* at its native chromosomal locus was replaced by a *parB*-*cerulean* fusion in SmABΔR, SmABCΔR, SmABCΔoriVΔR, and the wild type (S10A Fig). Growth of the parental strains and corresponding derivative strains was indistinguishable in TY medium (S10B Fig), suggesting that the ParB-cerulean protein was functional.

In addition, a fluorescence repressor operator system (FROS) for labeling of two freely selectable genomic loci was established. This system combined *tetO* and *lacO* operator arrays *^(46)^* with *tetR-mVenus* and *lacI-mCherry* placed under control of the P*_tauA_* promoter on the replicative and mobilizable low copy vector pFROS (S10A Fig). In this setup, the basal activity of P*_tauA_ ^(47)^* was sufficient to generate a well detectable fluorescent signal over analysis periods of at least up to 6 hours in our study (S10D Fig), which makes the system particularly suitable for time lapse applications.

### Polar localization of *oriC* and subpolar localization of both *oriV*s at the old cell pole, as well as positioning of *terC* at the new cell pole was conserved in SmABC

We asked for the effect of replicon fusions on the spatial organization of the genomic DNA in *S. meliloti* by comparing monopartite and tripartite genome architectures. To this end, SmABCΔR and wild type derivatives, both carrying the *parB-cerulean* marker, were equipped with *tetO* and *lacO* arrays for simultaneous labeling of *oriC*/*oriA*/*oriB*, *oriC/terC*, *oriC*/*oriA*/*terA* and *oriC*/ori*B*/*terB* (Fig 4A, S11A Fig). In addition, we constructed derivatives of both strains for simultaneous labeling of *oriC* via ParB-cerulean and one of 12 further genomic loci either by integration of a *tetO* or *lacO* array. The labeled positions included genomic loci adjacent to the sites used for fusion of the replicons (Fig 4A, S11A Fig). Prior to fluorescence microscopy analysis, these strains were validated by PFGE for their correct genome configuration (S1C Data, S1D Data), and pFROS was introduced to mediate fluorescent labeling of the genomically integrated *tetO* and *lacO* arrays. For subcellular 2D mapping of the labeled genomic loci by snap-shot imaging of live cells, we filtered cells by size with a maximum cell length of 2.0 µm and displaying a single ParB-Cerulean-mediated fluorescent focus in one of the two cell pole regions only (Fig 4B). This configuration is indicative of a G_1_-phase cell with *oriC* localized at the old cell pole. *^(13,30)^* On average, the cell lengths were 1.8 ± 0.1 µm for both wild type and SmABCΔR.

**Figure 4.**
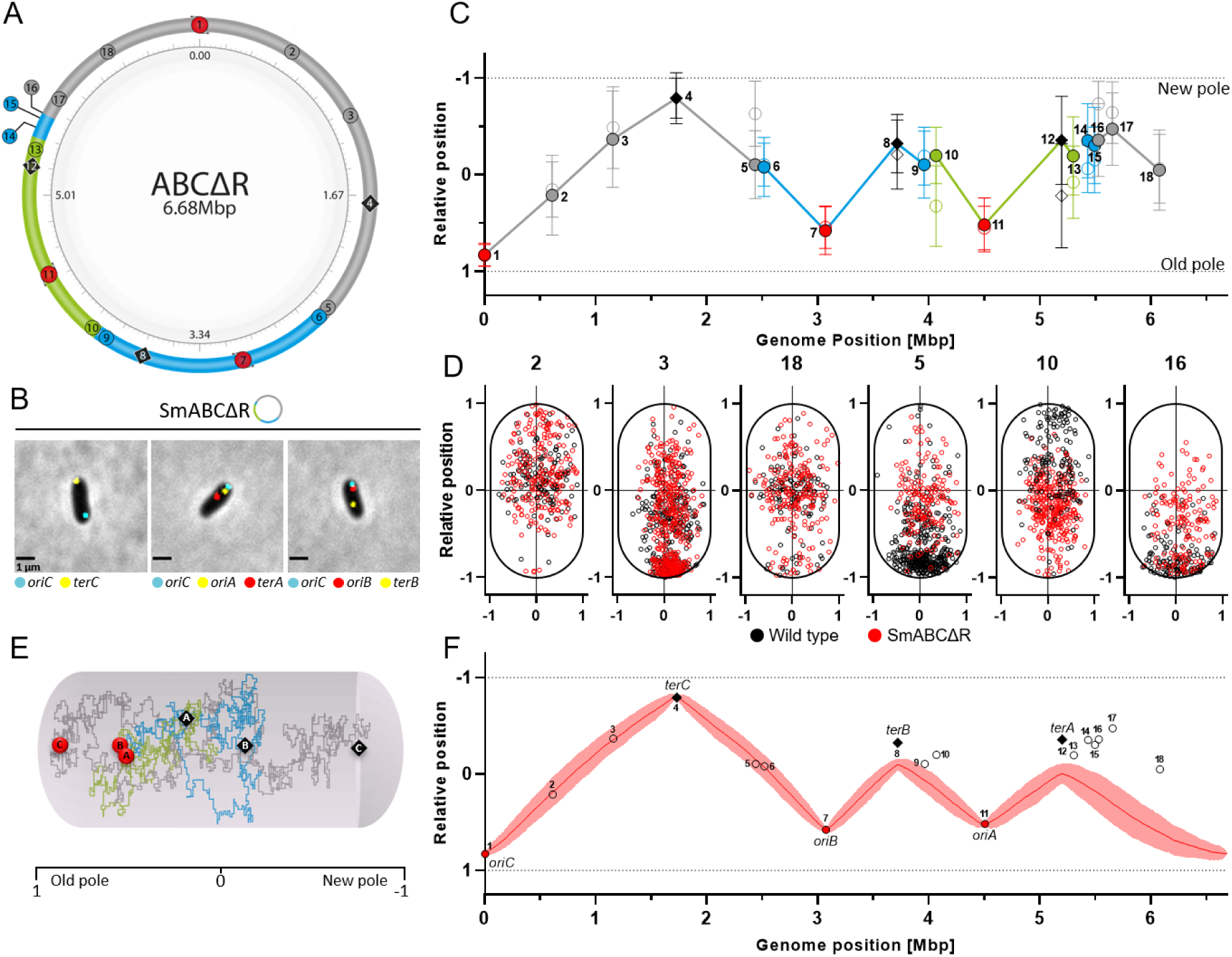
Investigation and modeling of the spatial DNA organization in SmABCΔR. (A) Schematic true to scale representation of the SmABCΔR monopartite genome with *tetO* and *lacO* integration sites 1-18 selected to reveal the spatial configuration of the genomic DNA. Color code: chromosome (grey), pSymA (green), pSymB (blue). Red circles: replication origins, black diamonds: terminus regions. **(B)** Example of snapshot images from labeled cells used for the 2D genome mapping study. Scale bar: 1 µm. **(C)** Normalized spatial localization of labeled loci within SmABCΔR (filled circles) compared to SmCreΔhsdR (non-filled circles) with wild type genome configuration as a function of the genome sequence coordinate. Old cell pole: 1, New cell pole: −1. **(D)** Scatter plots of selected strains illustrate examples of similar (Pos. no.: 2, 3, 18) and clearly different distribution (Pos. no.: 5, 10, 16) of marked loci within SmABCΔR (red dots) and SmCreΔhsdR (black dots). **(E)** Example configuration of simulations consolidating physical principles such as self-avoidance and compaction of DNA in a SmABCΔR cell represented as sphero-cylindrical shape. **(F)** Model of DNA self-organization in SmABCΔR compared to the experimental data. The model considers a spatial confinement (experimental standard deviation) for *terA* and *terB* in addition to *oriABC* and *terC* as fixpoints. Shown are the normalized locations in the cell as a function of the position on the genomic map. Experimental data of origins are indicated by red circles, terminus regions by black diamonds and remaining marker positions with non-filled circles. The red line depicts the model results averaged over 276 cells. Shaded areas represent the standard deviations. For the model a cell of 1800 nm and a loop-size of 1298 bp (DNA within a “blob”) was used.

Plotting the subcellular location of individual markers as function of the genomic position showed a tripartite triangle pattern, indicating arrangement of DNA segments between *ori* and *ter* along the longitudinal cell axis (Fig 4C). This is in agreement with a recent study also suggesting a longitudinal *ori-ter* distribution of a natural cointegrate of the circular and linear chromosomes in *A. tumefaciens*. *^(23)^* In wild type and SmABCΔR, *oriC* and *oriA*/*oriB* occupied polar and subpolar areas, respectively (S11B Fig). As expected from our filtering approach and in agreement with previous reports, *^(13,30)^* in both strains, the *oriC* signal showed a very low positional variance at one cell pole (S1 Table). Subpolar localization of *oriA* and *oriB* signals with similar variances in both strains suggests a spatial confinement of these elements to this region of the cell (S1 Table). The *terC* fluorescence signal was enriched with low positional variance at the cell pole opposite to the pole exhibiting the ParB-cerulean signal in wild type and SmABCΔR (S11B Fig, S1 Table). In contrast, the average spatial positions of *terA* and *terB* signals differed between these strains (S11B Fig, S1 Table). Especially, the spatial shift of *terA* in the triple replicon fusion strain argues against active positioning.

Consistent with tethering of *oriC* and *terC* to opposite cell poles, the average subcellular signal positions of chromosomal markers 2, 3 and 18, which map in close vicinity to *oriC* or *terC*, did not much differ between both strains (Fig 4D, S1 Table). Changes in the average spatial positions were found for the fluorescent signals of markers 5/6, 9/10 and 15/16, which flank the replicon fusions sites in SmABCΔR compared to the wild type (Fig 4C, S1 Table). This was expected since the markers of each pair are situated on different DNA molecules in the wild type, whereas they were brought into close proximity by the replicon fusions.

Tethering of *ori* and *ter* regions to factors localized at cell poles was previously reported for several bacteria, including the α-proteobacteria *C. crescentus* and *A. tumefaciens*. *^(48)^* For both species, there is evidence that polar tethering of *oriC* is mediated by binding of a *parS*-bound cluster of the DNA-binding CTPase ParB to a cell pole-associated protein scaffold. *^(49^*^-*52*^) In *C. crescentus*, tethering of *terC* to the new cell pole is mediated by the terminus recognition protein ZapT, the Z-ring associated proteins ZapA and ZauP that colocalize with FtsZ at the new cell pole. *^(53^*^-*55*^) It can be assumed that orthologous proteins and similar mechanisms are also involved in the spatial confinement of *oriC* and *terC* at the old and new cell poles, respectively, in *S. meliloti*. However, factors responsible for the spatial confinement of *oriA* and *oriB* to the subpolar region of the old pole are still enigmatic. Factors confining the three *ori*s and *terC* to cell poles are probably decisive for the spatial organization of the merged three replicons.

To test the hypothesis that the spatial organization of DNA in SmABCΔR is only determined by the positioning of locally confined *ori* and *ter* regions and the inherent features of the DNA as semi-flexible polymer of compacted units, we simulated the DNA arrangement for genome configurations differing in number and position of confined loci. We expanded a previously described model *^(56)^* by implementing not only one origin and one terminus as possible fixpoints, but three of each. By ergodic sampling over 200 configurations (Fig 4E) using the MOS-algorithm *^(57)^* the average genomic organization was obtained.

Initially, we generated a model including only *oriC* (S12A Fig) and *oriC*/*terC* (S12B Fig) as anchoring points of the 6.7 Mbp DNA molecule at the old and the new cell poles. We found that confining these two loci are not sufficient to describe the spatial arrangement of pSymA-and pSymB-derived DNA. By anchoring of all three origins to the experimentally determined average subcellular position, we gained a ternary model structure with highly variable DNA segments between the three points (S12C Fig). However, in this model structure, the DNA segment between *oriC* and *oriB* did not extend up to the opposite cell pole. We then confined *terC* to the new cell pole, and *oriC* to the polar, and *oriA* and *oriB* to the subpolar regions of the old cell pole (S12D Fig) since these loci showed the smallest positional variance in the experimental data (S12E Fig). With these four anchored points the generated model structure already reflected the experimentally determined positions of loci 2, 3 and 5 as well as locus 6 located on the chromosome and pSymB, respectively. However, it did not describe the experimentally determined average localization of *terA* and *terB*. Loci close to *terA* and *terB* showed a high positional variance in the experimental data (S12E Fig). By a spatial confinement of *terA* and *terB* within the experimental data variance we gained a tripartite triangle structure (Fig 4F), albeit this model did not reproduce the experimentally determined average localization of loci 13 to 18.

### Spatiotemporal dynamics of origin and terminus regions in replicon fusions strains

To study the effect of bi-and monopartite genome configurations and reduced number of replication initiation sites on DNA segregation, we analyzed the spatiotemporal dynamics of the *ori* and *ter* regions in SmABΔR, SmABCΔR and SmABCΔoriVΔR in comparison to the wild type. For this purpose, we followed the same labeling strategy as described above, and validated the strain’s genome configurations by PFGE (S1 Data). At the level of individual cells, microscopic snap-shot and time lapse data of combinations of labeled *oriC*/*oriA*/*oriB*, *oriC*/*terC*, *oriC*/*oriA*/*terA* and *oriC*/*oriB*/*terB* were generated.

#### The spatiotemporal choreography of origins and termini was similar in the replicon fusion strains and wild type

A previous study showed a clear spatiotemporal pattern of duplication and segregation of the three *ori*s in the *S. meliloti* wild type. *^(13)^* This was characterized by initial occurrence of two *oriC* copies at the old pole, indicative of the start of chromosome segregation (approximately in the first quarter of the cell cycle), and followed by translocation of one *oriC* copy to the new cell pole. The *oriV*s translocate from the subpolar region of the old cell pole to the midcell area, where, after *oriC* segregation is complete, duplication and subsequent segregation to the subpolar cell regions takes place (approximately in the second quarter of the cell cycle).

Initially, we analyzed the spatiotemporal dynamics of fluorescently labeled *ori* and *ter* loci in mother cells of *S. meliloti* wild type, SmABΔR and SmABCΔR. For each individual cell analyzed, cell cycle duration and cell size were normalized to 100 % (0/100 % – completion of cell division) and to 1 (0 – old cell pole, 1 – new cell pole), respectively, to facilitate comparative analyses. Regarding the spatiotemporal dynamics of *oriC* and the *oriV*s, we observed similar patterns for the three strains (Fig 5A/B, S2 Table, S2 Data), which are in agreement with the previous results for the wild type. *^(13)^*

**Figure 5.**
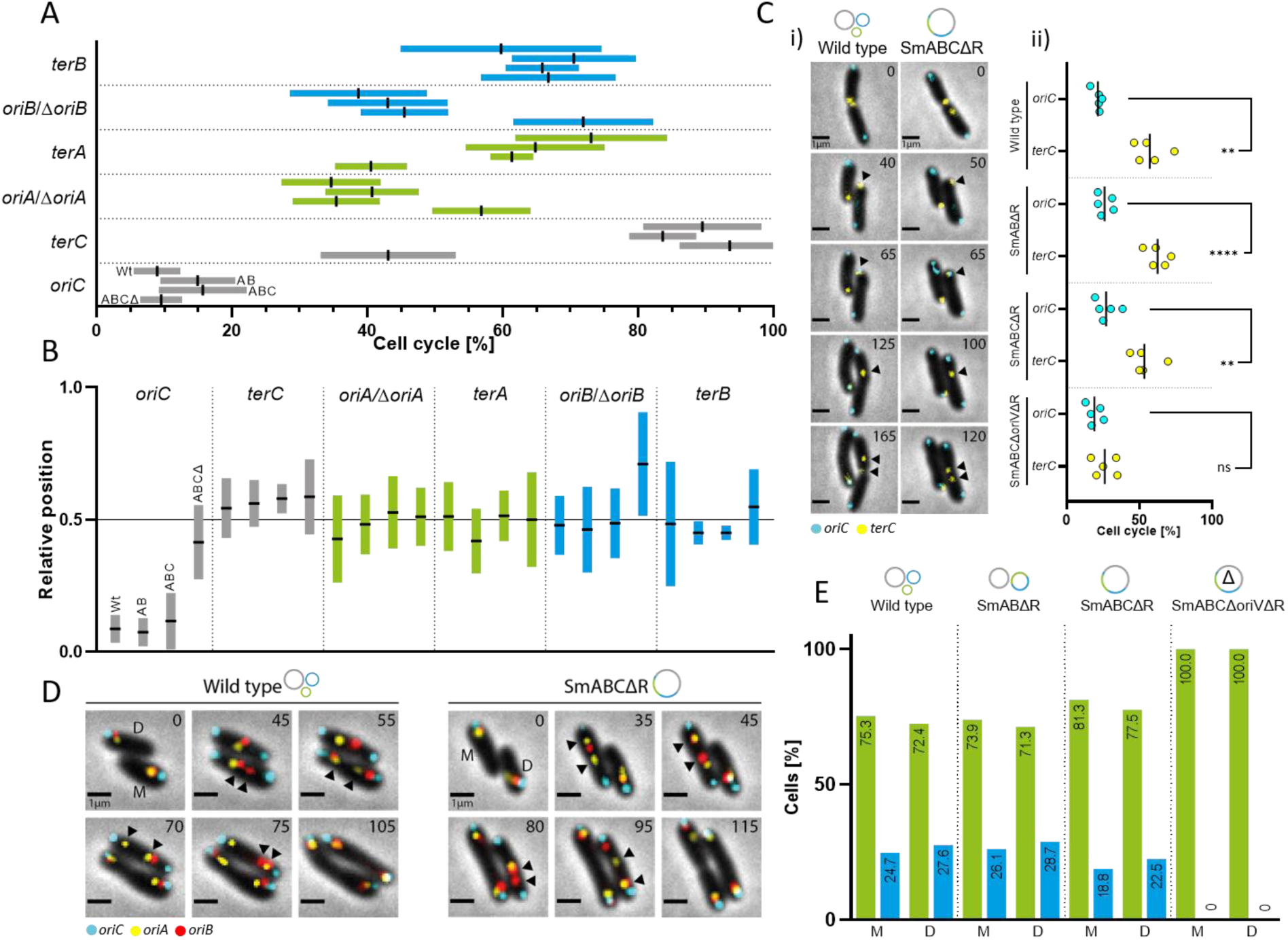
Spatiotemporal pattern of origin and terminus regions in native and reorganized multi-replicon backgrounds. (A) Temporal order of origin and terminus region segregation in SmCreΔhsdR (wt), SmABΔR (AB), SmABCΔR (ABC) and SmABCΔoriVΔR (ABCΔ). Colored bars with a black center indicate the standard deviation and the mean timepoint of sister foci separation within the cell cycle normalized to 100 %. Analyzed M-cells: 20 (*oriC*), 10 (*oriA/B*) and 5 (*terA/B/C*). **(B)** Localization of *ori* and *ter* foci segregation in SmCreΔhsdR (wt), SmABΔR, SmABCΔR and SmABCΔoriVΔR. The bar chart depicts the relative longitudinal position of a single focus before separation within the normalized cell (old pole: 0, new pole: 1, midcell: 0,5). Data represent the mean ± the standard deviation for *oriC* (n=20), *terC* (n=5), *oriA* (n=10), *terA* (n=5), *oriB* (n=10) and *terB* (n=5) in M-cells. **(C)** Spatiotemporal choreography of *oriC* and *terC* in SmCreΔhsdR (wt), SmABΔR, SmABCΔR and SmABCΔoriVΔR. (i) Time lapse series with *oriC* (ParB-cerulean) and *terC* (TetR-mVenus) in SmCreΔhsdR (wt) and SmABCΔR. Scale bar: 1 µm. (ii) Timepoint of *oriC* (cyan circles) arriving at the new cell pole in relation to *terC* (yellow circles) release from the same pole within a normalized cell cycle (0-100 %). Analyzed M-cells: 5 each. Black bar indicates the data mean. **(D)** Time lapse microscopy series of SmCreΔhsdR (wt) and SmABCΔR as examples of sequential segregation of *oriA* and *oriB* with varying order. Arrowheads depict the position of segregation start. **(E)** Percentage of cells with *oriA* foci (green) and *oriB* foci (blue) segregating first. Cells analyzed: SmCreΔhsdR (wt) (n=172), SmABΔR (n=179), SmABCΔR (n=169), and SmABCΔoriVΔR (n=197). Color code: chromosome (grey), pSymA (green), pSymB (blue). Mother cell (M), daughter cell (D). Note that SmABCΔoriVΔR lacks *oriA* and *oriB* due to *repC* deletions. For simplicity, localization of *repABΔC* loci is denoted as *oriA* and *oriB* in panels A, B, C and E.

Translocation of the *terC* foci from the new pole to midcell started in the three strains approximately in the middle of the cell cycle (Fig 5C). Two *terC* foci were observed at midcell for the first time in the last quarter of the cell cycle before cell division (Fig 5B/C). *terA* and *terB* foci already localized in the midcell area before duplicated foci were observed in this region (Fig 5B). Doubling of *terA* and *terB* foci was observed prior to *terC* segregation in the third quarter of the cell cycle (Fig 5A). In the daughter cells, choreography of individual *ori* and *ter* foci was similar to that in the mother cells described above (S13 Fig, S3 Table).

The previous study indicated a preferential temporal order of *oriA* and *oriB* segregation. *^(13)^* We asked if this order is conserved in cells with fused replicons. Upon detailed examination, we found that, duplication of *oriA* and *oriB* foci was very close in time in both the mother and daughter cells of wild type, SmABΔR and SmABCΔR (S14A Fig). It was observed either simultaneously (S14B Fig) or sequentially (Fig 5D), with *oriA* starting segregation before *oriB* in the majority of cells. In up to a fifth of wild type, SmABΔR and SmABCΔR cells analyzed (n=200 each), we found the doubling of *oriA* and *oriB* foci within the same time frame of 5 minutes (S14C Fig). In the remaining proportion of cells that showed sequential segregation of these origins, *oriA* segregation preceded that of *oriB* in three-fourth of these cells (Fig 5E), indicating that the previously observed temporal order *^(13)^* of duplication and segregation first of *oriA* and then of *oriB* was less stringent both in wild type and the replicon fusion strains.

In conclusion, the time-lapse analyses of the replicon fusion strains SmABΔR and SmABCΔR indicate that their spatiotemporal choreography of the origin and terminus regions is similar to that of the wild type with a tripartite genome configuration. This suggests that regulation of replication initiation and the processes relevant to segregation have been conserved.

#### The chronology of segregation and spatial position of *oriC* and Δ*oriA*/*B* regions are altered in SmABCΔoriVΔR

Strain SmABCΔoriVΔR carries monopartite genomic DNA containing *oriC* but lacking both secondary replicon origins (*repA1B1ΔC1,* Δ*oriB* and *repA2B2ΔC2,* Δ*oriA*). We studied the segregation chronology of active *oriC*, inactive Δ*oriA/B*, and terminus regions in this strain to learn about the segregation properties of the 6.7 Mbp single replicon.

In an initial fluorescence microcopy snapshot series, we analyzed putative G_1_-phase SmABCΔoriVΔR cells filtered with a cut-off of 2.0 µm cell length and deduced the relative distance of *oriC* and both Δ*oriA/B* foci to the cell equator. Striking differences were found between *oriC* localization in SmABCΔoriVΔR compared to wild type, SmABΔR and SmABCΔR cells. In contrast to wild type, SmABΔR and SmABCΔR cells that showed *oriC* foci clustering predominantly in the cell pole area, a considerable proportion of SmABCΔoriVΔR cells showed *oriC* localization in the midcell area (Fig 6A, S15 Fig). We also observed that in SmABCΔoriVΔR, both Δ*oriA/B* foci localized more frequently in the midcell area than in the cell pole area (S15 Fig), and that the proportion of cells showing colocalization of both Δ*oriA/B* foci with *oriC* in the midcell area was considerably higher than in wild type, SmABΔR and SmABCΔR cells (Fig 6B, S16 Fig). Collectively, this data suggests aberrant localization of *oriC* in more than one third of SmABCΔoriVΔR G_1_-phase cells.

**Figure 6.**
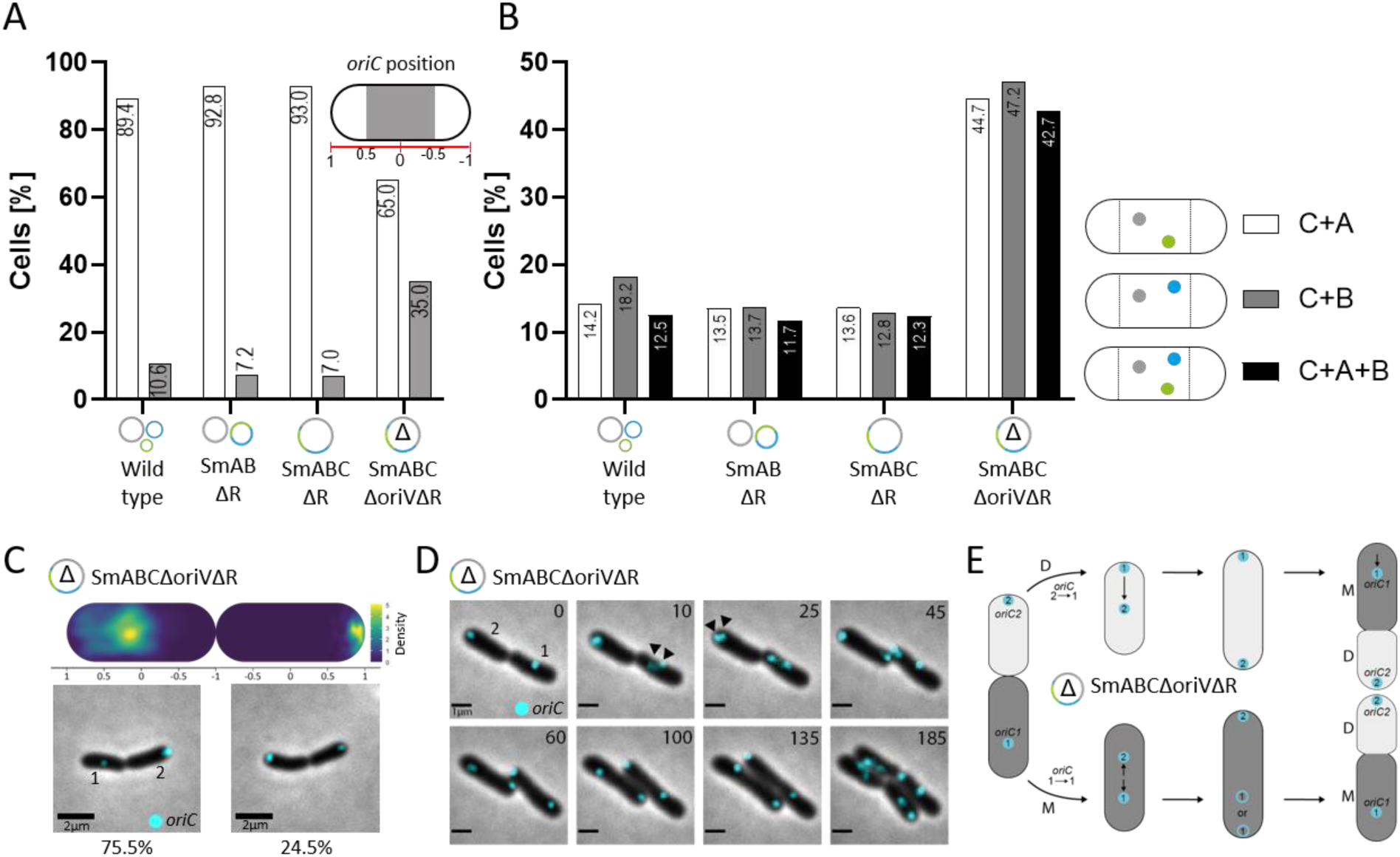
Mislocalization of the replication origin regions in SmABCΔoriVΔR. (A) Localization of *oriC* in SmCreΔhsdR (wt), SmABΔR, SmABCΔR and SmABCΔoriVΔR G1-phase cells. Bar chart depict the percentage of cells with *oriC* (ParB-cerulean) foci at the cell poles (white cell compartment, 0.5 to 1 and −0.5 to −1) and at midcell (grey cell compartment, 0.5 to −0.5). Cells analyzed: SmCreΔhsdR (wt) (n=480), SmABΔR (n=581), SmABCΔR (n=371), SmABCΔoriVΔR (n=936). **(B)** Co-localization of *oriA/B/C* in SmCreΔhsdR (wt), SmABΔR, SmABCΔR and SmABCΔoriVΔR G1-phase cells. Shown are the percentage of cells with localization of either *oriA/C* (white bar), *oriB/C* (grey bar) and *oriA/B/C* (black bar) in the midcell area (0.5 to −0.5). Cells analyzed: SmCreΔhsdR (wt) (n=639), SmABΔR (n=460), SmABCΔR (n=405), SmABCΔoriVΔR (n=246). Dots in the schematic cells represent *oriC* (grey), *oriA* (green) and *oriB* (blue). Note that SmABCΔoriVΔR lacks *oriA* and *oriB* due to *repC* deletions. For simplicity, localization of *repABΔC* loci is denoted as *oriA* and *oriB*. **(C)** Analysis of *oriC* localization in predivisional cells of SmABCΔoriVΔR. The density map indicates the position of the chromosomal origin in sibling 1 (left cell) and sibling 2 (right cell) of predivisional cells (n=400) in a normalized scale. Snapshot images and percentage of the two major *oriC* spatial position patterns in predivisional cells. Scale bar: 2 µm. **(D)** Time lapse microscopy series of generic *oriC* choreography in sibling cells of SmABCΔoriVΔR. Arrowheads depict the position of visible *oriC* foci segregation. Scale bar: 1µm. **(E)** Model of *oriC* choreography in SmABCΔoriVΔR. The model differentiates between *oriC* coordination in daughter (D, light grey) and mother (M, anthracite) cells. Cyan cycles with numbers (1: *oriC1*, 2: *oriC2*) represent the deduced *oriC* positioning within the cells.

To follow up on this observation, we performed time-lapse fluorescence microscopy of SmABCΔoriVΔR cells. Analysis of this data revealed two *oriC* localization patterns dominating in either the mother or the daughter cells (Fig 6C). These patterns are visualized in a schematic summary in Fig 6E. Pattern 1 was characterized by visible *oriC* foci segregation in the midcell area (Fig 5B, Fig 6D), which ultimately resulted in localization of one *oriC* focus (defined as *oriC2*) at the new cell pole and the other (defined as *oriC1*) in the mid-area of the mother cell compartment (S17 Fig). Pattern 2 was characterized by duplication of the *oriC* focus (*oriC1*) at the old cell pole followed by translocation of one of the *oriC* foci (*oriC2*) to the new pole (Fig 6D). Right before cell division, *oriC1* localized either at the old cell pole or in the mid-area of the mother cell compartment in 30% and 70% of the cells, respectively (S17 Fig). We identified the pattern 2 cells as the daughter cells since they most frequently adopted the *oriC* localization pattern of the ancestral cell (i.e. mother cell, pattern 1 cell) after one cell cycle. Moreover, we found that in mostly all mother cells analyzed displaying aberrant *oriC* localization also the Δ*oriA* and Δ*oriB* regions lost their subpolar localization (S2D Data). Notably, Δ*oriB* duplication in SmABCΔoriVΔR cells was observed in the new cell pole compartment and not in the midcell region as in wild type, SmABΔR and SmABCΔR cells (Fig 5B). Loss of polar *oriC* and subpolar Δ*oriA* and Δ*oriB* localization suggests that the deletion of *repC/oriV* directly or indirectly affect mechanisms relevant for anchoring of these sequence elements. Furthermore, the frequent colocalization of these regions (Fig 6B) even after loss of their original cellular position would be consistent with *ori*-*ori* clustering as described for the replicons in *A. tumefaciens*. *^(52)^*

In addition to the spatial alterations in segregation patterns in SmABCΔoriVΔR, we observed also alterations in temporal patterns associated to *ori* and *ter* regions. Translocation of the *terC* focus occurred already in the beginning of the second cell cycle quarter, and not shortly after the midpoint of the cell cycle (Fig 5Cii). This shift in timing was also reflected in earlier doubling of *terC* and also *terA* foci (Fig 5A). Moreover, duplicated foci representing the Δ*oriA* regions appeared later (halfway point of the cell cycle vs. second quarter of the cell cycle (Fig 5A). Thus, in SmABCΔoriVΔR cells, visible segregation of *terA* occurred before that of the Δ*oriA* region. Also the order of duplication of *terB* and Δ*oriB* foci was changed, with duplication of *terB* occurring on average before that of Δ*oriB* (Fig.5A). These alterations in chronology of foci duplication were observed in mother and in daughter cells (S13A Fig, S3A Table), and correlate with the position on the replicon sequence and the distance to *oriC*.

### A missense mutation in cell cycle kinase-encoding *cckA* increased fitness of SmABCΔoriVΔR, but is unlikely to be responsible for aberrant *ori* localizations

In addition to genome sequencing of SmAB, SmABC, and SmABCΔoriV (see above), we also determined the genome sequence of SmABΔR, SmABCΔR, and SmABCΔoriVΔR (S6 Table). In particular, a SNV in the coding sequence *SMc00471* of the cell cycle histidine kinase CckA in SmABCΔoriVΔR attracted our attention. This SNV that causes amino acid substitution R436H (S18A Fig, S6 Table) was not found in the precursor strains.

To test whether this missense mutation was responsible for the aberrant *oriC* localization pattern in strain SmABCΔoriVΔR we attempted to revert the SNV to the wild-type *cckA* sequence in this strain, and failed. We then introduced this missense mutation into the wild type and SmABCΔR and analyzed the localization of *oriC* right before cell division in G_1_-phase cells, using ParB-Cerlulean for fluorescent labeling (Fig 7A, S18B Fig, S18C Fig). Regarding polar (approx. 90 %) and midcell (approx. 10 %) localization of the ParB-Cerlulean mediated fluorescent focus, we found no difference between the strains carrying the SNV in *cckA* and the corresponding strains with wild type *cckA* (Fig 7A). However, in this comparison, growth of the strains with CckA^R436H^ was slightly reduced (S18D Fig).

**Figure 7.**
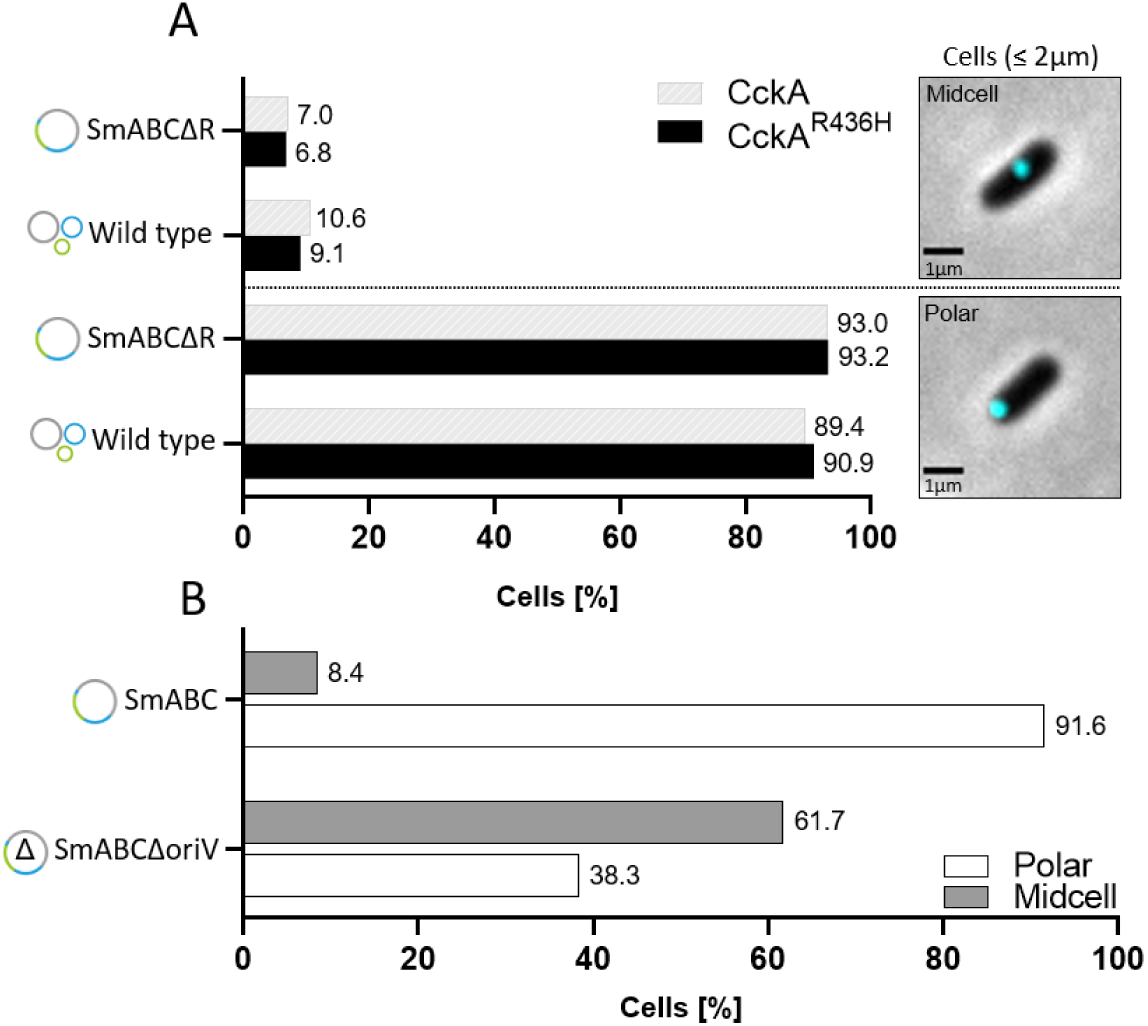
Localization of the chromosomal origin in fusion strains with wild type CckA and CckA^R436H^. (A) Localization of *oriC* (ParB-cerulean) in SmCreΔhsdR (wt) and SmABCΔR G1-phase cells with *cckAR436H* (black bars) compared to respective strains with wild type *cckA* (grey shaded bars). Cells analyzed: SmCreΔhsdR n= 480 (*cckAwt*), n=407 (*cckAR436H*), SmABCΔR n= 371 (*cckAwt*), n=352 (*cckAR436H*). Snapshot images as example for polar and midcell localization of *oriC*. Scale bar: 1 µm. **(B)** Comparison of polar and midcell localization of *oriC* in SmABCΔoriV (precursor of SmABCΔoriVΔR) and SmABC, both with wild type *cckA*. White bars depict the percentage of cells with polar *oriC* localization whereas grey bars represent midcell localization. Cells analyzed: SmABCΔoriV (n= 472), SmABC (n=584).

In addition, we used ParB-Cerlulean to analyze the *oriC* localization pattern in SmABCΔoriV, the direct precursor of SmABCΔoriVΔR. SmABCΔoriV carries the *cckA* wild type sequence and is already deleted for both secondary replicon replication origins (Δ*oriA*/*B*). We found that the *oriC* localization pattern in SmABCΔoriV already deviates from the wild type-like pattern in SmABC, which has three intact replication origins (Fig 7B). This implies that CckA^R436H^ did not cause the observed mislocalization of *oriC* in SmABCΔoriVΔR. However, we found a strong difference in growth between SmABCΔoriV (CckA) and SmABCΔoriVΔR (CckA^R436H^) (S18E Fig), indicating that the *cckA*_R436H_ allele mitigates the strong growth deficiency caused by deletion of the *oriV*s. In a growth comparison, SmABCΔoriVΔR (CckA^R436H^) reached ∼ 65 % of the growth rate of SmABC and the wild type in the exponential phase (Fig 8A).

**Figure 8.**
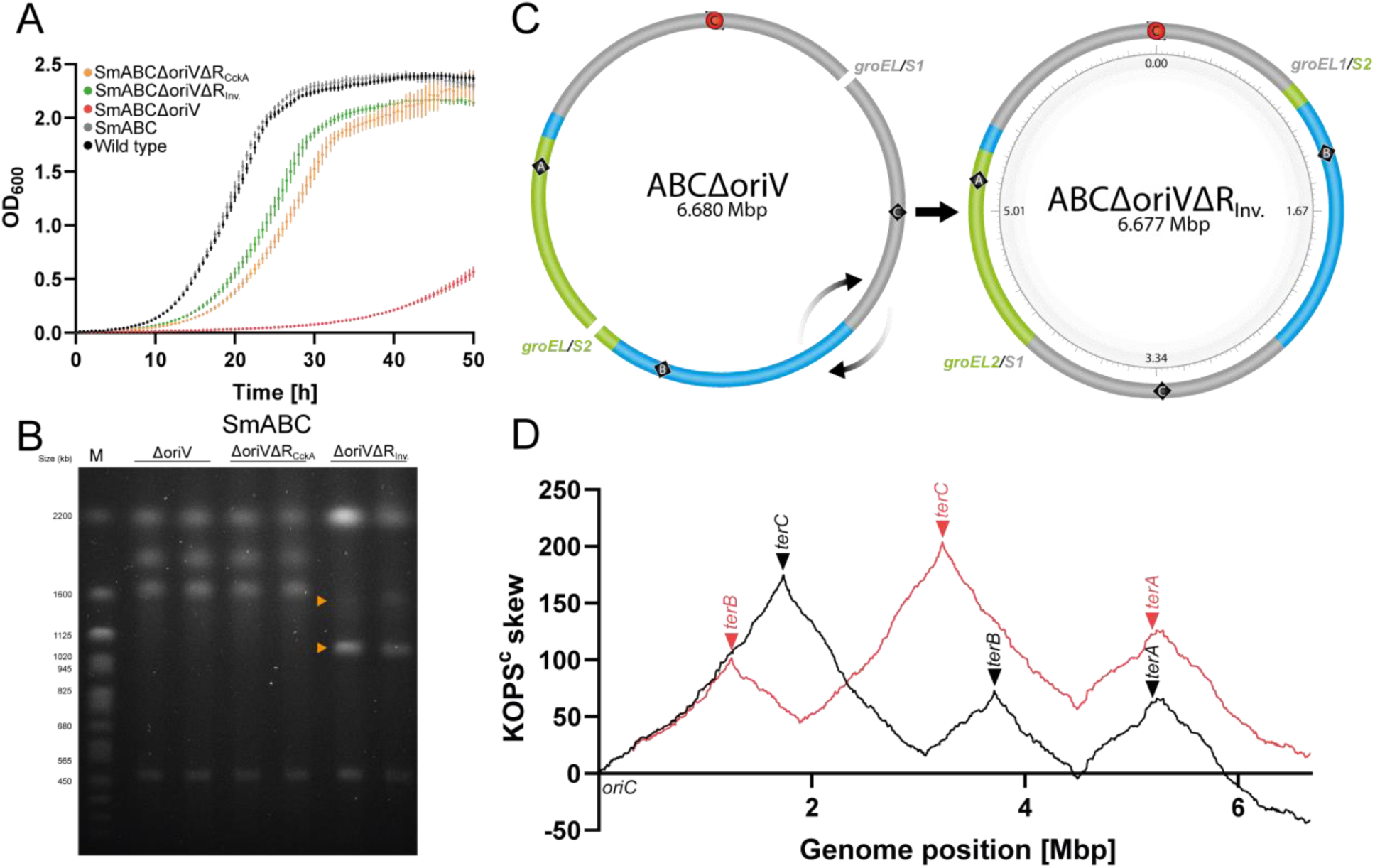
SmABCΔoriVΔR derivative with genomic rearrangement. (A) Growth of SmABCΔoriV (generation time: 7.2 h), SmABCΔoriVΔR (CckA^R436H^) (generation time: 3.7 h) and SmABCΔoriVΔRInv. (generation time: 3.2 h), compared to SmABC (generation time: 2.4 h) and the wild type (generation time: 2.5 h). The graph represents the mean values from 6 technical replicates at each time point. **(B)** Analysis of SmABCΔoriVΔRInv by PFGE revealed a distinct banding pattern (orange arrowheads) compared to SmABCΔoriVΔR with CckA^R436H^ and the precursor strain SmABCΔoriV. Expected fragment sizes in Mbp: 2.53, 1.94, 1.67, and 0.53. Size standard (M): *S. cerevisiae*, Biorad. **(C)** Schematic representation of genomic restructuring in SmABCΔoriVΔRInv. *De novo* genome identified a partial inversion between paralogous sequences *groEL*/*S1* (chromosome), and *groEL*/*S2* (pSymA). This partial inversion relocated the *terC* region opposite to the remaining *oriC* (174°), analogous to its previous position in the chromosome (170°). **(D)** Distribution of KOPS in SmABCΔoriVΔRInv. (red line) compared to the direct precursor strain SmABCΔoriV (black line).

Cell cycle regulation by the CckA-ChpT-CtrA signaling pathway is wide-spread in α-proteobacteria. *^(58)^* The CckA phosphorelay controls the phosphorylation status of ChpT which in turn regulates activity and stability of the cell cycle master regulator CtrA through phosphorylation. *^(59)^* CckA phosphatase activity enables initiation of DNA replication through dephosphorylation and degradation of CtrA. (*59*,*60*) The *S. meliloti* (S18A Fig) and *C. crescentus* CckA domain composition *^(61)^* is very similar. The R436H substitution in *S. meliloti* CckA locates in one of the PAS domains, which in *C. crescentus* were shown to regulate switching between the CckA kinase and phosphatase activities. *^(61,62)^* Narayanan and coworkers *^(63)^* identified a point mutation in the PAS-B domain of *C. crescentus* CckA that suppresses a topoisomerase IV inhibitor-induced chromosome segregation defect, possibly by slowing down the chromosome replication cycle. CckA^R436H^ is therefore proposed to mitigate chromosome replication and/or segregation defects in a similar manner. We speculate that the R436H substitution promotes kinase or reduces phosphatase activity of *S. meliloti* CckA, which results in higher levels of CtrA-P repressing replication initiation and slowing down the cell cycle.

#### A 3.2 Mbp genomic inversion relocalizes *terC* opposite to *oriC*, and alleviates the growth deficit of SmABCΔoriV

While establishing strain SmABCΔoriVΔR, we detected a further derivative, which showed ∼ 75 % of the growth rate of SmABC and the wild type (Fig. 8A). This clone showed altered banding patterns in PFGE analysis (Fig. 8B), suggestive of genomic rearrangement events. Combining nanopore and Illumina sequencing data, we *de novo* assembled the genome sequence of this strain, and detected a 3.2 Mbp large inversion. The inversion in this strain, hereafter called SmABCΔoriVΔR_Inv_, occurred between the *groEL1*/*S1* and *groEL2*/*S2* paralogs, which map to the chromosome and pSymA sequences, respectively. This inversion positioned the *terC region* opposite to *oriC* (Fig. 8C), and realigned KOPS across the single replicon (Fig. 8D). The CckA^R436H^ missense mutation characteristic for SmABCΔoriVΔR was not detectable in SmABCΔoriVΔR_Inv_ (S6 Tab.). Remarkably, the point mutation in *cckA* alleviated the growth deficiency of the ancestral strain almost as much as this massive genetic rearrangement (Fig. 8A).

Previous studies of engineered *E. coli* strains indicate that disturbing chromosome organization patterns by integration of one or more additional *oriC* copies or re-localization of *oriC* to ectopic locations can cause replication-transcription conflicts and issues with replication fork trap regions, which affect DNA replication and segregation, and promote selection of phenotype-moderating genome rearrangements and genetic suppressions. *^(64,65)^* In naturally occurring single-chromosome Vibrio (NSCV) strains of *V. cholerae*, a second origin was found to be either active or silenced depending on the position of the cointegration event, *^(66)^* whereas in laboratory-generated fusions the chromosome 2 replication machinery was not functional. *^(67,68)^* Recently, a derivative of the ultrafast growing *Vibrio natriegens* with fused chromosomes 1 and 2 was constructed, which is not impaired in growth under standard laboratory culture conditions. *^(69)^* The design of this strain preserved only the *ori* and *ter* region of chromosome 1 and maintained genomic symmetries. The results studies in different bacterial species and our study in *S. meliloti* suggest that retaining replichore symmetry and replichore-orienting sequence element distributions as well as avoiding replication termination traps are important design rules for the fitness of replicon fusion strains. But even strains that do not sufficiently fulfill these rules but are still viable can be evolvable into fitter strains.

## CONCLUSION

Members of the α-, β-, or γ-proteobacteria are specialists in occupying ecological niches, not least due to their genetic diversity, which is enhanced by segmented genome configurations. Because of their metabolic diversity, several of these bacteria are already used extensively in the fields of biotechnology, climate protection, plant genetics and sustainable agriculture and are thus of great value for the transformation of a future bioeconomy. Although there is a benefit from the segmented genomic makeup in terms of functional gain through facilitated horizontal gene transfer, the genome complexity often limits genetic engineering efforts that are essential for strain development. Replicon fusions appear to be an effective strategy for reducing genome complexity while preserving gene content. A caveat of this strategy is that changes in genomic organization can lead to impairment of fundamental cell cycle functions and thus to reduced fitness. The challenge is therefore to avoid such negative effects through appropriate design and/or evolution of the engineered strains, and to well characterize the new chassis in the desired culture conditions.

Our study provides *S. meliloti* strains with bi-and monopartite genome configurations through site-specific recombination-mediated targeted replicon fusions. In the fused replicons, we aimed at retaining organizational properties, such as replichore ratios, GC skew, as well as distribution and orientation of KOPS and coding sequences. In contrast to a natural *S. meliloti* isolate with triple replicon cointegrate that showed a high frequency of revertants by homologous recombination, *^(21)^* our replicon fusion strategy reduced the likelihood of revertants. This enabled investigating spatial DNA organization and spatiotemporal segregation patterns of a monopartite triple-replicon bacterial genome, mostly independent of spontaneously occurring genome rearrangements in the studied strains. These analyses and growth tests showed that this strain can be applied under standard laboratory culture conditions. The strongly reduced fitness of the derivative strain, which is characterized by *oriC* as the sole origin of replication and several deviations from the patterns of replicon architecture, underlines the importance of these architectural rules. However, suppressor mutations that largely restored the fitness of this strain show that suitable strains can also be obtained from an unfavorable design through evolution. These *S. meliloti* strains with monopartite and single origin of replication genome configuration and good growth characteristics are now available for genome engineering applications that were previously hampered or prevented by the tripartite genome structure.

## MATERIALS AND METHODS

### Bacterial strains and cultivation conditions

Bacterial strains used in this study are derivatives of *E. coli* K12 and *S. meliloti* Rm1021 (S7 Table). *E. coli* was grown at 37°C in lysogeny broth (LB) medium. *^(70)^ S. meliloti* strains were cultivated at 30°C in either tryptone yeast extract (TY) medium *^(71)^* or modified morpholinepropanesulfonic acid (MOPS)-buffered minimal medium *^(72)^* at 200 rpm. If required, the following antibiotics were used accordingly: gentamicin (8 µg/ml for *E. coli*, 30 µg/ml for *S. meliloti*), kanamycin (50 µg/ml for *E. coli*, 200 µg/ml for *S. meliloti*), streptomycin (600 µg/ml for *S. meliloti*), spectinomycin (100 µg/ml for *E. coli*, 200 µg/ml for *S. meliloti*) or tetracycline (10 µg/ml for *E. coli* and 3 µg/ml for *S. meliloti*). Solid medium was supplemented with 1.5% (w/v) BD Difco™ technical agar (Fisher Scientific).

### DNA manipulation and plasmid extraction

Plasmids used in this study are listed in S8 Table. Standard molecular techniques were employed for cloning and transfer of nucleic acids. *^(70)^* DNA fragments were PCR amplified using Q5® High-Fidelity DNA Polymerase (New England Biolabs) or Taq DNA Polymerase (New England Biolabs). DNA oligonucleotides were provided by Sigma-Aldrich (USA) and Integrated DNA Technologies (USA) (S11 Table). For DNA purification and gel extractions the E.Z.N.A.® Cycle-Pure Kit (Omega Bio-Tek) and illustra^™^ GFX^™^ PCR DNA and Gel Band Purification Kit (GE Healthcare Life Sciences) were used, respectively. For phosphorylation of the 5′ hydroxyl terminus of PCR amplicons and oligonucleotides T4 Polynucleotide Kinase (Thermo Scientific) was applied. Dephosphorylation of DNA ends was performed by use of FastAP^™^ Thermosensitive Alkaline Phosphatase (Thermo Scientific). Filling in of 5′-overhangs in double stranded DNA to form blunt ends was achieved using the large fragment of DNA Polymerase I (Klenow fragment) (Thermo Scientific). T4 Ligase (Thermo Scientific) was used for ligation. Plasmid DNA was isolated using the “E.Z.N.A. Plasmid Mini Kit” (Omega Bio-Tek). All enzymatic catalyzation and purification steps were performed according to the manufacturer’s protocols and instructions. For sequence verification of plasmid and amplified DNA the sanger sequencing service of Eurofins Genomics (Germany) was used. For detailed information on the construction of individual plasmids refer to S10 Table.

### Strain construction

Strains generated in this study are listed in S9 Table. Transfer of plasmids to *S. meliloti* was achieved by conjugation using *E. coli* S17-1 *^(73)^* or by electroporation as previously described. *^(31)^* Cells for electroporation were prepared as described in Ferri et al.. *^(74)^* Markerless integrations through double homologous recombination were carried out using pK18mobsacB derivatives and sucrose selection. *^(75)^*

On the basis of a Cre/*lox* toolbox and *S. meliloti* Rm1021 derivative SmCreΔhsdR *^(31)^* the tripartite genome was merged in two consecutive steps. First, pSymA and pSymB were fused with each other, giving rise to the megaplasmid hybrid pSymAB harbored by *S. meliloti* strain SmAB. Therefore, pK18mobsacB derivatives pJD98 and pJD99 were used to integrate *lox* sites and antibiotic selection markers into SmCreΔhsdR for the site-specific recombination. After removal of active *lox* sites, SmAB was sequentially transformed with constructs pJD130 and pJD126 again providing *lox* sites and an additional antibiotic selection marker for integration of the chromosome and pSymB. Cre-mediated integration of pSymAB into the chromosome gave rise to SmABC with monopartite genome configuration. Cre/*lox* applications were performed as described before. *^(31)^* Illustration of SmAB and SmABC strain generation and detailed information about the construction process is given in S1 Fig and S9 Table, respectively.

In order to remove DNA replication origins of strain SmABC, deletion constructs pJD201 and pJD202 were used for sequential excision of the megaplasmid-encoded copies of *repC* and corresponding *repBC* intergenic regions. The deletion of both regions in SmABC resulted in strain SmABCΔoriV. For functional inactivation of the inherent partitioning system in pSymA and pSymB, the constructs pMW210 (*repA1B1C1*), pMW211 (*repA2B2C2*) and pJD225 (*repA3B3*) were used to delete the hole operon and the most proximal partitioning sites in SmCreΔhsdR, SmAB and SmABC. In an attempt to delete the partitioning sites upstream of *repA2B2C2*, pMW230 was used. Additionally, the constructs pJD206 (*repA2*), pJD207 (*repA1*) and pJD266 (*repA3*) were used to inactivate the partitioning system and the entire operon by single crossover integrations into the respective RepA coding regions. Gene deletions and plasmid integrations were verified by PCR.

By use of deletion constructs pJD222 and pJD229, *S. meliloti* strains SmAB, SmABC and SmABCΔoriV were further cured from spectinomycin and gentamicin resistance cassettes, respectively, giving rise to strains SmABΔR, SmABCΔR and SmABCΔoriVΔR accessible for constructs of the replicon labeling system. For *in vivo* studies of DNA organization and spatiotemporal dynamics of origin and terminus by fluorescence microscopy, a triple label system based on the fluorescent reporter gene fusions *tetR-mVenus* and *lacI-mCherry* (derived from the FROS *^(46)^*) and *parB-cerulean* was developed. Initially, *S. meliloti* strains SmCreΔhsdR, SmABΔR, SmABCΔR and SmABCΔoriVΔR were transformed with pMW198. Deletion of the plasmid backbone via sucrose selection resulted in an in-frame fusion of the native *parB* gene with *cerulean* (*oriC* label). For analysis of the replisome formation and dynamics, pK19ms DnaN-mCherry *^(13)^* was used for markerless integration of the fluorophore genetic construct. pK18mobsacB derivatives pMW186 (providing 120 copies of the *tetO* array, *tetO*_120_) and pMW188 (carrying 120 copies of the *lacO* array, *lacO*_120_) were used for marker-free labeling of *oriA* (SMa2383-SMa2385 intergenic region) and *oriB* (SMb20041-SMb20042 intergenic region), respectively. pK18mob2 derivatives pJD169 (*tetO*_120_), pMW193 (*lacO*_120_) and pJD170 (*tetO*_120_) enabled labeling of *terC* (SMc01205-SMc01204 intergenic region), *terA* (SMa1188 (*nosX*)-SMa1191 (*hmp*) intergenic region) and *terB* (SMb21555 (*kefB2*)-SMb21556 intergenic region), respectively. Regarding the *ori/ter* labeling, *oriC* pre-labeled strains were transformed with pJD169 (*oriC*/*terC* labelling), pMW186 and pMW193 (*oriC*/*oriA*/*terA* labelling), or pMW188 and pJD170 (*oriC*/*oriB*/*terB* labelling). Additionally, integrative pK18mob2 derivatives pAM25, pAM30, pAM31, pAM32 (Km^R^, *tet0*_120_) and pACYC177 derivatives pAM7, pAM13, pAM45, pAM47, pAM68, pAM69, pAM70 and pAM72 (Gm^R^, *lacO*_120_) were used to label further genomic loci in context of the genome organization analysis. Repair of CckA^R436H^ in SmABCΔoriVΔR was attempted using a double homologous recombination strategy integrating pMW257 for markerless replacement of the *cckA* SNV (C→T) with wild type coding sequence of *cckA*. Reproduction of the CckA^R436H^ mutation in strains of the wild type and SmABCΔR was achieved by markerless integration of *cckA* SNV (C → T) trough pMW256. Analysis of *oriC* localization in strains SmABC and SmABCΔoriV was realized by single crossover integration of pMW261.

### Strain validation

Pulsed-Field Gel Electrophoresis (PFGE) was used as method to validate the genome architecture after major fusion and integration/deletion steps. The applied PFGE protocol for DNA preparation and digestion was carried out as described for strain validations in Checcucci et al.. *^(20)^* To gain fusion strain characteristic banding patterns the restriction digestion of genomic DNA was performed with PacI (New England Biolabs, USA). For PFGE analysis, ¼ agarose plug with treated genomic DNA was separated in an 0.7% agarose gel (Pulse Field Certified Agarose, Bio-Rad, USA) and 0.5x TBE buffer at 12°C (44.5mM Tris-HCl, 44.5mM boric acid, 1mM EDTA) using the Rotaphor® System 6.0 (Analytik Jena, Germany) according to the manufacturer’s instructions. Separation of DNA fragments was achieved with 130V-100V for 50-175sec at 130°-110° (run time 18h), 130V-80V for 175sec-500sec at 110° (run time 18h) and 80V-50V for 500sec-2000sec at 106° (run time 40h) with a logarithmic course of increase or decrease between varying parameters, respectively.

For sequence specific analysis, including verification of proper deletions and detection of single-nucleotide variants (SNVs), all basic strains (SmCreΔhsdR, SmAB, SmABC, SmABCΔoriV, SmABΔ, SmABCΔR and SmABCΔoriVΔR) were subjected to next generation DNA sequencing using the MiSeq™ System (Illumina, USA). For preparation of genomic DNA, *S. meliloti* cells were grown in TY supplemented with appropriate antibiotics and harvested at optical density_600_ (OD_600_) of 1.0 by centrifugation at 3000g (4°C). Sample preparation was performed as previously described. *^(76)^* The investigation for single nucleotide variations was carried out using the Basic Variant Detection tool (v.2.1) of CLC genomic workbench (v.20.0.4) with a minimum coverage of eight, minimum count of four and minimum frequency of 50% for mapped reads. Next, each SNV was manually analyzed by eye and, in case of doubt, additionally validated by Sanger sequencing. Assembly of the SmABCΔoriVΔR_Inv_ genome was performed using data from a whole genome nanopore sequencing approach (MinION; Oxford Nanopore Technologies plc) and the *de novo* assembly algorithm for long reads of the CLC Genomic Workbench (v.22.0.2). The single contig genomic scaffold was then refined with data (short read) from the MiSeq™ system using the polish with reads tool.

### Growth experiments

Prior to inoculation, overnight cultures were washed with 0.9 % NaCl and adjusted to OD_600_ of 0.01-0.15 in TY or MOPS buffered medium supplemented with 600 mg/ml streptomycin. Cultures were incubated in a 100 µl volume in a 96 well microtiter plate at 30°C and with shaking at 200 rpm. OD_600_ of cell cultures was measured every 30 min with a microplate reader (Tecan Infinite 200 PRO, Tecan, Switzerland).

### Live cell microscopy and image analysis

*S. meliloti* cell cultures were grown in TY medium (glass tubes) supplemented with suitable antibiotics to OD_600_ of 0.25 for time lapse and OD_600_ of 0.5 for snapshot microscopy. To enrich cultures with G_1_-phase cells for snapshot analysis, strains were grown to OD_600_ of 1.6-1.8, diluted to OD_600_ of 0.5 and subsequently used for microscopy. 1 µl of these cultures were then placed onto 1 % (w/v) molecular biology-grade agarose (Eurogentec, Belgium) pads containing ddH_2_O (snap shots) or MOPS minimal medium and suitable antibiotics (time lapse), covered with a cover glass and sealed with VALAP. *^(77)^* For visual examination of *S. meliloti* cells by phase contrast and epifluorescence microscopy an Eclipse Ti-E inverse research microscope (Nikon, Japan) equipped with a 100x CFI Plan Apo1 oil objective (numerical aperture of 1.45), a green DPSS solid state laser (561 nm, 50 mW; Sapphire) and a multiline Argonlaser (457/488/514 nm, 65 mW; Melles Griot) with AHF HC filter sets F36-513 DAPI (excitation band pass [ex bp] 387/11 nm, beam splitter [bs] 409 nm, emission [em] bp 447/60 nm), F36-504 mCherry (ex bp 562/40 nm, bs 593 nm, em 624/40 nm), F36-528 mVenus (ex bp 500/24 nm, bs 520 nm, and em bp 542/27 nm) was used. Exposure times ranged from 200 ms to 2 s. Image acquisition and adjustment was done with an Andor iXon3 885 electron-multiplyingcharge-coupled device (EMCCD) camera and the software NIS-Elements v.4.13 (Nikon, Japan), respectively. Time-lapse analysis was performed at 30°C in a microscope incubator and images were acquired every 2 or 5 minutes. Analysis of snap-shot and time-lapse microscopy images was performed using ImageJ plug-in MicrobeJ. *^(78)^* G_1_-phase cells were filtered for a maximal length of 2.0 µm and presence a single ParB-cerulean focus indicative of non-segregated *oriC*.

### Marker frequency analysis

For marker frequency analysis, *S. meliloti* strains SmABC and SmABCΔoriV were grown in 50 ml TY medium supplemented with 600 mg/ml streptomycin at an initial OD_600_ of 0.1. After incubation at 30°C and 200 rpm, samples were taken at OD_600_ of 0.6 (exponential phase) or OD_600_ of ∼ 2.6 (overnight culture, stationary phase). Cells were harvested by centrifugation (4000 g, 4°C) and immediately frozen in liquid nitrogen. Preparation and acquisition of Illumina Miseq data were performed as previously described. *^(76)^* Paired-end reads were then mapped by the QuasR R package (v1.6.2) onto the *S. meliloti* replicons. Only unique hits were considered. Subsequently, the coverage was determined from the obtained mapped genomic DNA reads using the genomecov from the bedtools toolbox (v2.25.0). The average coverage of all samples ranged between 18 and 23. The coverage was normalized by the total coverage (sum of coverage) of each sample. To identify minimal variations in the copy number along the replicons, we used sliding window averaging. The size of the window comprised 200 kb. After averaging, the value at a certain position reflects the average coverage of about 2 % of the replicon left and right of the indicated position. This process averages out random noise and local sequence specific variation. To determine the copy number without prior information about the terminus region, the lower 10 % quantile of all windows was used to determine the reads in the terminus region. All windows where then normalized by this value, resulting in a copy number relative to the terminus region.

### Modeling

For the simulations we used a model for DNA described by Buenemann and Lenz. *^(56)^* The basic assumptions of the model are 1) DNA can be modeled as a sequence of compacted units (S1 Text); 2) compact units can be restricted in their spatial arrangement e.g. by the action of proteins; and 3) the measured organization of the chromosome in the cell results from averaging over many individual configurations that meet these constraints. For *C. crescentus* this model revealed that self-avoidance of DNA, specific positioning of the origin (and terminus) region and the compaction of DNA are sufficient to explain the strong linear correlation between specific positions on the chromosome and their longitudinal arrangement within the cell. *^(57)^* To predict the spatial organization of the merged replicons in the *S. meliloti* replicon fusion strain SmABC an expansion of the model by implementing not only one origin and one terminus as fixpoints, but three each was done. For realization, the A* algorithm was added to the model, which made it possible to generate random walks between any number of fixed points (S2 Text).

### Bioinformatic analysis

GC and GC^c^ skew analysis of the tri-, bi- and monopartite *S. meliloti* genome was performed using GenSkew (http://genskew.csb.univie.ac.at, Feb. 2020). GC skew depictions of the individual replicons as shown in Fig 1 were generated using the CGView server (http://stothard.afns.ualberta.ca/cgview_server/ Feb. 2020). Oligonucleotide skews for KOPS were calculated with fuzznuc (http://emboss.toulouse.inra.fr/cgi-bin/emboss/fuzznuc?_pref_hide_optional=1, Mar. 2020). Functional domain analysis of *S. meliloti* CckA was done using the NCBI conserved domain database (CDSEARCH/cdd) *^(79)^* with low complexity filter, composition-based adjustment and an E-value threshold of 0.01 (Oct. 2021).

## ASSOCIATED CONTENT

### Data Availability Statement

The sequence data presented in this article are available at the ArrayExpress Archive under the accession number E-MTAB-11878

## AUTHOR INFORMATION

### Corresponding Author

*Phone: +49 6421 28-24451.

### Author Contributions

AB, MW and JD designed the experiments. MW and JD carried out the experiments and analyzed the experimental data. PS performed computational analysis of sequence data for MFA. MW and JS analyzed genome sequence data. DG and PL modeled the spatial organization of DNA. AB and MW wrote the paper.

### Notes

The authors declare no competing financial interest.

## ACKNOWLEDGMENTS

Whole genome sequencing was performed by the Screening and Automation Technology Core Facility (SAT) of the Center for Synthetic Microbiology (SYNMIKRO). We thank Bernadette Boomers for technical assistance, Elizaveta Krol for providing primers, plasmids pG18mob-mVenus and pSRKGm-CmChr, Anna Motnenko for providing pAM labeling plasmids, Yannick End for plasmid pYEM1, Vanessa Munoz-Gutierrez for providing primers and Patrick Manz for technical support related to microscopy. This study was supported by the Collaborative Research Centre TRR 174 “Spatiotemporal dynamics of bacterial cells” (German Research Foundation) and the LOEWE program of the State of Hesse (Germany).

## ABBREVIATIONS

*ori*: origin
*ter*: terminus
SNV: single nucleotide variation
MFA: marker frequency analysis.

## SUPPORTING INFORMATION

**S1 Fig.** Construction of *S. meliloti* SmAB and SmABC (PDF)

**S2 Fig.** Visualization of the asymmetric nucleotide composition in SmCreΔhsdR (wt), SmAB and SmABC (PDF)

**S3 Fig.** Gene orientation bias in SmCreΔhsdR (wt), SmAB and SmABC (PDF)

**S4 Fig.** Oligonucleotide bias of FtsK orienting polar sequences (KOPS) (PDF)

**S5 Fig.** Cell shape of SmCreΔhsdR (wt), SmAB, SmABC in TY and TY with 0.4M NaCl (PDF)

**S6 Fig.** Growth of SmCreΔhsdR (wt), SmAB and SmABC (PDF)

**S7 Fig.** Characterization of *S. meliloti* replicon fusion strains lacking *repC* and *repBC* intergenic region (PDF)

**S8 Fig.** Marker frequency analysis to assess origin activities and location of replication terminus region in the wild type strain SmCreΔhsdR (A), SmABC (B) and SmABCΔoriV (C) (PDF)

**S9 Fig.** Replisome formation and dynamics in SmABCΔR vs SmABCΔoriVΔR (PDF) **S10 Fig.** Establishment of a genome architecture independent labeling strategy (PDF) **S11 Fig.** Genome organization in SmABCΔR compared to the wild type (PDF)

**S12 Fig.** Modeling of spatial DNA arrangement in SmABCΔR (PDF).

**S13 Fig.** Cumulating time-lapse data of SmCreΔhsdR (wt), SmAB (AB), SmABC (ABC) and SmABCΔoriVΔR (ABCΔ) daughter cells (PDF)

**S14 Fig.** *oriA* and *oriB* segregation (PDF)

**S15 Fig.** Comparison of *oriC*, *oriA/*Δ*oriA* and *oriB/*Δ*oriB* distribution in SmCreΔhsdR (wt), SmABΔR, SmABCΔR and SmABCΔoriVΔR (PDF)

**S16 Fig.** Origin co-localization scheme for SmCreΔhsdR (wt), SmABΔR, SmABCΔR and SmABCΔoriVΔR (PDF)

**S17 Fig.** *oriC* localization pattern in SmABCΔoriVΔR cells after one cell-division cycle (PDF)

**S18 Fig.** Analysis of CckA^R436H^ in *S. meliloti* wild type and genome fusion strains (PDF)

**S1 Table.** Underlying data of S11B Fig (PDF)

**S2 Table.** Timepoint and Localization of origin and terminus foci segregation in SmCreΔhsdR (wt), SmAB, SmABC and SmABCΔoriVΔR mother cells [M] (PDF)

**S3 Table.** Values corresponding to S13A Fig. daughter cells [D] (PDF)

**S4 Table.** Coding sequences that were removed or truncated upon the replicon fusion procedure (PDF)

**S5 Table.** Single nucleotide variations associated with the fusion procedure (PDF)

**S6 Table.** Single nucleotide variations and nucleotide deletions in *S. meliloti* genome fusion strains (PDF)

**S7 Table.** Bacterial strains used in this study (PDF)

**S8 Table.** Plasmids used in this study (PDF)

**S9 Table.** Construction of *S. meliloti* replicon fusion strains and derivatives (PDF)

**S10 Table.** Plasmid construction (PDF)

**S11 Table.** Oligonucleotides used in this study (PDF)

**S1 Data.** Validation of antibiotic marker deletion and genome configuration in *S. meliloti* fusion strains used for microscopy (PDF)

**S2 Data.** Visualization of single cell time-lapse data in SmCreΔhsdR (A), SmABΔR (B), SmABCΔR (C) and SmABCΔoriVΔR (D) during the cell cycle (PDF)

**S1 Text.** Model of compacted DNA (PDF)

**S2 Text.** Monte Carlo sampling of configuration space (PDF)

**S3 Text.** SI References (PDF)

