## Supplemental for "Engineering a *Sinorhizobium meliloti* chassis with monopartite, single replicon genome configuration"

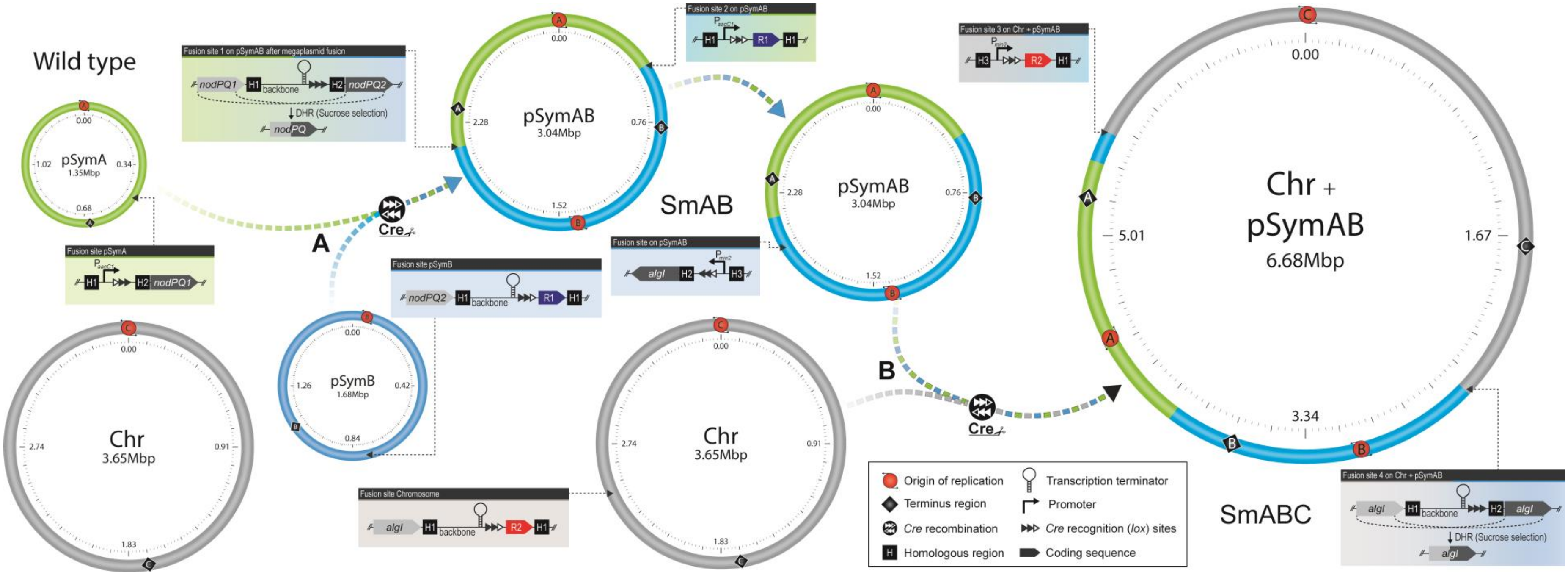

**S1 Fig. Construction of *S. meliloti* SmAB and SmABC. (A)** Initially, *S. meliloti* SmCreΔhsdR (wild type) was transformed with integrative pK18mobsacB derivatives pJD98 and pJD99, thereby providing *loxL* and *loxR* sites (black triangles, non-filled indicates mutation) at 125°(pSymA) and 169°(pSymB) for Cre-mediated megaplasmid fusion and components of a spectinomycin resistance cassette allowing for positive selection after Cre/*lox* recombination. The plasmid backbone of pJD99 was deleted via *sacB* mediated sucrose selection. In contrast, the backbone of pJD98 was retained to enable sucrose selection in a later step. Due to Cre-mediated replicon fusion, the homologous *nodPQ1*(SMa0855/SMa0857) and *nodPQ2* (SMb21223/SMb21224) operon came into close proximity, flanking the pJD98 derived plasmid backbone, thereby enabling sucrose selection-mediated deletion of the vector backbone and a wild type *loxP* site (resulting from *loxL/R* mediated Cre recombination). Thus, the resulting strain SmAB could be easily used for further Cre/*lox* reactions. $P_{aacC1}$: Constitutive resistance cassette promoter. **(B)** *S. meliloti* strain SmAB was then transformed with pK18mobsacB derivative pJD130 which provides the hybrid replicon pSymAB with a *loxL* site at 238° followed by the constitutive promoter P*min2*. Sucrose selection-mediated deletion of the plasmid backbone resulted in strain JDSm111 which was then transformed with pJD126. In this way the chromosome was equipped with a *loxR* site at 246° and the promoterless gentamicin resistance gene *aacC1*, thereby allowing for selection on gentamicin resistance after Cre/*lox* recombination. Thus, the resulting strain SmABC exhibited an entirely merged genome and antibiotic resistances against tetracycline, spectinomycin and gentamicin. *lox* sites were integrated close to the *algI* homologs of pSymB (SMb20843) and the chromosome (SMc01551), thereby enabling sucrose selection-mediated deletion of a remaining *loxP* site and the plasmid backbone. *A/B/C:* DNA replication origin/terminus region of pSymA/pSymB/chromosome. Mbp: Megabase pairs. R: antibiotic resistance gene. Color code: chromosome (grey), pSymA (green), pSymB (blue).

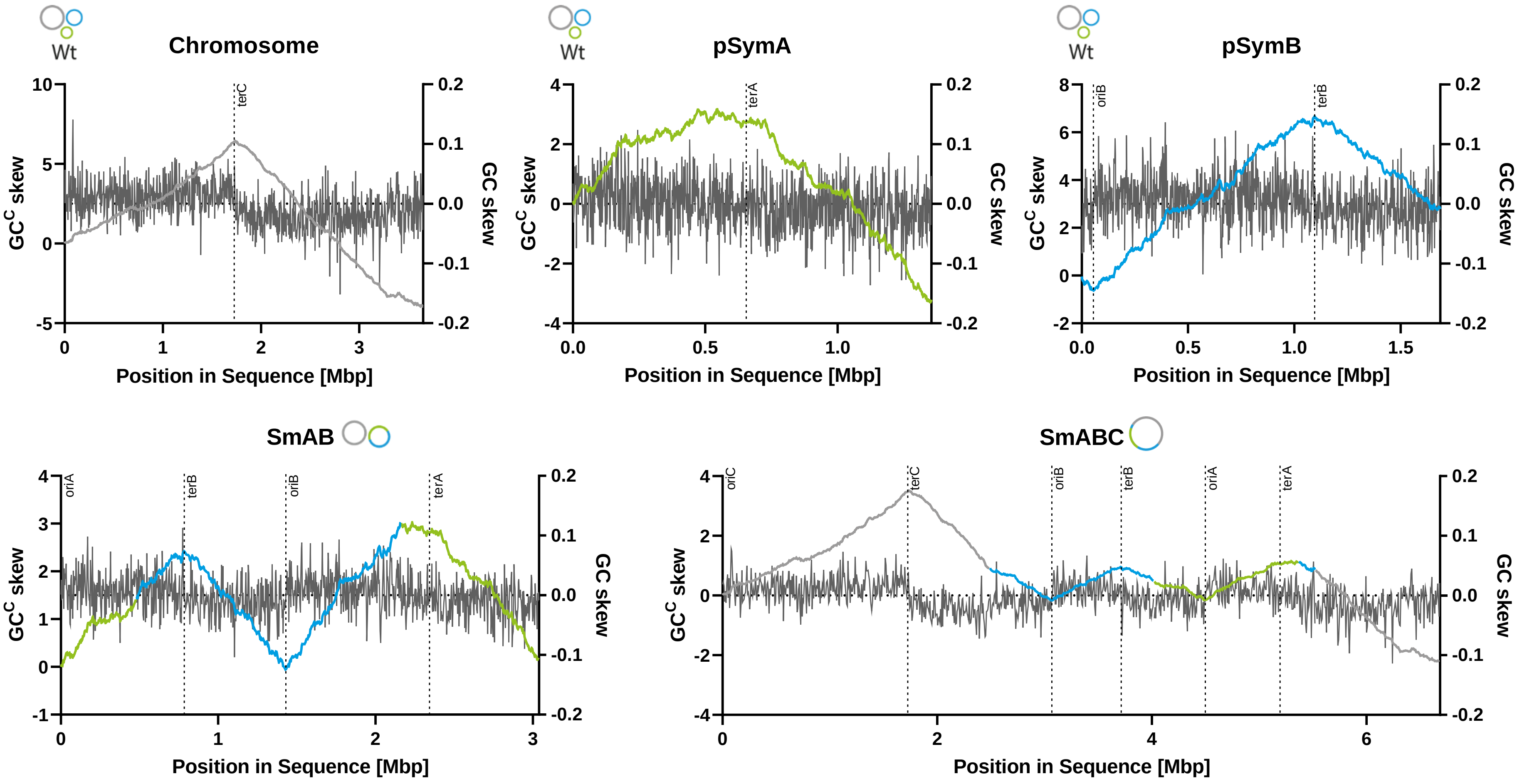

**S2 Fig. Visualization of the asymmetric nucleotide composition in SmCreΔhsdR (wt), SmAB and SmABC.** The abundance of nucleotides is represented by the GC skew value of each replicon sequence. Analyzed sequences were subdivided into 1000 parts (windows) and the GC skew was calculated for each window as (G – C) / (G + C). The GC skew graph depicts the value of individual windows at certain positions moving with a defined step size along the analyzed sequence. The cumulative GC skew ($GC^C$ skew) was calculated by addition of all GC-values from adjacent windows up to a specific position thereby most likely representing the replichore bias with minimum values at origin and maximum values at terminus regions. The underlying nucleotide skew data were generated using GenSkew (http://genskew.csb.univie.ac.at). Analysis parameter: SmCreΔhsdR (Chromosome): GC-content: 62.7 %, window and step size: 3650 bp, max. $GC^C$ value: 6.44 (Pos: 1.73 Mbp), min. $GC^C$ value: -3.94 (3.65 Mbp). SmCreΔhsdR (pSymA): GC-content: 60.4 %, window and step size: 1354 bp, max. $GC^C$ value: 3.16 (Pos: 0.55 Mbp), min. $GC^C$ value: -3.29 (1.35 Mbp). SmCreΔhsdR (pSmyB): GC-content: 62.4 %, window and step size: 1686 bp, max. $GC^C$ value: 6.65 (Pos: 1.10 Mbp), min. $GC^C$ value: -0.64 (0.05 Mbp). SmAB (pSymAB): GC-content: 61.5 %, window and step size: 3038 bp, max. $GC^C$ value: 3.02 (Pos: 2.23 Mbp), min. $GC^C$ value: -0.06 (1.43 Mbp). SmABC (Chr. + pSymAB): GC-content: 62.2 %, window and step size: 6683 bp, max. $GC^C$ value: 3.52 (Pos: 1.74 Mbp), min. $GC^C$ value: -2.23 (6.68 Mbp). Positions of terminus regions are based on architecture imparting motif sequence (AIMS) predictions (Hendrickson & Lawrence, 2006). Color code: chromosome (grey), pSymA (green), pSymB (blue).

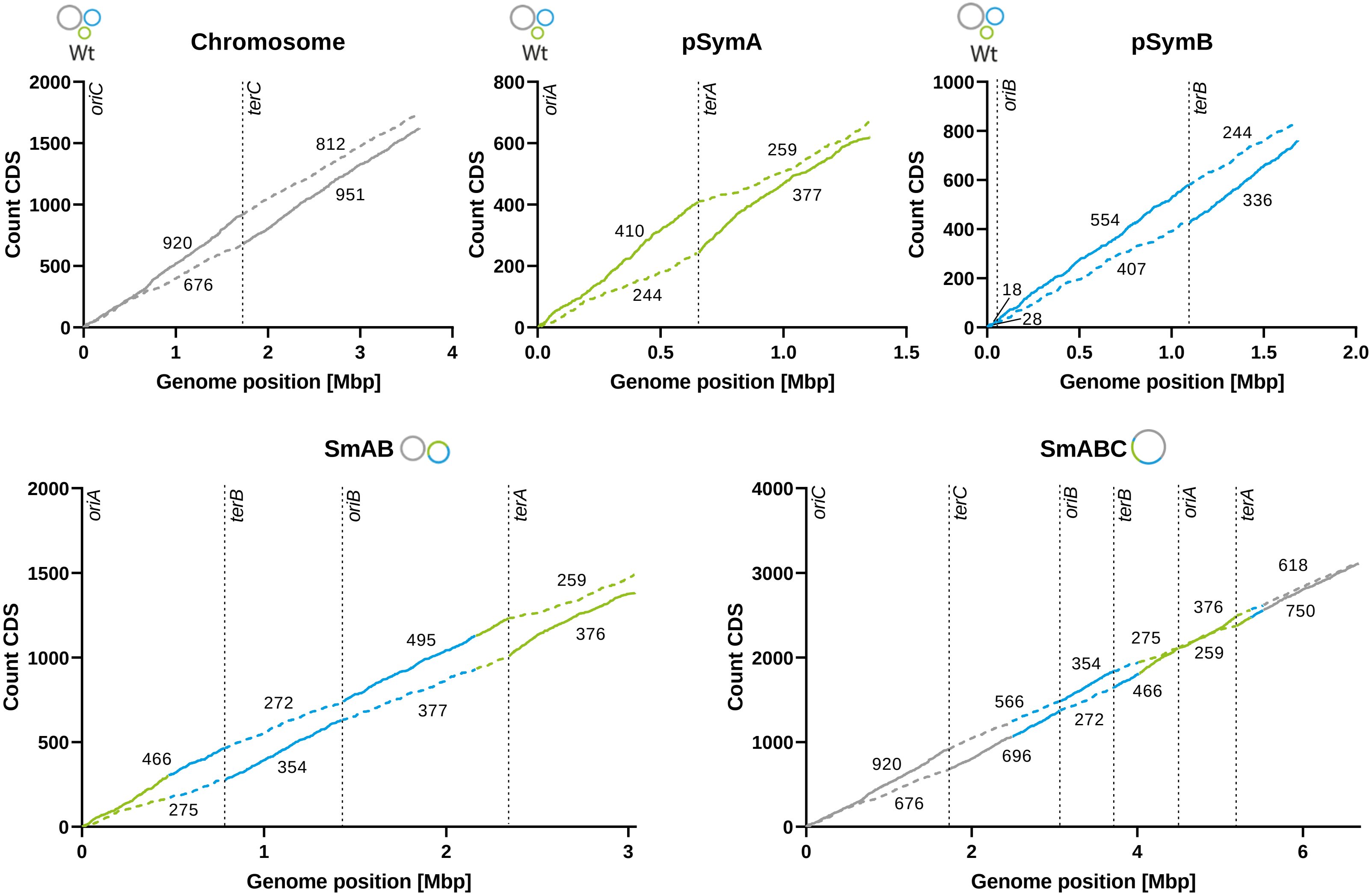

**S3 Fig. Gene orientation bias in SmCreΔhsdR (wt), SmAB and SmABC.** The number of coding sequences (CDS) located on the individual replicons revealed a gene strand bias between the leading (solid line) and the lagging (dashed line) strand of a replichore with genes predominantly accumulating on the leading strand. SmCreΔhsdR (wt) Chr: 1871 CDS (56 %) on the leading and 1488 CDS (44 %) on the lagging strand, pSymA: 787 CDS (61 %) on the leading and 503 CDS (39 %) on the lagging strand, pSymB: 918 CDS (58 %) on the leading and 669 CDS (42 %) on the lagging strand. SmAB (pSymAB): 1691 CDS (59 %) on the leading and 1183 CDS (41 %) on the lagging strands. SmABC (ABC): 3562 CDS (57 %) on the leading and 2666 CDS (43 %) on the lagging strands. Terminus regions depicted as predicted by Hendrickson & Lawrence, 2006. Color code: chromosome (grey), pSymA (green), pSymB (blue). *oriA*/*B*/*C*: origin of replication, *terA*/*B*/*C*: terminus region.

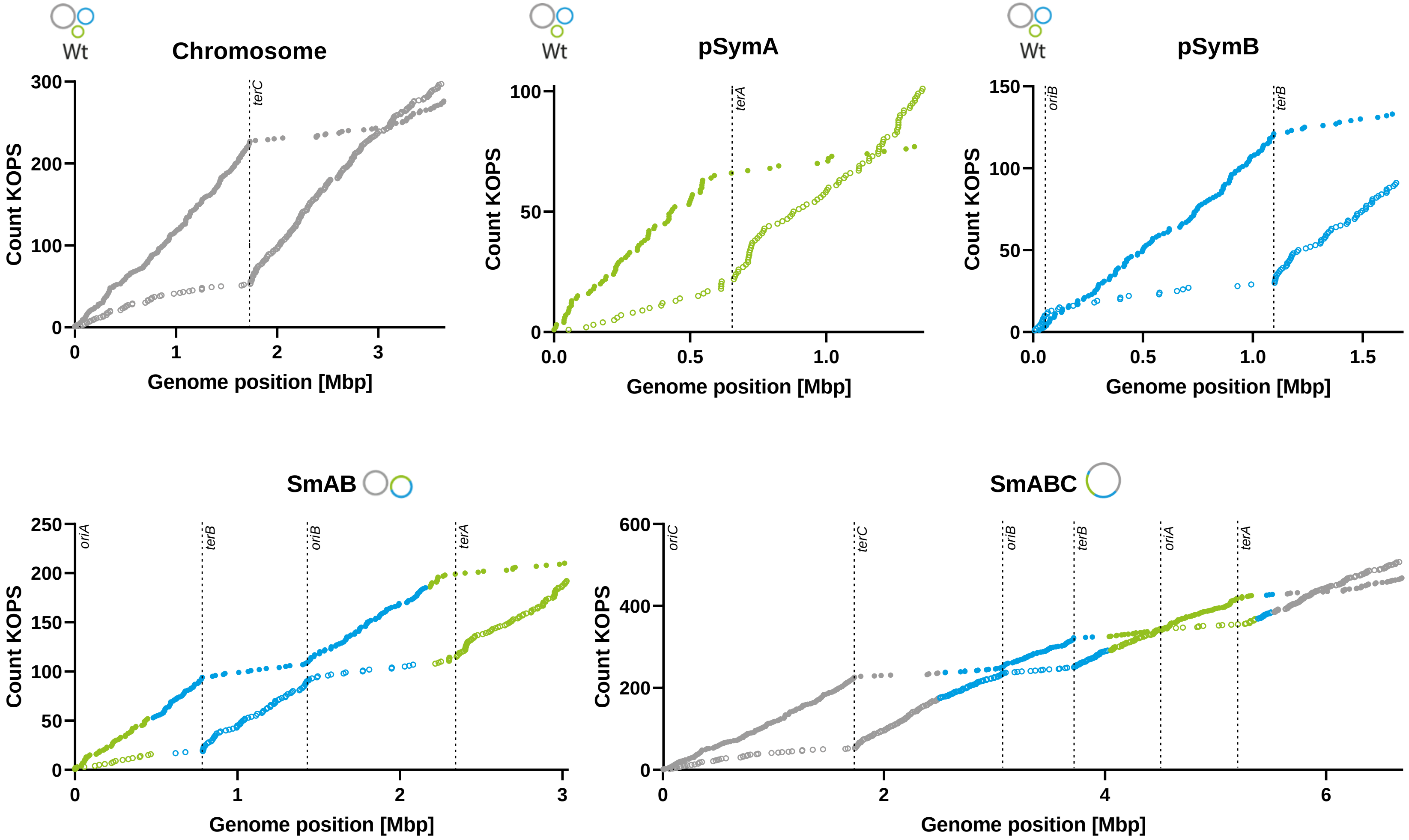

**S4 Fig. Oligonucleotide bias of FtsK orienting polar sequences (KOPS).** Count of the *E. coli* KOPS consensus motif GGGNAGGG (Bigot et al., 2005) on the forward (filled circles) and complementary strand (non-filled circles) respectively is depicted as a function of the position on the genome sequence. The KOPS sequence as an architecture imparting motif was used to display the strand asymmetry bias on the individual replichores. KOPS distribution was analyzed by use of fuzznuc (http://emboss.toulouse.inra.fr/cgi-bin/emboss/fuzznuc?_pref_hide_optional=1). Terminus regions depicted as predicted by Hendrickson & Lawrence, 2006. Color code: chromosome (grey), pSymA (green), pSymB (blue). *oriA*/*B*/*C*: origin of replication, *terA*/*B*/*C*: terminus region. Mbp: megabase pairs.

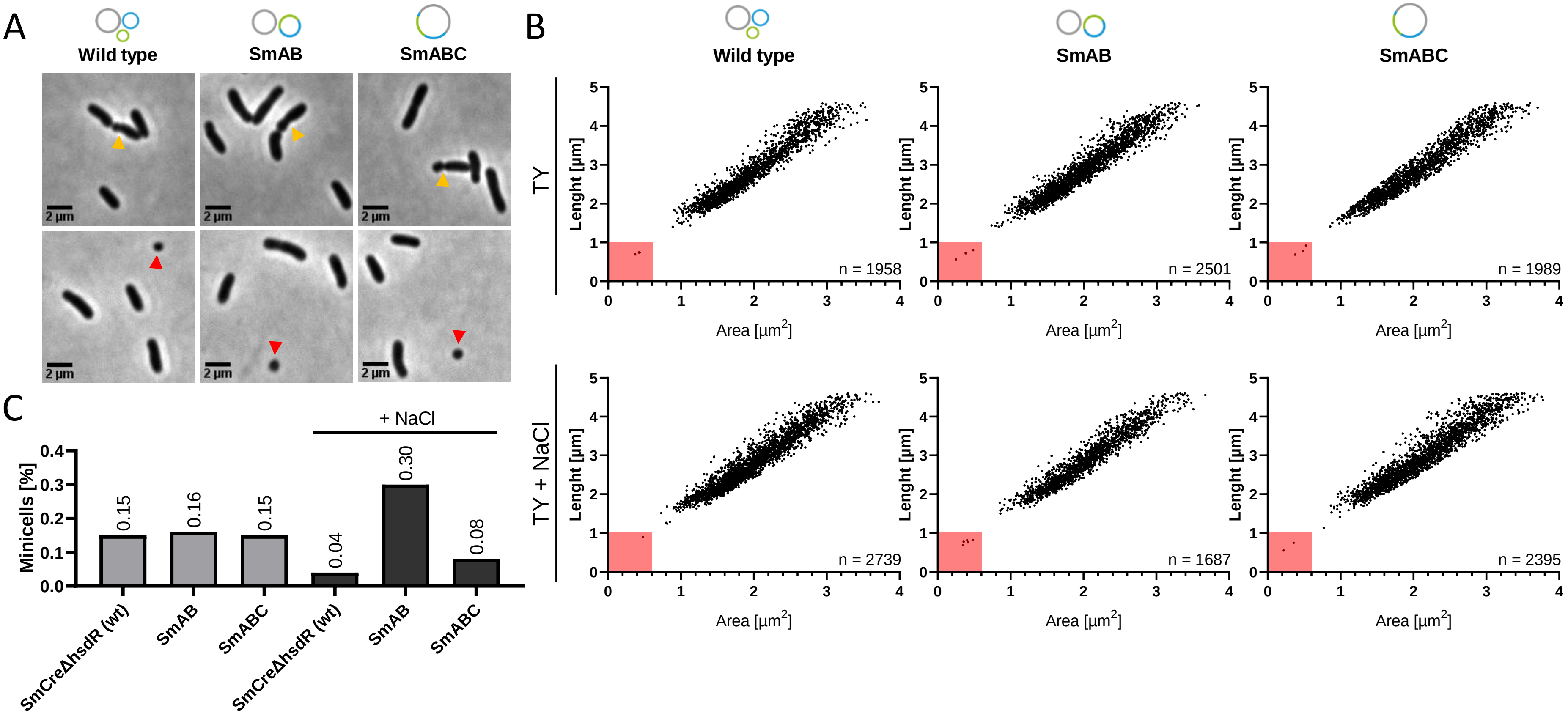

**S5 Fig. Cell shape of SmCreΔhsdR (wt), SmAB, SmABC in TY and TY with 0.4M NaCl. (A)** Microscopic snapshot analysis revealed a small subpopulation of cells with misplaced division septum (orange arrowhead) probably resulting in coccus-shaped cell types here termed as minicells (red arrowhead). Scale bar: 2 µm. **(B)** Scatter plot with cell length [µm] as a function of cell area [µm$^2$] demonstrate the size distribution of SmCreΔhsdR (wt), SmAB and SmABC strains in an exponential growth phase culture. The proportion of minicells defined with cell length <1 µm and area <0.6 µm$^2$ is indicated by the red square. **(C)** Percentage of minicells in TY with wt: 0.15 % ([n=1958), SmAB: 0.16 % (n=2501), SmABC 0,15 % (n=1989) and TY [0.4 M NaCl] with wt: 0.04 % (n=2739), SmAB: 0.3 % (n=1687), SmABC: 0.08 % (n=2395).

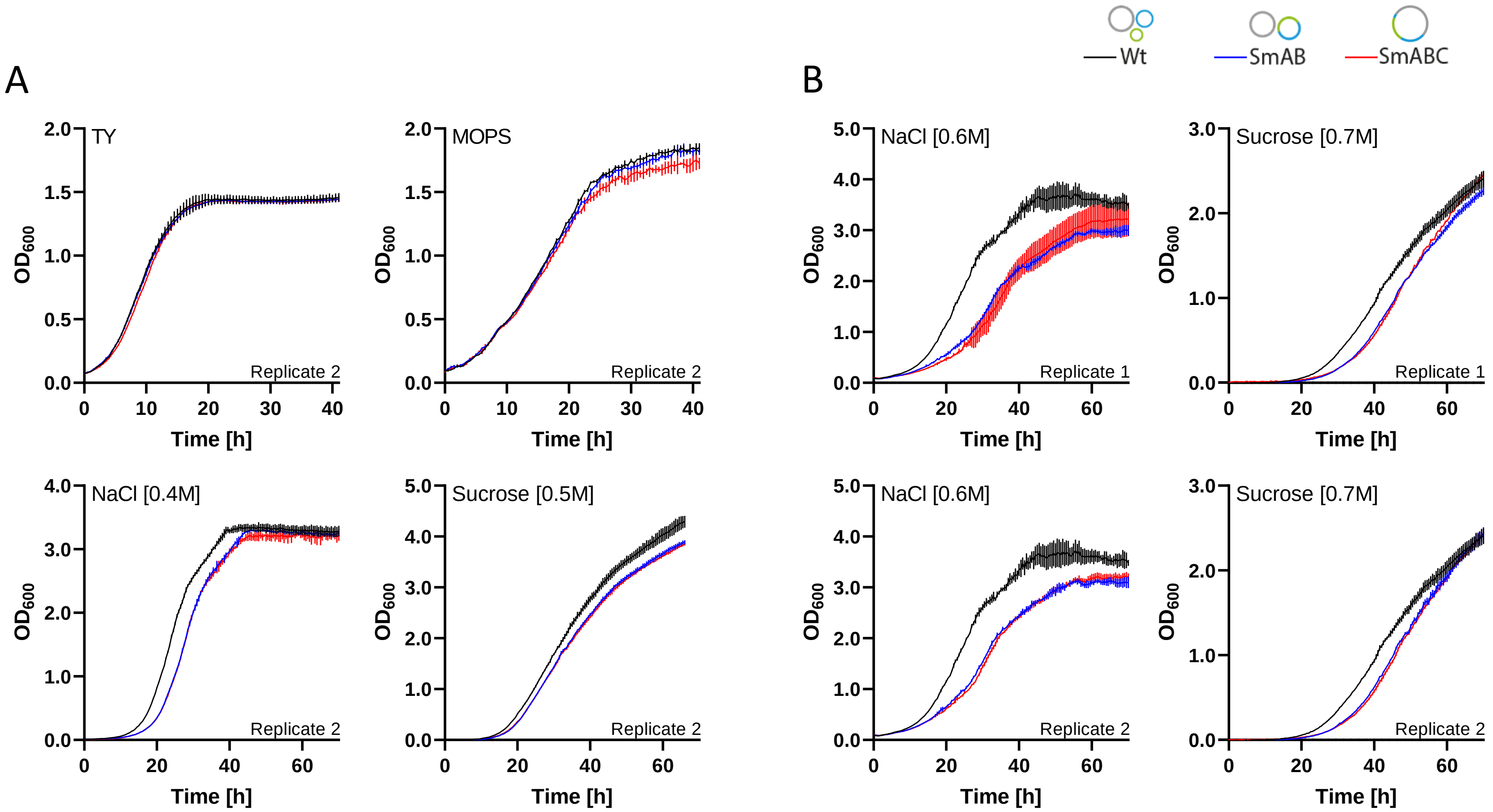

**S6 Fig. Growth of SmCreΔhsdR (wt), SmAB and SmABC**. **(A)** Growth of SmAB and SmABC replicate 2 compared to the precursor strain SmCreΔhsdR (wt) with wild type genome configuration as a control in rich medium (TY), low phosphate minimal medium (MOPS) and TY supplemented with either 0.4 M NaCl or 0.5 M sucrose. **(B)** Hypersaline (0.6 M NaCl) and high sugar (0.7 M sucrose) condition revealed an increasing negative effect on growth behavior for strains SmAB and SmABC. However, even SmCreΔhsdR with wild type-like genome configuration exhibit a diminished growth capacity under this stress conditions. For the analysis, cell cultures were grown for 40 h and 70 h, respectively at 30 °C under shaking conditions (200 rpm). Optical density was measured every 30 minutes at 600 nm ($OD_{600}$). Prior to inoculation, overnight cultures were washed with 0.9 % NaCl and adjusted to an $OD_{600}$ ~ 0.01 (TY+ 0.4 M NaCl, TY+ 0.5 M and 0.7 M sucrose) or 0.1 (TY, MOPS and TY+ 0.6 M NaCl) in respective media supplemented with 600 µg/ml streptomycin. Error bars indicate standard deviation calculated from three technical replicates.

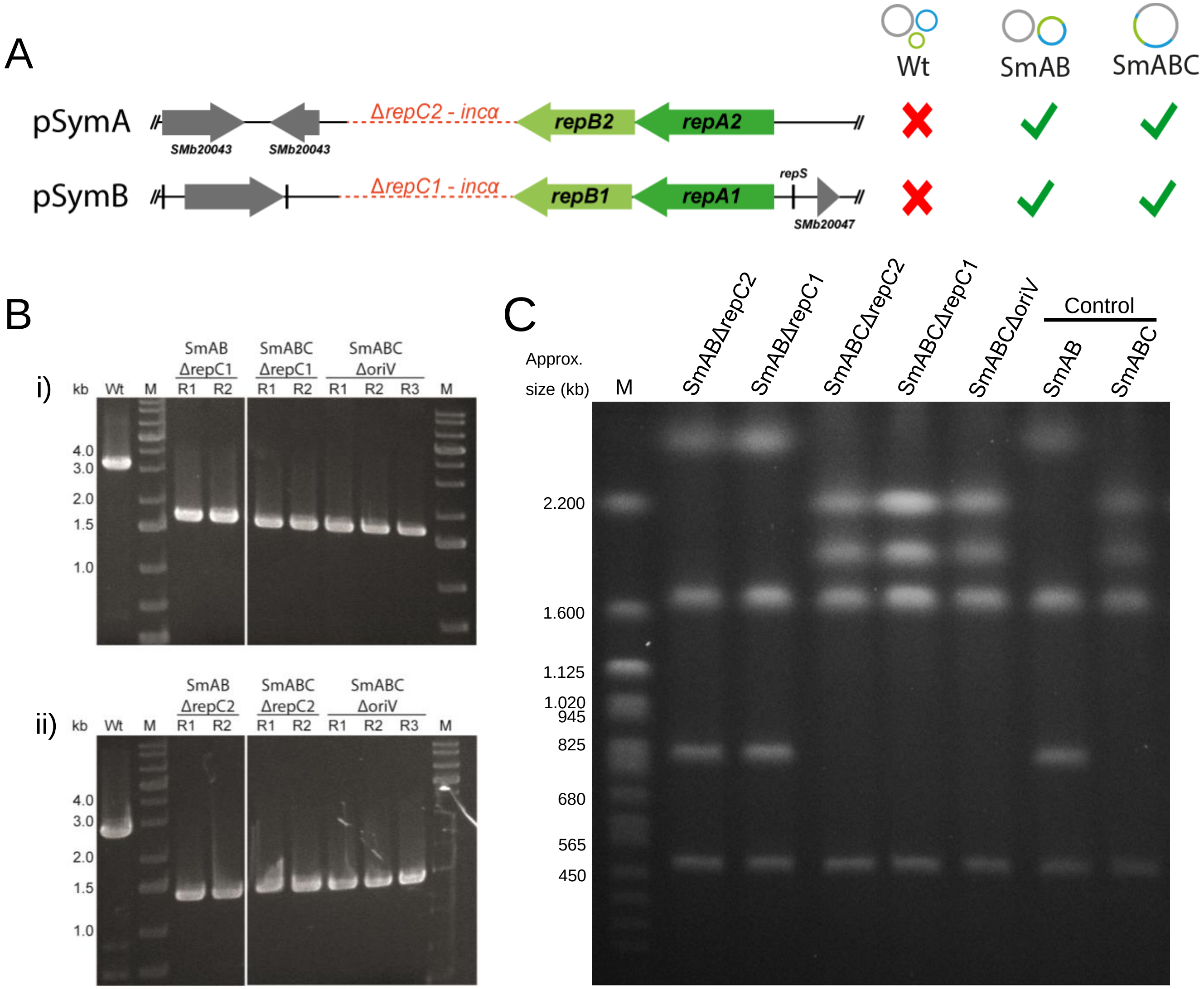

**S7 Fig. Characterization of *S. meliloti* replicon fusion strains lacking *repC* and *repBC* intergenic region. (A)** *repC* deletion constructs pJD202 (Δ*repC1*) and pJD201 (Δ*repC2*) enabled deletion of the secondary replicon *oriV*s in *S. meliloti* strains SmAB and SmABC but not in the wild type. **(B)** Proper deletion in the corresponding strains SmABΔrepC1/2, SmABCΔrepC1/2 and SmABCΔoriV (lacking both *repC1* and *repC2*) was verified via PCR amplification of the deletion site using primers i) 852+853 (wt: 3.15 kb, Δ*repC1*:1.67 kb) and ii) 850+851 (wt: 2.85 kb, Δ*repC2*: 1.39 kb). Additionaly, *repC1/2* deletion in SmABCΔoriV was confirmed using paired-end MiSeq Illumina® sequencing. **(C)** Pulsed-field gel electrophoresis (0.5 x TBE, 0.7 % agarose, separation 72 h) banding pattern of PacI-digested gDNA from *S. meliloti* SmAB derivatives (fragments sizes in Mbp: 3.65, 1.67, 0.83, 0.53) and SmABC derivatives (fragment sizes in Mbp: 2.53, 1.94, 1.67, 0.53) verifies the expected genome configuration. M: PFGE marker *S. cerevisiae*, Bio-Rad.

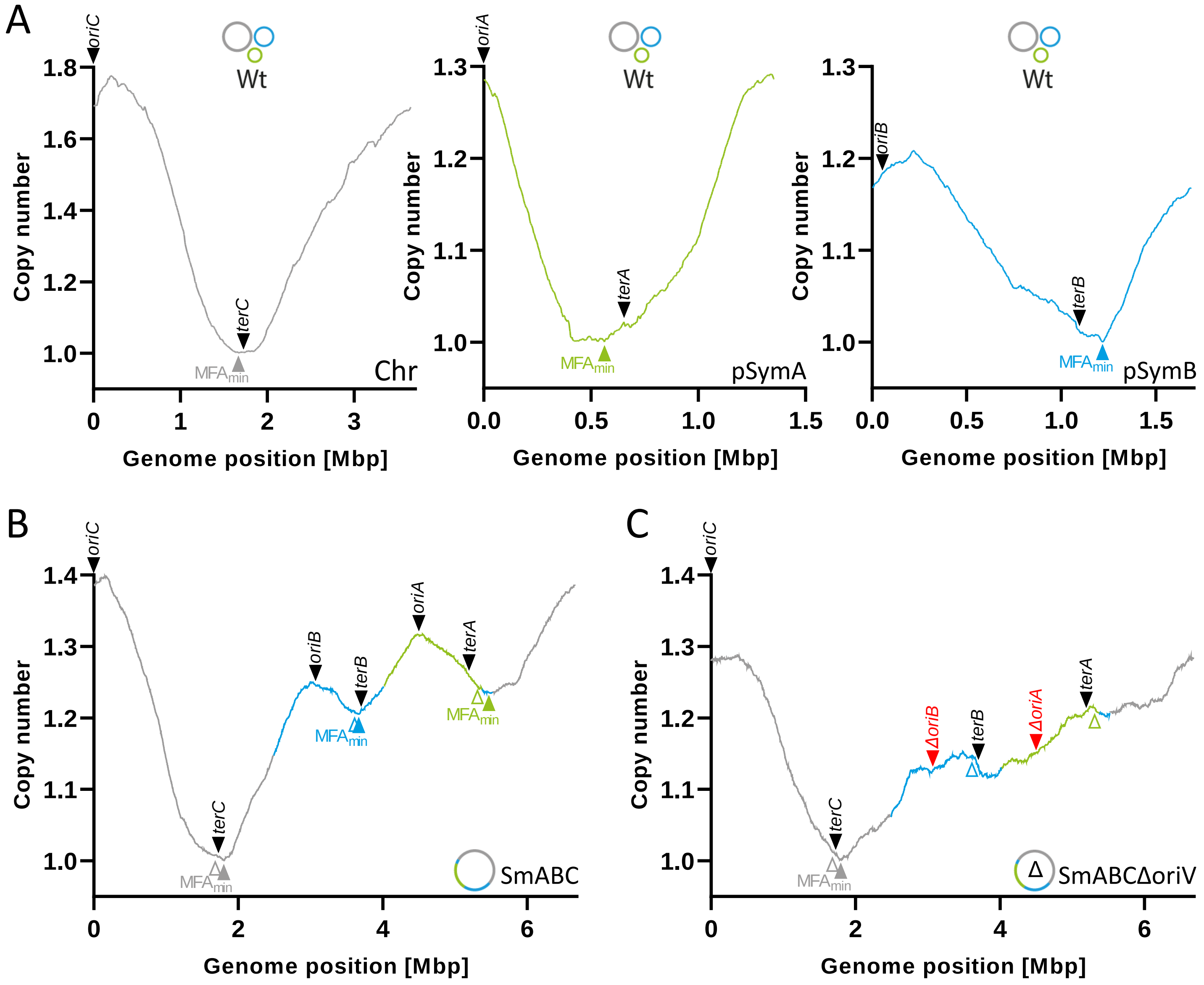

**S8 Fig. Marker frequency analysis to assess origin activities and location of replication terminus region in the wild type strain SmCreΔhsdR (A), SmABC (B) and SmABCΔoriV (C).** Shown are the averaged copy number profiles based on Illumina sequencing as the log2 ratio of each nucleotide in sequences calculated using a sliding window of 200 bp. Black arrowheads depict the position of individual origin regions (Sibley et al., 2006, Cervantes-Rivera et al., 2011) and terminus regions as predicted by Hendrickson and Lawrence, 2006. Red arrowheads in the profile of SmABCΔoriV represent the deleted *oriA* (Δ*oriA*) and *oriB* (Δ*oriB*) region. Colored (grey, blue, green) arrowheads indicate the local minima of the marker frequency analysis ($MFA_{min}$) in each strain. Non-filled arrowheads represent the position of the wild type $MFA_{min}$ in profiles of SmABC and SmABCΔoriV. Color code: chromosome (grey), pSymA (green), pSymB (blue).

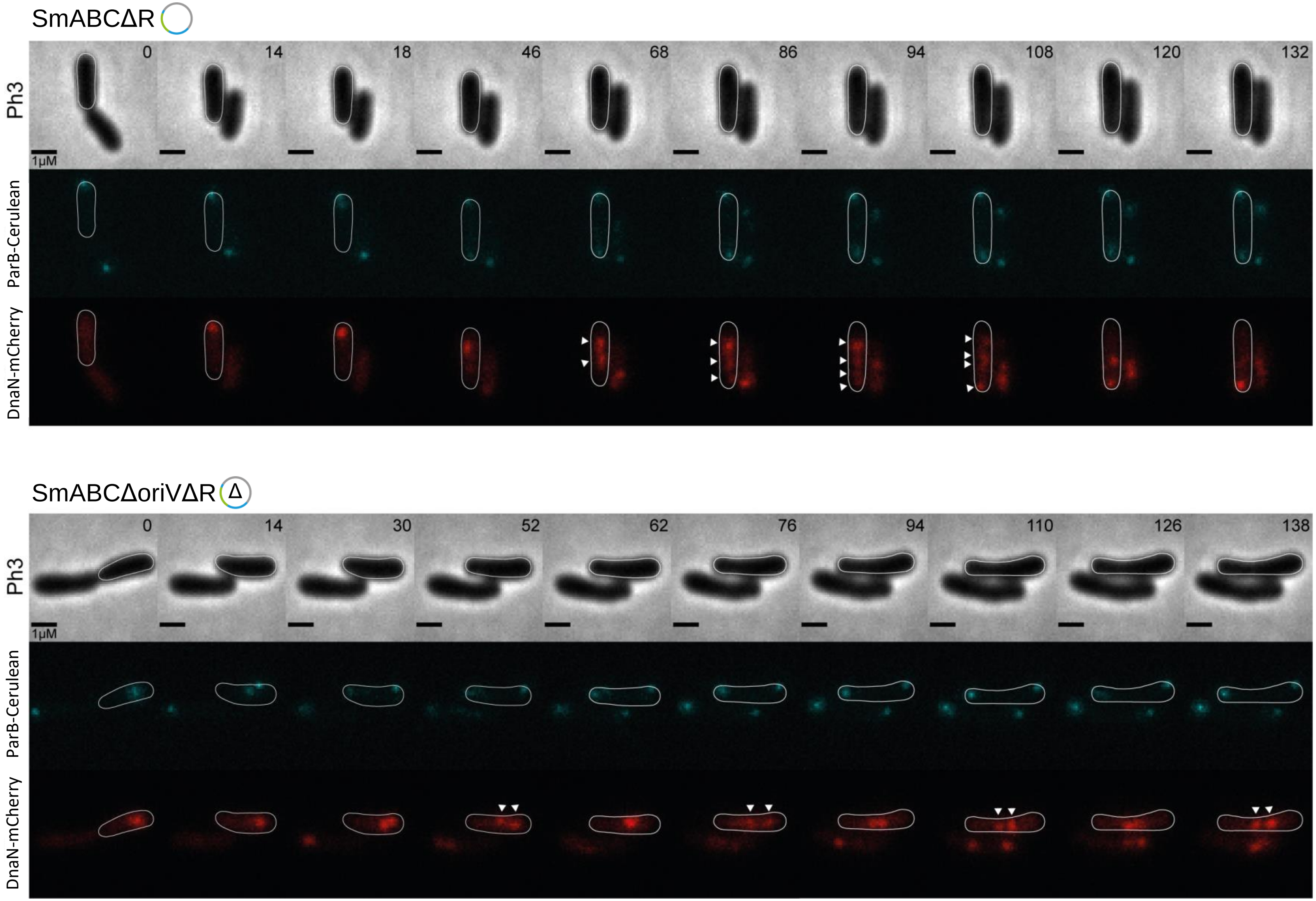

**S9 Fig. Replisome formation and dynamics in SmABCΔR vs SmABCΔoriVΔR.** For visualization and comparison of the intracellular replisome dynamics native *dnaN* in MWSm230 (SmABCΔR with *parB::cerulean*) and MWSm285 (SmABCΔoriVΔR with *parB::cerulean*) was replaced by a *dnaN-mCherry* translational fusion using pK19ms DnaN-mCherry. The resulting strains SmABCΔR-DnaN-mCh and SmABCΔoriVΔR-DnaN-mCh were analyzed in a 2 min interval time lapse series. The occurrence of 1-4 mCherry fluorescent foci in SmABCΔR and 1-2 foci in SmABCΔoriVΔR (indicated by white arrowheads) in selected time spots demonstrate a reduced replisome formation for SmABCΔoriVΔR suggesting loss of the replication initiation capacity of the secondary replicons' *repABC* modules deleted for *repC* and its intrinsic *oriV*. ParB-Cerulean foci visualize the location of the *oriC*(s) over the period of time.

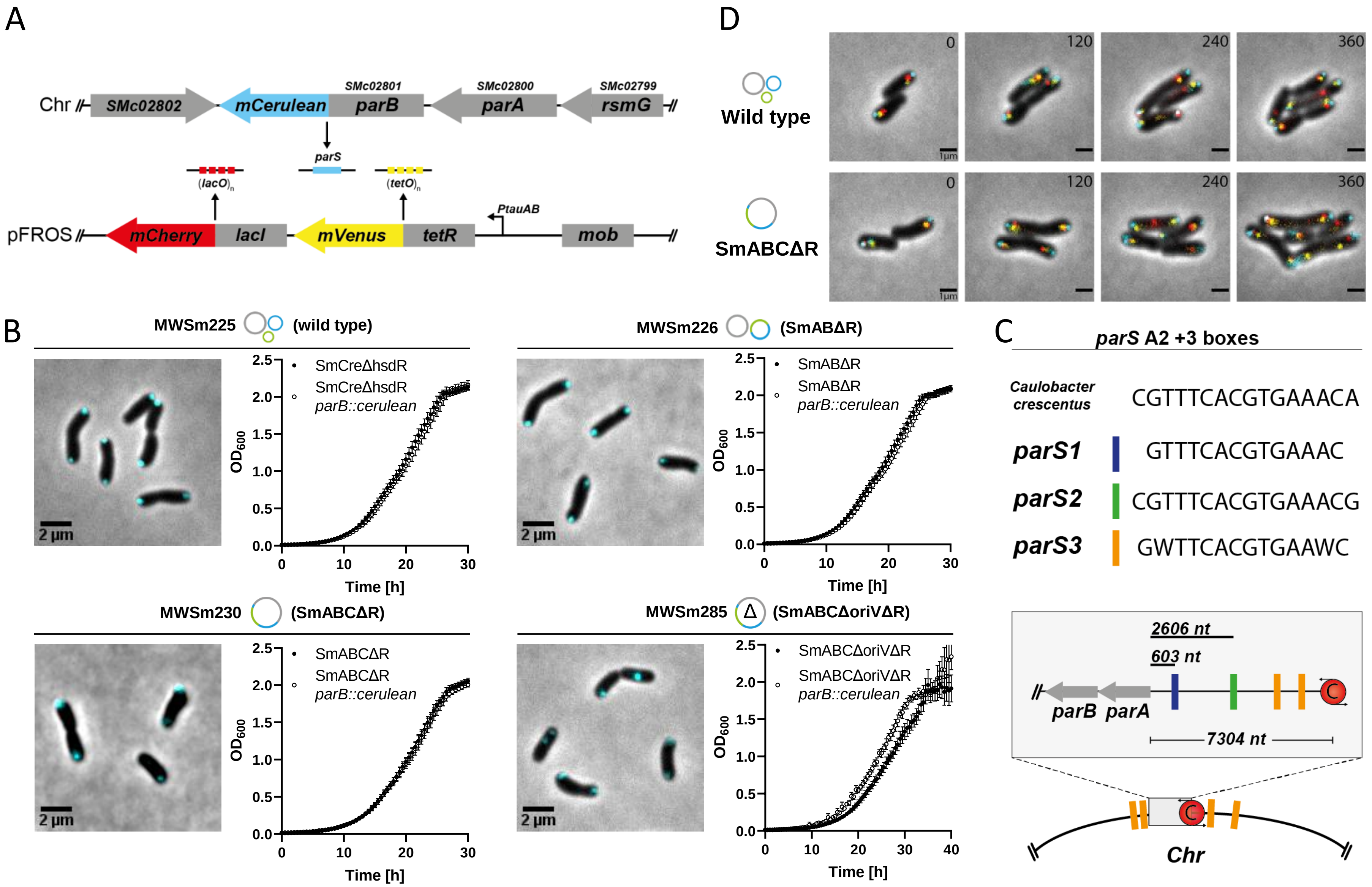

**S10 Fig. Establishment of a genome architecture independent labeling strategy. (A)** Components of the triple color system. A marker free in-frame fusion of *mcerulean* with *parB* (realized by integration of pMW198 via double homologous recombination) enables a constitutively expressed labeling of the chromosomal origin (*oriC)* in all relevant *S. meliloti* genome variants. The second part of the system (FROS) consists of a plasmid-based expression of *lacI-mCherry* and *tetR-mVenus* and enables the labeling of individual positions by genomic integration of related $tetO_{120}$ and $lacO_{120}$ arrays. Equipped with a mobilization site, the pFROS plasmid can be either transferred via conjugation or electroporation in all *S. meliloti* genome variants. **(B)** Validation of suitable ParB-Cerulean function in SmCreΔhsdR (wt), SmABΔR, SmABCΔR and SmABCΔoriVΔR. Snapshot images revealed a formation of either one or two fluorescence foci predominant at the pole indicating the system to constitute as proper *oriC* label. Growth curves of strains with *parB-cerulean* fusion show no negative effect on growth behavior when compared to direct precursor strains. **(C)** Putative *parS* A2+3 boxes for ParB binding on the *S. meliloti* chromosome. An incomplete A2+3 box is located 603 nt upstream of *parA* (*parS1*, GTTTCACGTGAAAC, position smc3647435), and a further complete palindrome (which is also covered by *parS1*) is situated 2606 nt upstream of *parA* (*parS2*, CGTTTCACGTGAAACG, genome position smc3649438). Interestingly, a degenerated variant (*parS3*, GWTTCACGTGAAWC; W= A or T) exclusively occurs on the chromosome and covers a 90 kb region including *parAB-oriC* (at genome positions smc1945, -54110, -3618431, -3634210, -3647435, -3649439, -3651011, -3652552). **D:** Fluorescence microscopy analysis of SmCreΔhsdR (wt) and SmABCΔR revealed a sufficient fluorescent signal intensity after 2h, 4h and 6h. This demonstrates that expression of the FROS reporter genes from pFROS can be achieved without any inducer (basal promotor activity coupled to plasmid copy number) over a long period of time which makes the system in particular attractive for long term analysis, like time lapse applications.

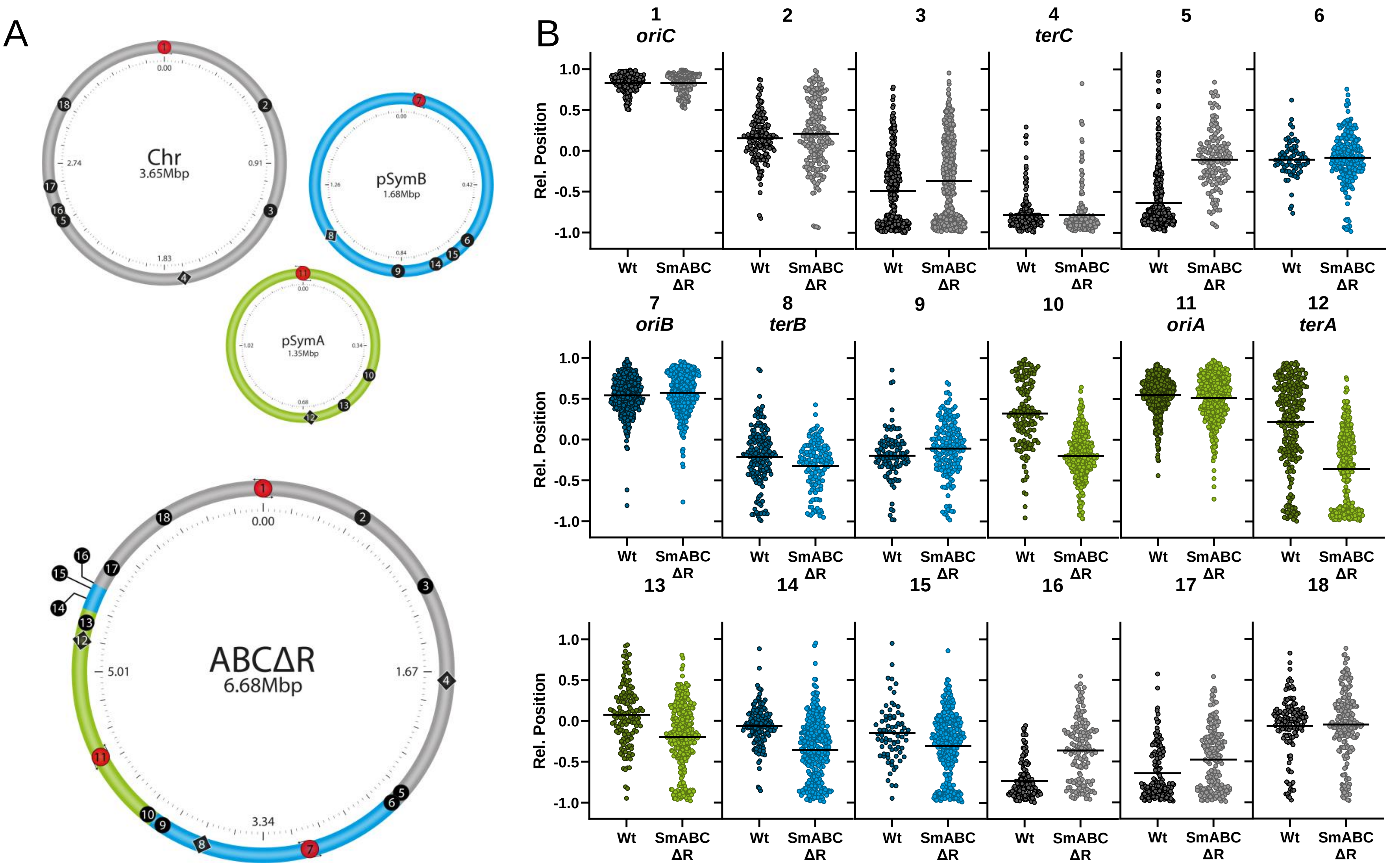

**S11 Fig. Genome organization in SmABCΔR compared to the wild type. (A)** Marker integration sites in genomic DNA of SmABCΔR and SmCreΔhsdR (wt) (for detailed information on the position of genomic integrations please refer to S1 Table) **(B)** Comparison of the fluorescent foci distribution in early $G_1$-phase cells (cell length max: 2.0 µm, single ParB-Cerulean focus) of SmCreΔhsdR (wt) and SmABCΔR. Old pole: 1, new pole: -1. Color code: chromosome (grey), pSymA (green), pSymB (blue). Origins are represented as red circles and terminus regions as black diamonds.

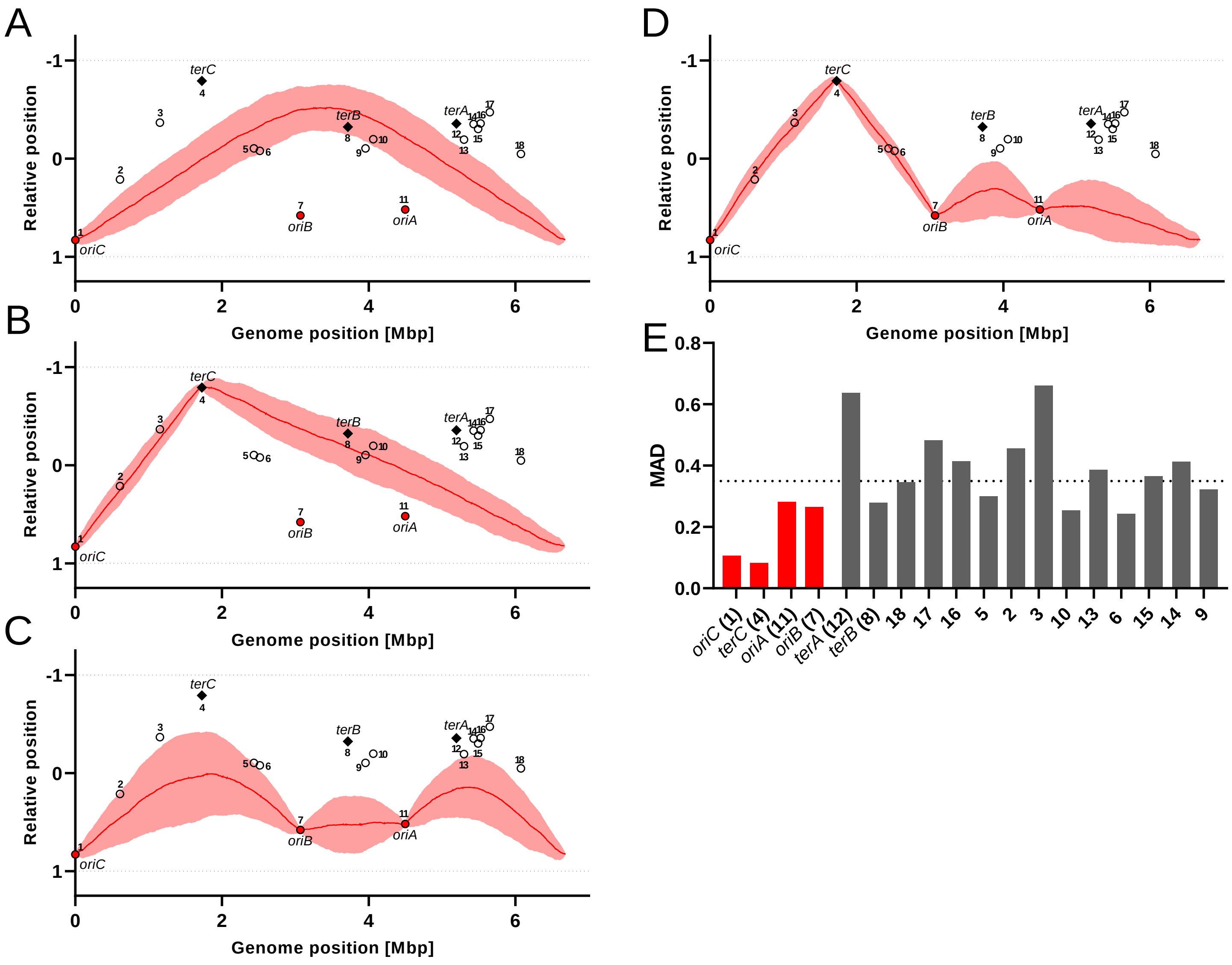

**S12 Fig. Modeling of spatial DNA arrangement in SmABCΔR.** Model prediction for the organization the ABC fusion in SmABC with **(A)** *oriC* **(B)** *oriC* and *terC* **(C)** *oriABC* **(D)** *oriABC* and *terC* as fixpoints at the mean experimental position. Shown are the normalized positions in the cell (old pole: 1; new pole: -1) as a function of the position on the genomic map [Mbp]. Experimental data of origins are indicated by red circles, terminus regions by black diamonds and remaining marker positions with non-filled circles. The red line depicts the model results averaged over 200 cells. Shaded areas represent the standard deviations. For each simulation, a cell size of 1800 nm and a loop size of 1298 bp (DNA within a "blob") was assumed. **(E)** Bar chart of the dispersion of the experimental marker data for SmABCΔR. The statistical dispersion is measured via the mean absolute deviation (MAD). The horizontal dashed line indicates the average dispersion (0.35) over all marker.

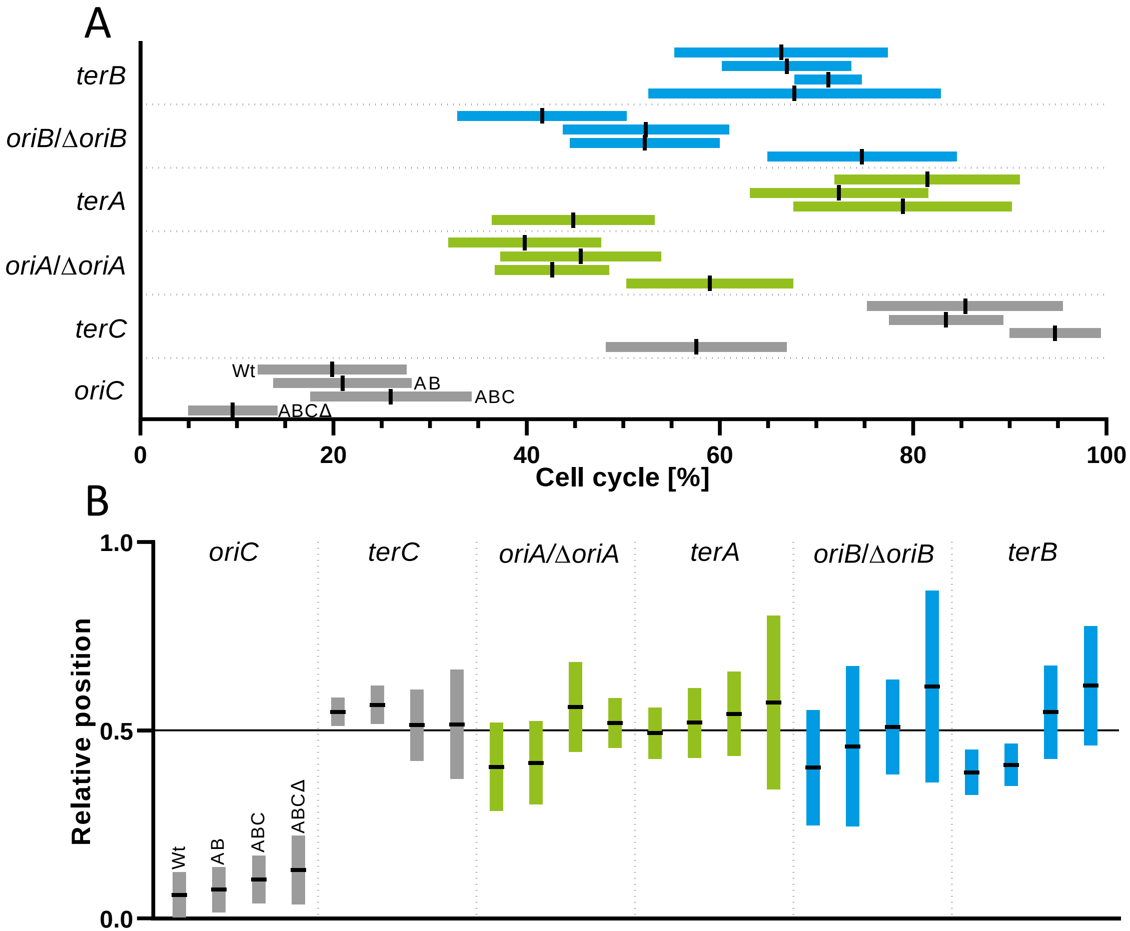

**S13 Fig. Cumulating time-lapse data of SmCreΔhsdR (wt), SmAB (AB), SmABC (ABC) and SmABCΔoriVΔR (ABCΔ) daughter cells. (A)** Timepoint of visual segregation of origin and terminus foci. Bar chart depicting the timepoint of visual foci segregation within the *S. meliloti* cell cycle normalized to 100 % (0 % - cell cycle start, 100 % - cell cycle completed). Bars are indicative for the standard deviation (SD) with the mean as black centered lines. **(B)** Localization of origin and terminus foci segregation. Bar chart depicting the longitudinal position of visual foci segregation within the *S. meliloti* cell normalized to 1 (0 - old pole, 1 – new pole). Bars are indicative for the standard deviation (SD) with the mean as black centered lines. Color code: chromosome (grey), pSymA (green), pSymB (blue).

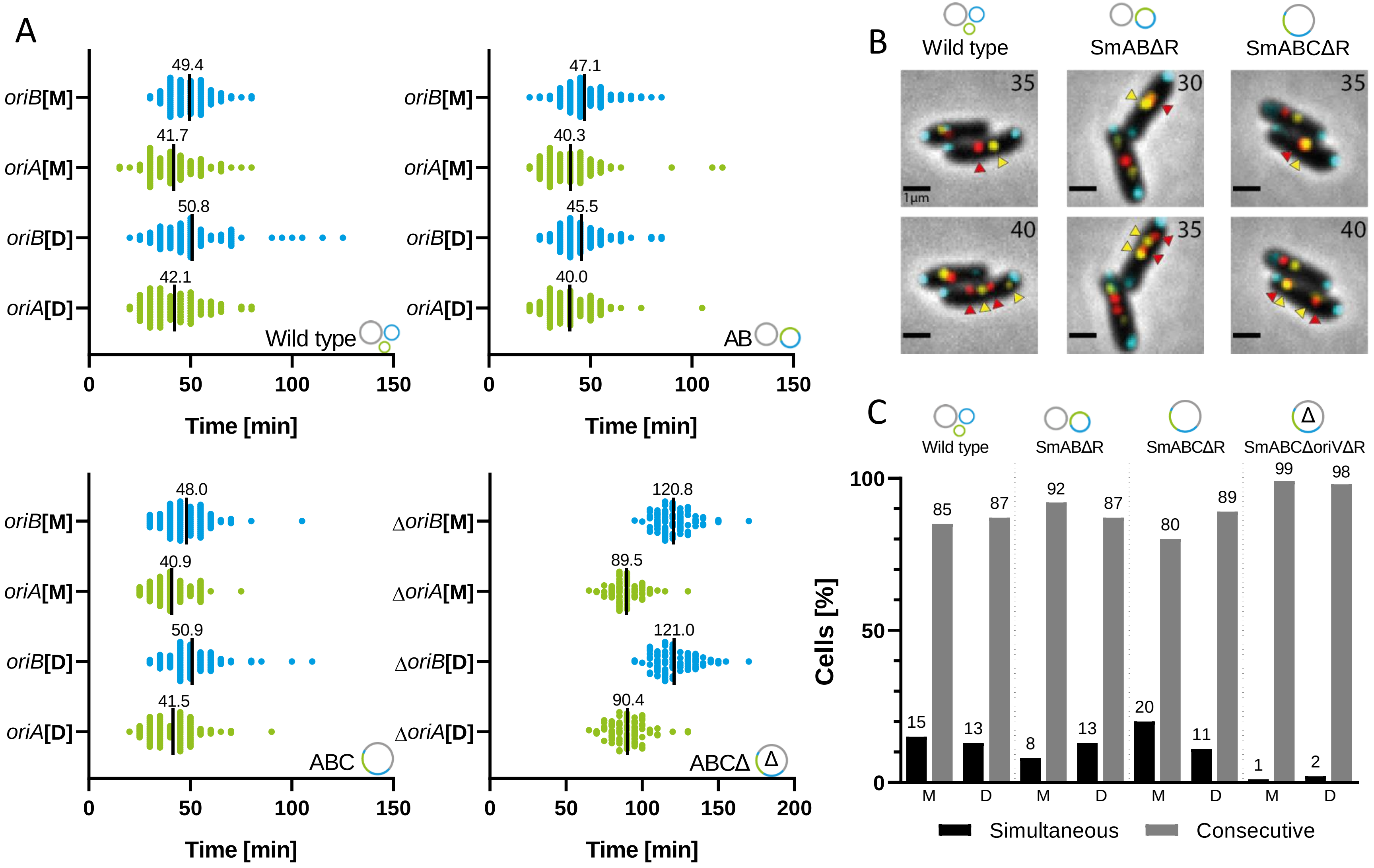

**S14 Fig. *oriA* and *oriB* segregation. (A)** Scatter plot with timing of the *oriA* and *oriB* segregation events post *oriC* segregation in the SmCreΔhsdR (wild type) , SmABΔR (AB), SmABCΔR (ABC) and SmABCΔoriVΔR (ABCΔ) subdivided in mother and dauther cells (n=100 each). Black lines depict the mean values of the plot data. **(B)** Microscopy images of cells with simultaneously segregating *oriA*/Δ*oriA* and *oriB*/Δ*oriB* foci within the chosen time lapse settings (5 minutes intervals between the individual images). Arrowheads depict the localization of *oriA*/Δ*oriA* (yellow) and *oriB*/Δ*oriB* (red) before (upper panel) and after (lower panel) the segregation event. Scale bar: 1 µm. **(C)** Percentage of cells per strain (n=200 each) with simultaneous and consecutive occurring segregation event. Color code: chromosome (grey), pSymA (green), pSymB (blue).

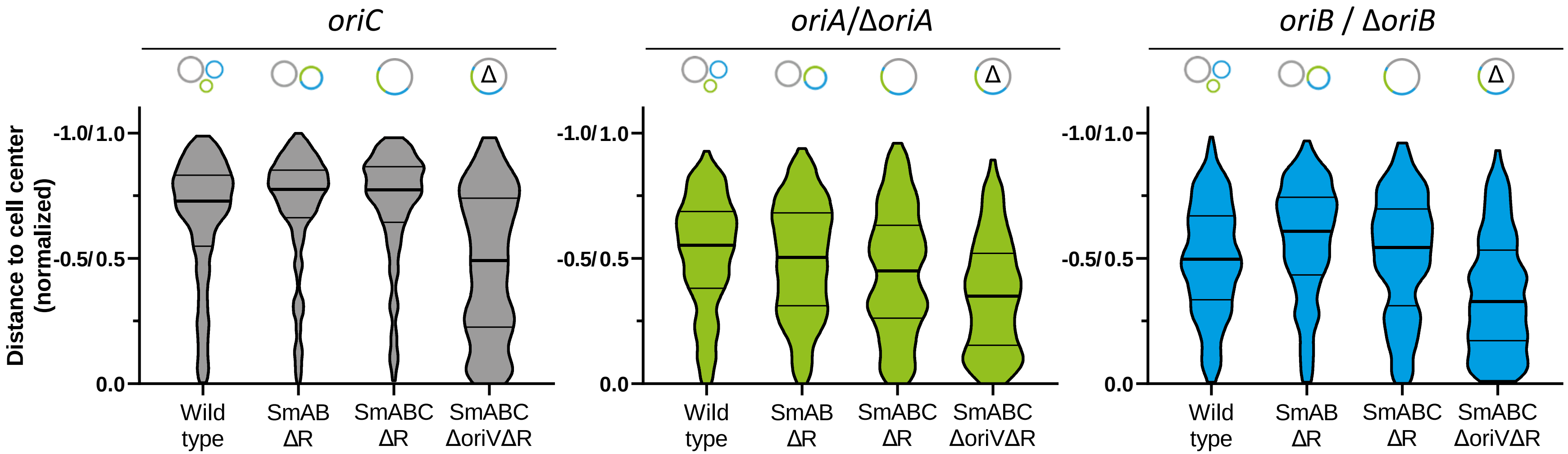

**S15 Fig. Comparison of *oriC*, *oriA*/Δ*oriA* and *oriB*/Δ*oriB* distribution in SmCreΔhsdR (wt), SmABΔR, SmABCΔR and SmABCΔoriVΔR.** Violin plots depict the normalized distance of the individual origin regions to the cell center (cell poles: 1 and -1, cell center: 0). Plots are based on snapshot analysis of $G_1$-phase cells filtered by size with a cut-off of 2.0 µm. Wild type (n=639), SmABΔR (n=460), SmABCΔR (n=405) and SmABCΔoriVΔR (n=246). Color code: chromosome (grey), pSymA (green), pSymB (blue).

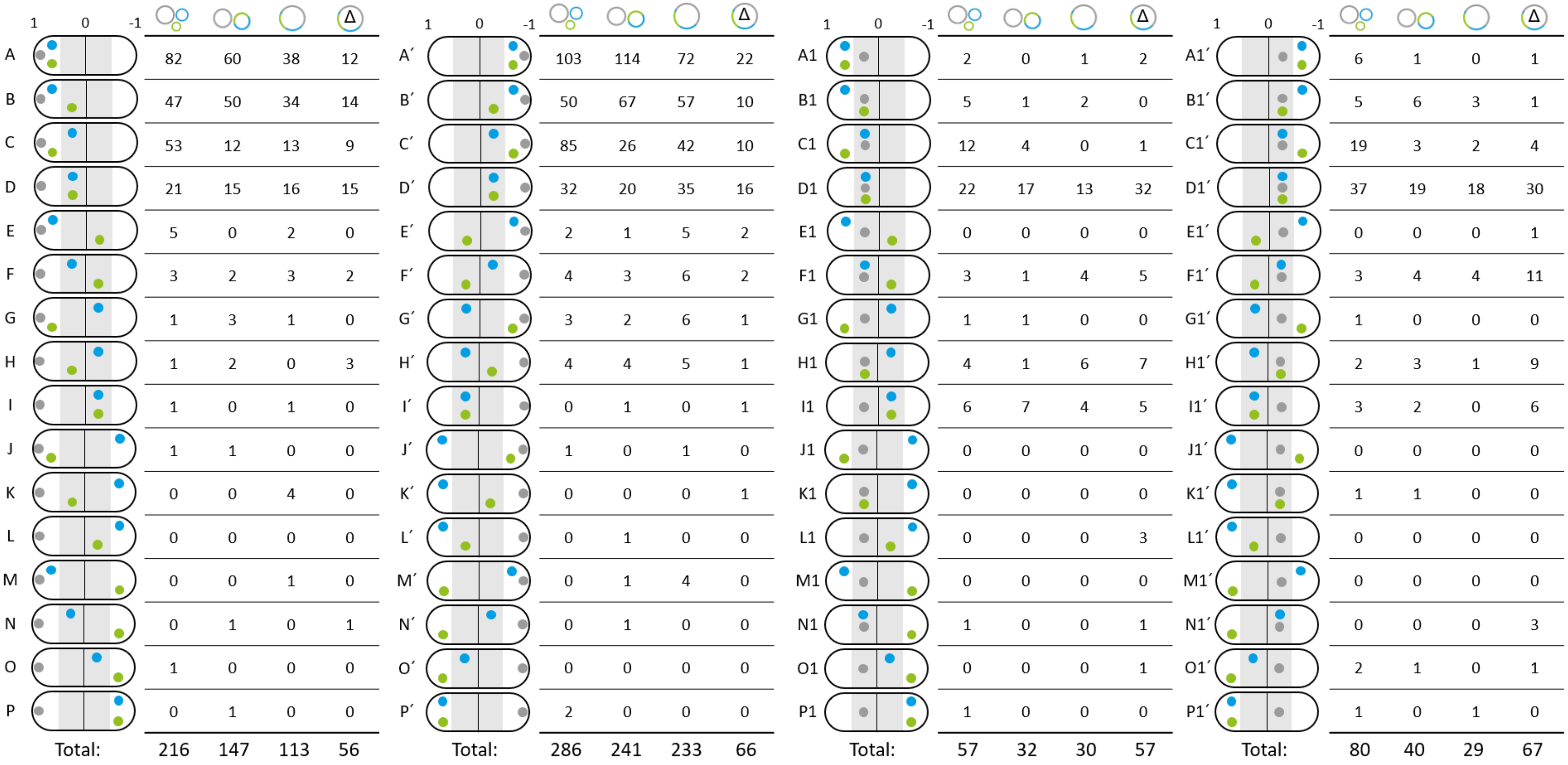

| | | | | Δ |
|---|---|---|---|---|
| A | 82 | 60 | 38 | 12 |
| B | 47 | 50 | 34 | 14 |
| C | 53 | 12 | 13 | 9 |
| D | 21 | 15 | 16 | 15 |
| E | 5 | 0 | 2 | 0 |
| F | 3 | 2 | 3 | 2 |
| G | 1 | 3 | 1 | 0 |
| H | 1 | 2 | 0 | 3 |
| I | 1 | 0 | 1 | 0 |
| J | 1 | 1 | 0 | 0 |
| K | 0 | 0 | 4 | 0 |
| L | 0 | 0 | 0 | 0 |
| M | 0 | 0 | 1 | 0 |
| N | 0 | 1 | 0 | 1 |
| O | 1 | 0 | 0 | 0 |
| P | 0 | 1 | 0 | 0 |
| Total: | 216 | 147 | 113 | 56 |

| | | | | Δ |
|---|---|---|---|---|
| A′ | 103 | 114 | 72 | 22 |
| B′ | 50 | 67 | 57 | 10 |
| C′ | 85 | 26 | 42 | 10 |
| D′ | 32 | 20 | 35 | 16 |
| E′ | 2 | 1 | 5 | 2 |
| F′ | 4 | 3 | 6 | 2 |
| G′ | 3 | 2 | 6 | 1 |
| H′ | 4 | 4 | 5 | 1 |
| I′ | 0 | 1 | 0 | 1 |
| J′ | 1 | 0 | 1 | 0 |
| K′ | 0 | 0 | 0 | 1 |
| L′ | 0 | 1 | 0 | 0 |
| M′ | 0 | 1 | 4 | 0 |
| N′ | 0 | 1 | 0 | 0 |
| O′ | 0 | 0 | 0 | 0 |
| P′ | 2 | 0 | 0 | 0 |
| Total: | 286 | 241 | 233 | 66 |

| | | | | Δ |
|---|---|---|---|---|
| A1 | 2 | 0 | 1 | 2 |
| B1 | 5 | 1 | 2 | 0 |
| C1 | 12 | 4 | 0 | 1 |
| D1 | 22 | 17 | 13 | 32 |
| E1 | 0 | 0 | 0 | 0 |
| F1 | 3 | 1 | 4 | 5 |
| G1 | 1 | 1 | 0 | 0 |
| H1 | 4 | 1 | 6 | 7 |
| I1 | 6 | 7 | 4 | 5 |
| J1 | 0 | 0 | 0 | 0 |
| K1 | 0 | 0 | 0 | 0 |
| L1 | 0 | 0 | 0 | 3 |
| M1 | 0 | 0 | 0 | 0 |
| N1 | 1 | 0 | 0 | 1 |
| O1 | 0 | 0 | 0 | 1 |
| P1 | 1 | 0 | 0 | 0 |
| Total: | 57 | 32 | 30 | 57 |

| | | | | Δ |
|---|---|---|---|---|
| A1′ | 6 | 1 | 0 | 1 |
| B1′ | 5 | 6 | 3 | 1 |
| C1′ | 19 | 3 | 2 | 4 |
| D1′ | 37 | 19 | 18 | 30 |
| E1′ | 0 | 0 | 0 | 1 |
| F1′ | 3 | 4 | 4 | 11 |
| G1′ | 1 | 0 | 0 | 0 |
| H1′ | 2 | 3 | 1 | 9 |
| I1′ | 3 | 2 | 0 | 6 |
| J1′ | 0 | 0 | 0 | 0 |
| K1′ | 1 | 1 | 0 | 0 |
| L1′ | 0 | 0 | 0 | 0 |
| M1′ | 0 | 0 | 0 | 0 |
| N1′ | 0 | 0 | 0 | 3 |
| O1′ | 2 | 1 | 0 | 1 |
| P1′ | 1 | 0 | 1 | 0 |
| Total: | 80 | 40 | 29 | 67 |

**S16 Fig. Origin co-localization scheme for SmCreΔhsdR (wt), SmABΔR, SmABCΔR and SmABCΔoriVΔR.** The scheme depict the distribution pattern of *oriA/ΔoriA* (green spots) *oriB*/Δ*oriB* (blue spots) and *oriC* (grey spots) in relation to each other. Therefore, cells were divided into four compartments (1 to 0.5, 0.5 to 0, 0 to -0.5 and -0.5 to -1). Cells analyzed: SmCreΔhsdR (wt) (n=639), SmABΔR (n=460), SmABCΔR (n=405) and SmABCΔoriVΔR (n=246).

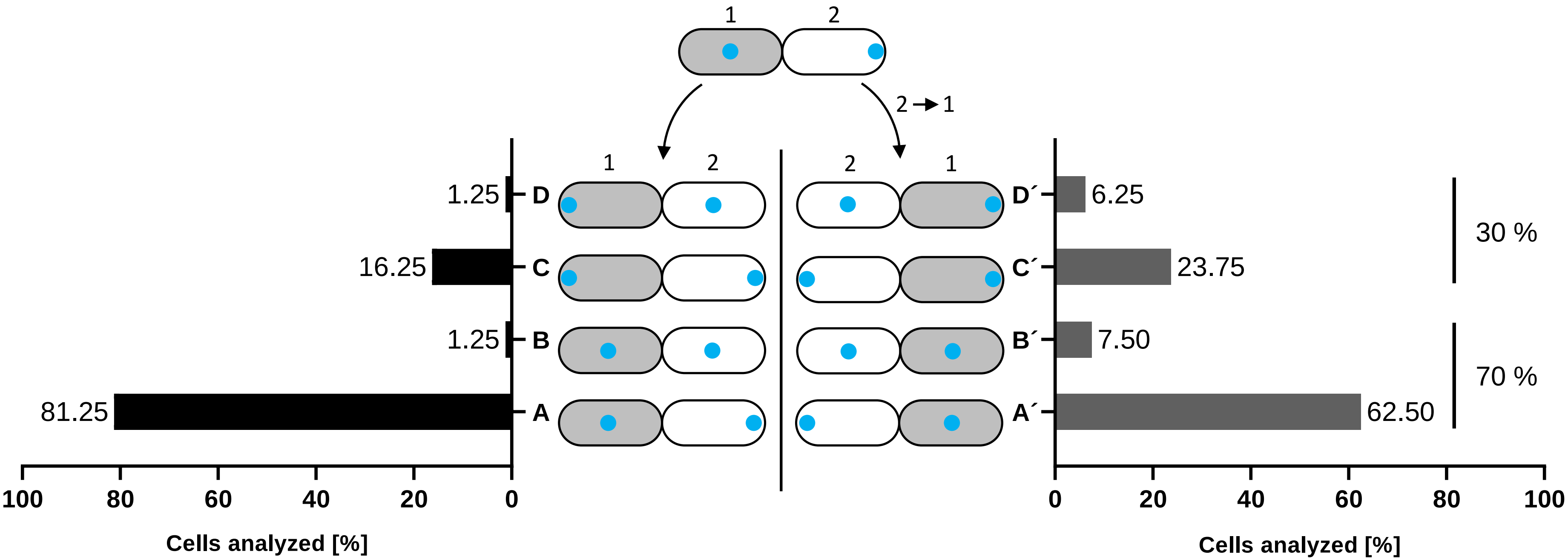

**S17 Fig. *oriC* localization pattern in SmABCΔoriVΔR cells after one cell division cycle .** Depicted are the percentage of the predominant *oriC* positions (blue spot) in the next generation of sibling cells emerging from either sibling 1 (grey) or sibling 2 (white). Cells analyzed: n=80.

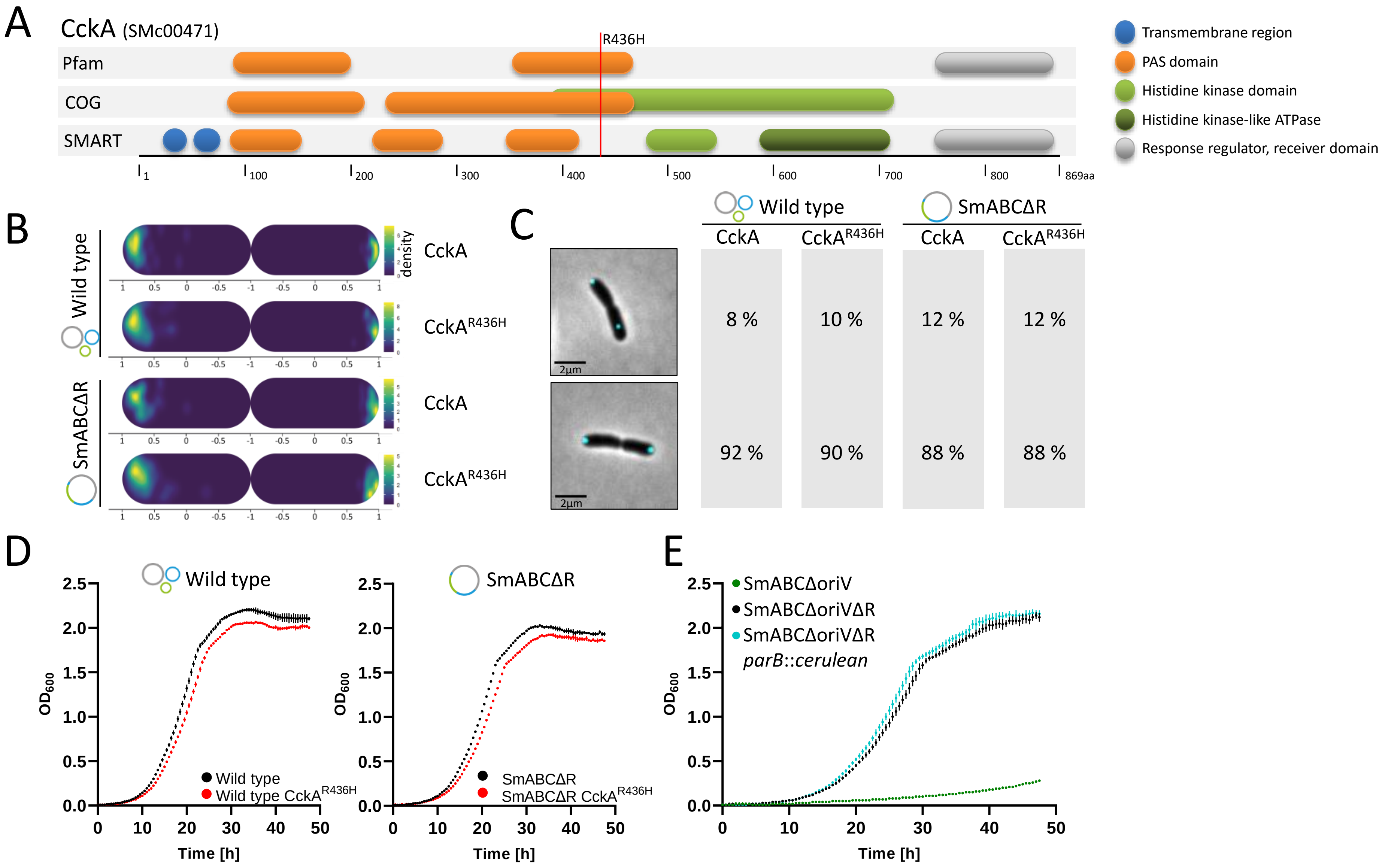

**S18 Fig. Analysis of CckA$^{R436H}$ in *S. meliloti* wild type and genome fusion strains. (A)** Schematic illustration of functional domains of CckA predicted by Pfam, COG and SMART via NCBI CDD (Marchler-Bauer et al., 2011, 2015 and 1017). The red line depicts the position of the missense mutation R436H. **(B)** Densitiy map of *oriC* (ParB-cerulean) localization in predivisional sibling cells of SmCreΔhsdR (wt) and SmABCΔR with original and mutagenized *cckA*. **(C)** Percentage of predividional sibling cells of SmCreΔhsdR (wt) and SmABCΔR with either both *oriCs* located at the poles (lower picture and values) or with at least one *oriC* focus located in the midcell area (upper picture and values). **(D)** Growth curves of SmCreΔhsdR (wt) and SmABCΔR with original CckA compared to the respective strains with CckA$^{R436H}$. **(E)** Growth comparison of SmABCΔoriVΔR with the precursor strain SmABCΔoriV and SmABCΔoriVΔR with *parB::cerulean* fusion.

**S1 Table. Underlying data of S11B Fig.** Mean and standard deviation (SD) of the relative longitudinal positions of fluorescent foci within analyzed cells of SmABCΔR and the wild type. Old cell pole at 1, new cell pole at -1.

| | | Wild type | | | | SmABCΔR | | | |
|---|---|---|---|---|---|---|---|---|---|
| **Loci no.** | **Location** | **Genome pos. (Mbp)** | **Mean rel. position** | **SD** | **n** | **Genome pos. (Mbp)** | **Mean rel. position** | **SD** | **n** |
| 1 | Chromosome (*oriC*) | 0.00 | 0.83 | 0.11 | 206 | 0.00 | 0.83 | 0.12 | 217 |
| 2 | Chromosome | 0.61 | 0.15 | 0.28 | 154 | 0.61 | 0.21 | 0.41 | 211 |
| 3 | Chromosome | 1.15 | -0.49 | 0.42 | 338 | 1.15 | -0.37 | 0.49 | 427 |
| 4 | Chromosome (*terC*) | 1.72 | -0.79 | 0.20 | 206 | 1.72 | -0.79 | 0.26 | 217 |
| 5 | Chromosome | 2.44 | -0.63 | 0.34 | 409 | 2.44 | -0.10 | 0.35 | 152 |
| 6 | pSymB | 0.61 | -0.11 | 0.22 | 76 | 2.52 | -0.08 | 0.30 | 238 |
| 7 | pSymB (*oriB*) | 0.05 | 0.54 | 0.22 | 466 | 3.07 | 0.58 | 0.24 | 405 |
| 8 | pSymB (*terB*) | 1.10 | -0.21 | 0.35 | 170 | 3.71 | -0.32 | 0.30 | 145 |
| 9 | pSymB | 0.85 | -0.19 | 0.30 | 108 | 3.96 | -0.10 | 0.34 | 191 |
| 10 | pSymA | 0.43 | 0.32 | 0.41 | 152 | 4.06 | -0.20 | 0.29 | 316 |
| 11 | pSymA (*oriA*) | 1.35 | 0.55 | 0.22 | 466 | 4.50 | 0.52 | 0.28 | 405 |
| 12 | pSymA (*terA*) | 0.65 | 0.22 | 0.54 | 276 | 5.19 | -0.36 | 0.45 | 341 |
| 13 | pSymA | 0.55 | 0.08 | 0.37 | 140 | 5.30 | -0.20 | 0.40 | 293 |
| 14 | pSymB | 0.73 | -0.06 | 0.24 | 150 | 5.43 | -0.35 | 0.38 | 320 |
| 15 | pSymB | 0.67 | -0.15 | 0.34 | 76 | 5.49 | -0.30 | 0.39 | 329 |
| 16 | Chromosome | 2.49 | -0.73 | 0.23 | 137 | 5.53 | -0.36 | 0.38 | 152 |
| 17 | Chromosome | 2.62 | -0.64 | 0.32 | 161 | 5.65 | -0.47 | 0.37 | 180 |
| 18 | Chromosome | 3.05 | -0.06 | 0.35 | 136 | 6.08 | -0.05 | 0.41 | 203 |

**S2 Table. Timepoint and Localization of origin and terminus foci segregation in SmCreΔhsdR (wt), SmAB, SmABC and SmABCΔoriVΔR mother cells [M]. (A)** Corresponding values to Fig. 5A (main manuscript). Mean and standard deviation (SD) of corresponding origin / *repAB*Δ*C* (Δ*oriA* or Δ*oriB*) and terminus foci segregation normalized to a cell cycle duration of 100 % (0 % - cell cycle start, 100 % - cell cycle completed) in the cells analyzed (n). **(B)** Corresponding values to Fig.5B (main manuscript). Mean and standard deviation (SD) of the relative longitudinal position of corresponding origin/ *repAB*Δ*C* (Δ*oriA* or Δ*oriB*) and terminus foci segregation normalized to 1 (0 - old pole (OP), 1 – new pole (NP)) within the cells analyzed (n).

A

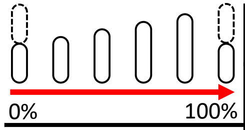

| | ***oriC*** | | ***terC*** | | ***oriA/ΔoriA*** | | ***terA*** | | ***oriB/ΔoriB*** | | ***terB*** | |
|---|---|---|---|---|---|---|---|---|---|---|---|---|
| | **Mean ± SD** | **n** | **Mean ± SD** | **n** | **Mean ± SD** | **n** | **Mean ± SD** | **n** | **Mean ± SD** | **n** | **Mean ± SD** | **n** |
| **Wild type [M]** | 8.9 ± 3.5 | 20 | 89.5 ± 8.7 | 5 | 34.7 ± 7.3 | 10 | 73.1 ± 11.2 | 5 | 38.7 ± 10.1 | 10 | 59.8 ± 14.8 | 5 |
| **SmABΔR [M]** | 14.9 ± 5.5 | 20 | 83.6 ± 5.0 | 5 | 40.7 ± 6.9 | 10 | 64.8 ± 10.3 | 5 | 43.0 ± 8.8 | 10 | 70.5 ± 9.2 | 5 |
| **SmABCΔR [M]** | 15.7 ± 6.5 | 20 | 93.5 ± 7.4 | 5 | 35.4 ± 6.4 | 10 | 61.4 ± 3.2 | 5 | 45.5 ± 6.5 | 10 | 65.8 ± 5.4 | 5 |
| **SmABCΔoriVΔR [M]** | 9.5 ± 3.1 | 20 | 43.1 ± 10.0 | 5 | 56.8 ± 7.3 | 10 | 40.5 ± 5.3 | 5 | 71.9 ± 10.3 | 10 | 66.7 ± 10.0 | 5 |

B

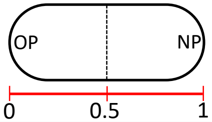

| | ***oriC*** | | ***terC*** | | ***oriA/ΔoriA*** | | ***terA*** | | ***oriB/ΔoriB*** | | ***terB*** | |
|---|---|---|---|---|---|---|---|---|---|---|---|---|
| | **Mean ± SD** | **n** | **Mean ± SD** | **n** | **Mean ± SD** | **n** | **Mean ± SD** | **n** | **Mean ± SD** | **n** | **Mean ± SD** | **n** |
| **Wild type [M]** | 0.09 ± 0.05 | 20 | 0.54 ± 0.11 | 5 | 0.43 ± 0.17 | 10 | 0.51 ± 0.13 | 5 | 0.48 ± 0.11 | 10 | 0.48 ± 0.24 | 5 |
| **SmABΔR [M]** | 0.07 ± 0.05 | 20 | 0.56 ± 0.09 | 5 | 0.48 ± 0.11 | 10 | 0.42 ± 0.12 | 5 | 0.46 ± 0.16 | 10 | 0.45 ± 0.04 | 5 |
| **SmABCΔR [M]** | 0.12 ± 0.11 | 20 | 0.58 ± 0.06 | 5 | 0.53 ± 0.14 | 10 | 0.51 ± 0.10 | 5 | 0.49 ± 0.13 | 10 | 0.45 ± 0.03 | 5 |
| **SmABCΔoriVΔR [M]** | 0.41 ± 0.14 | 20 | 0.59 ± 0.14 | 5 | 0.51 ± 0.11 | 10 | 0.50 ± 0.18 | 5 | 0.71 ± 0.20 | 10 | 0.55 ± 0.14 | 5 |

**S3 Table. Values corresponding to S13A Fig. daughter cells [D]. (A)** Mean and standard deviation (SD) of origin / *repABΔC* (Δ*oriA* or Δ*oriB*) and terminus foci segregation normalized to a cell cycle duration of 100 % (0 % - cell cycle start, 100 % - cell cycle completed) in the daughter cells [D] analyzed (n). **(B)** Mean and standard deviation (SD) of the relative longitudinal position of corresponding origin/ *repABΔC* (Δ*oriA* or Δ*oriB*) and terminus foci segregation normalized to 1 (0 - old pole, 1 – new pole) within the daughter cells analyzed (n).

A

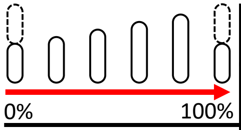

| | ***oriC*** | | ***terC*** | | ***oriA/ΔoriA*** | | ***terA*** | | ***oriB/ΔoriB*** | | ***terB*** | |
|---|---|---|---|---|---|---|---|---|---|---|---|---|
| | **Mean ± SD** | **n** | **Mean ± SD** | **n** | **Mean ± SD** | **n** | **Mean ± SD** | **n** | **Mean ± SD** | **n** | **Mean ± SD** | **n** |
| **Wild type [D]** | 19.9 ± 7.7 | 18 | 85.4 ± 10.1 | 5 | 39.8 ± 7.9 | 8 | 81.5 ± 9.6 | 5 | 41.6 ± 8.8 | 8 | 66.3 ± 11.0 | 5 |
| **SmABΔR [D]** | 20.9 ± 7.2 | 20 | 83.4 ± 5.9 | 5 | 45.6 ± 8.3 | 10 | 72.3 ± 9.2 | 5 | 52.3 ± 8.6 | 10 | 66.9 ± 6.7 | 5 |
| **SmABCΔR [D]** | 25.9 ± 8.4 | 20 | 94.7 ± 4.8 | 5 | 42.6 ± 5.9 | 10 | 78.9 ± 11.3 | 5 | 52.2 ± 7.8 | 10 | 71.2 ± 3.5 | 5 |
| **SmABCΔoriVΔR [D]** | 9.6 ± 4.6 | 20 | 57.6 ± 9.4 | 5 | 59.0 ± 8.6 | 10 | 44.8 ± 8.5 | 5 | 74.7 ± 9.8 | 10 | 67.7 ± 15.1 | 5 |

B

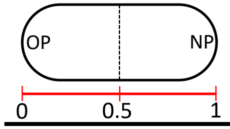

| | ***oriC*** | | ***terC*** | | ***oriA/ΔoriA*** | | ***terA*** | | ***oriB/ΔoriB*** | | ***terB*** | |
|---|---|---|---|---|---|---|---|---|---|---|---|---|
| | **Mean ± SD** | **n** | **Mean ± SD** | **n** | **Mean ± SD** | **n** | **Mean ± SD** | **n** | **Mean ± SD** | **n** | **Mean ± SD** | **n** |
| **Wild type [D]** | 0.06 ± 0.06 | 20 | 0.55 ± 0.04 | 5 | 0.40 ± 0.12 | 10 | 0.49 ± 0.07 | 5 | 0.40 ± 0.15 | 10 | 0.39 ± 0.06 | 5 |
| **SmABΔR [D]** | 0.08 ± 0.06 | 20 | 0.57 ± 0.05 | 5 | 0.41 ± 0.11 | 10 | 0.52 ± 0.09 | 5 | 0.46 ± 0.21 | 10 | 0.41 ± 0.06 | 5 |
| **SmABCΔR [D]** | 0.10 ± 0.06 | 20 | 0.51 ± 0.09 | 5 | 0.56 ± 0.12 | 10 | 0.54 ± 0.11 | 5 | 0.51 ± 0.13 | 10 | 0.55 ± 0.12 | 5 |
| **SmABCΔoriVΔR [D]** | 0.13 ± 0.09 | 20 | 0.52 ± 0.15 | 5 | 0.52 ± 0.07 | 10 | 0.57 ± 0.23 | 5 | 0.62 ± 0.26 | 10 | 0.62 ± 0.16 | 5 |

**S4 Table. Coding sequences that were removed or truncated upon the replicon fusion procedure.** For details on the individual fusion site (FS) please refer to S1 and S2 Fig.

| Strain | Annotation | Fusion | Product | Type of alteration |
|---|---|---|---|---|
| **SmAB** | | | | |
| | SMa0854 (*nodG*) | FS2 (A+B) | NodG 3-oxoacyl-(acyl carrier protein) reductase | Truncated (C-terminal 89 bp missing) |
| | SMb21225 | FS2 (A+B) | Inositol-phosphate phosphatase | Truncated (N-terminal 4 bp missing) |
| **SmABC** | | | | |
| | SMc01552 | FS4 (C+B) | Hypothetical transmembrane protein | Removed |
| | SMc01553 | FS4 (C+B) | Acyl carrier protein | Removed |
| | SMc01554 | FS4 (C+B) | Conserved hypothetical protein | Removed |
| | SMb21013 | FS3 (C+B) | Hypothetical protein | Removed |
| | SMb21014 | FS3 (C+B) | Putative protein | Truncated (N-terminal 122 bp left) |

**S5 Table. Single nucleotide variations associated with the fusion procedure.** Reference positions refer to the genome of *S. meliloti* SmCreΔhsdR which was used as the reference for the basic variant detection analysis CDS: coding sequence, * paralogue, MBOAT: membrane bound O-acyl transferase

| Strain | Mapping | Reference Pos. | Reference | Allele | Position | Fusion site | Annotation | Product | Mutation type |
|---|---|---|---|---|---|---|---|---|---|
| **SmAB** | | | | | | | | | |
| | pSymB | 789713 | A | T | CDS | FS2 (A+B) | SMb21225 | Inositol-phosphate phosphatase | Missense mutation (Ile → Phe) |
| **SmABC** | | | | | | | | | |
| | Chr | 2488149 | C | T | CDS | FS4 (C+B) | SMc01551 (*algI**) | Putative MBOAT | Silent mutation (Leu → Leu) |
| | Chr | 2488158 | A | C | CDS | FS4 (C+B) | SMc01551 (*algI**) | Putative MBOAT | Missense mutation (Glu → Asp) |
| | Chr | 2488163 | G | A | CDS | FS4 (C+B) | SMc01551 (*algI**) | Putative MBOAT | Missense mutation (Ser → Asn) |
| | pSymB | 633549 | A | G | CDS | FS4 (C+B) | SMb20842 | Hypothetical transmembrane protein | Silent mutation (Gly → Gly) |
| | pSymB | 635961 | C | T | CDS | FS4 (C+B) | SMb20843 (*algI*) | Predicted membrane protein, MBOAT | Silent mutation (Ser → Ser) |
| | pSymB | 635979 | T | C | CDS | FS4 (C+B) | SMb20843 (*algI*) | Predicted membrane protein, MBOAT | Silent mutation (Gln → Gln) |
| | pSymB | 635982 | G | A | CDS | FS4 (C+B) | SMb20843 (*algI*) | Predicted membrane protein, MBOAT | Silent mutation (Gly → Gly) |
| | pSymB | 635988 | T | C | CDS | FS4 (C+B) | SMb20843 (*algI*) | Predicted membrane protein, MBOAT | Silent mutation (Gly → Gly) |
| | pSymB | 789713 | A | T | CDS | FS2 (A+B) | SMb21225 | Inositol-phosphate phosphatase | Missense mutation (Ile → Phe) |

**S6 Table. Single nucleotide variations and nucleotide deletions in *S. meliloti* genome fusion strains.** Reference positions refer to the genome of *S. meliloti* SmCreΔhsdR which was used as the reference for the basic variant detection analysis CDS: coding sequence, IGR: intergenic region.

| Strain | Mapping | Reference Pos. | Reference | Allele | Position | Annotation | Product | Mutation type |
|---|---|---|---|---|---|---|---|---|
| **SmAB** | | | | | | | | |
| | pSymB | 842124 | C | - | CDS | SMb21268 | Putative ABC transporter | Deletion |
| **SmABC** | | | | | | | | |
| | pSymB | 279482 | G | T | CDS | SMb20275 | Conserved hypothetical protein | Missense mutation (Ala → Glu) |
| | pSymB | 842124 | C | - | CDS | SMb21268 | Putative ABC transporter | Deletion |
| **SmABCΔoriV** | | | | | | | | |
| | pSymA | 81732 | G | - | IGR | SMa0148 - SMa0149 | - | Deletion |
| | pSymB | 279482 | G | T | CDS | SMb20275 | Conserved hypothetical protein | Missense mutation (Ala → Glu) |
| | pSymB | 842124 | C | - | CDS | SMb21268 | Putative ABC transporter | Deletion |
| **SmABΔR** | | | | | | | | |
| | pSymB | 842124 | C | - | CDS | SMb21268 | Putative ABC transporter | Deletion |
| **SmABCΔR** | | | | | | | | |
| | Chr | 18124 | C | A | CDS | SMc02776 | Altronate hydrolase, UxaA family | Missense mutation (Thr → Asn) |
| | pSymB | 842124 | C | - | CDS | SMb21268 | Putative ABC transporter | Deletion |
| **SmABCΔoriVΔR** | | | | | | | | |
| | Chr | 1834524 | C | T | CDS | SMc00268 | Dehydrogenase | Silent mutation (Ala → Ala) |
| | Chr | 1950166 | C | T | CDS | SMc00471 (*cckA*) | Cell cycle histidine kinase | Missense mutation (Arg → His) |
| | pSymA | 81732 | G | - | IGR | SMa0148 - SMa0149 | - | Deletion |
| | pSymB | 279482 | G | T | CDS | SMb20275 | Conserved hypothetical protein | Missense mutation (Ala → Glu) |
| | pSymB | 436989 | G | C | CDS | SMb20422 | Putative alcohol dehydrogenase | Missense mutation (Ala → Gly) |
| | pSymB | 842124 | C | - | CDS | SMb21268 | Putative ABC transporter | Deletion |
| **SmABCΔoriVΔR$_{Inv.}$** | | | | | | | | |
| | Chr | 3506063 | C | T | CDS | SMc03844 | Conserved hypothetical protein | Silent mutation (Val → Val) |
| | pSymA | 81732 | G | - | IGR | SMa0148 - SMa0149 | - | Deletion |
| | pSymB | 279482 | G | T | CDS | SMb20275 | Conserved hypothetical protein | Missense mutation (Ala → Glu) |
| | pSymB | 842124 | C | - | CDS | SMb21268 | Putative ABC transporter | Deletion |
| | pSymB | 850347 | G | T | IGR | SMb21275 - SMb21277 | - | - |
| | pSymB | 852374 | T | A | CDS | SMb21278 (*ade2*) | Probable adenine deaminase | Missense mutation (Asp → Glu) |

**S7 Table (1/3). Bacterial strains used in this study.**

| Strains | Description/ Genotype | References |
|---|---|---|
| ***E.coli*** | | |
| DH5α | Strain for cloning and plasmid extraction, F⁻ *endA1 glnV44 thi-1 recA1 relA1 gyrA96 deoR nupG purB20* φ80d*lac*ZΔM15 Δ(*lacZYA-argF*)U169, hsdR17($r_K^- m_K^+$), λ⁻ | Hanahan, 1983 |
| S17-1 | Donor strain for conjugation with *S. meliloti*, *E. coli* 294 Thi RP4-2-Tc::Mu-Km::Tn7 integrated into the chromosome | Simon et al., 1983 |
| ***S. meliloti*** | | |
| BO33 | Rm1021 carrying excision module for Cre/lox studies | Harrison et al., 2011 |
| SmCreΔhsdR | Rm1021 *cre* expression strain with *hsdR* deletion, *tauX*::*cre-tetRA* ($Tc^r$) | Döhlemann et al., 2016 |
| JDSm97 | SmCreΔhsdR pre-fusion derivative with $P_{aacC1}$ and *loxL* on pSmyA and promoterless *aadA1* and *loxR* on pSymB | This study |
| JDSm98 | JDSm97 after Cre mediated fusion of secondary replicons pSymA and pSymB | This study |
| SmAB | JDSm 98 after sucrose selection mediated and removal of the pJD98 plasmid backbone and active wild type *loxP* site ($Tc^r$, $Spec^r$) | This study |
| JDSm111 | SmAB strain with *loxL* and constitutive promoter P*min2* on pSymAB | This study |
| JDSm118 | JDSm111 with promoterless gentamicin resistance gene *aacC1* and *loxR* site on the chromosome | This study |
| JDSm121 | JDSm118 after Cre mediated fusion of the chromosome and the secondary hybrid replicon pSymAB | This study |
| SmABC | SmAB derivative with an entirely fused genome ($Tc^r$, $Gm^r$, $Spec^r$) | This study |
| SmABΔrepC1 | SmAB derivative lacking *repC1* and the *repC1B1* intergenic region ($Tc^r$, $Spec^r$) | This study |
| SmABΔrepC2 | SmAB derivative lacking *repC2* and the *repC2B2* intergenic region ($Tc^r$, $Spec^r$) | This study |
| SmABCΔrepC1 | SmABC lacking the coding region of *repC1* and corresponding *repBC* integrneic region ($Tc^r$, $Gm^r$, $Spec^r$) | This study |
| SmABCΔrepC2 | SmABC lacking the coding region of *repC2* and corresponding *repBC* integrneic region ($Tc^r$, $Gm^r$, $Spec^r$) | This study |
| SmABCΔoriV | SmABC cured from megaplasmid derived *oriV*s. ($Tc^r$, $Spec^r$, $Gm^r$) | This study |
| SmCreΔhsdR-repA3B3KO | SmCreΔhsdR derivative with *repA3B3* inactivation, *repA3*::pK18mob2 ($Tc^r$, $Km^r$) | This study |
| SmCreΔhsdRΔrepA3B3 | SmCreΔhsdR derivative *repA3B3* deletion, Δsmb1614564 – smb1616866 ($Tc^r$) | This study |
| SmAB-repA3B3KO | SmAB derivative with *repA3B3* inactivation, *repA3*::pK18mob2 ($Tc^r$, $Spec^r$, $Km^r$) | This study |
| SmABΔrepA3B3 | SmAB derivative with *repA3B3* deletion, Δsmb1614564 – smb1616866 ($Tc^r$, $Spec^r$) | This study |
| SmAB-repA1B1C1KO | SmAB derivative with *repA1B1C1* inactivation, *repA1*::pK18mob2 ($Tc^r$, $Spec^r$, $Km^r$) | This study |
| SmABΔrepA1B1C1 | SmAB derivative with *repA1B1C1* deletion, Δsmb53521 – smb59006 ($Tc^r$, $Spec^r$) | This study |
| SmABΔrepA1B1C1-repA3B3KO | SmAB with *repA1B1C1* deletion and *repA3B3* inactivation, Δsmb53521 – smb59006, *repA3*::pK18mob2 ($Tc^r$, $Spec^r$, $Km^r$) | This study |
| SmABC-repA3B3KO | SmABC derivative with *repA3B3* inactivation, *repA3*::pK18mob2 ($Tc^r$, $Gm^r$, $Spec^r$, $Km^r$) | This study |
| SmABCΔrepA3B3 | SmABC derivative with *repA3B3* deletion, Δsmb1614564 – smb1616866 ($Tc^r$, $Gm^r$, $Spec^r$) | This study |
| SmABC-repA1B1C1KO | SmABC derivative with *repA1B1C1* inactivation, *repA1*::pK18mob2 ($Tc^r$, $Gm^r$, $Spec^r$, $Km^r$) | This study |
| SmABCΔrepA1B1C1 | SmAB derivative with *repA1B1C1* deletion, Δsmb53521 – smb59006 ($Tc^r$, $Gm^r$, $Spec^r$) | This study |
| SmABCΔrepA1B1C1-repA3B3KO | SmABC with *repA1B1C1* deletion and *repA3B3* inactivation, Δsmb53521 – smb59006, *repA3*::pK18mob2 ($Tc^r$, $Gm^r$, $Spec^r$, $Km^r$) | This study |
| SmABΔR | SmAB lacking spectinomycin resistance cassette. ($Tc^r$) | This study |
| SmABCΔR | SmABC derivative cured from Gm and Spec antibiotic resistance cassettes ($Tc^r$) | This study |
| SmABCΔoriVΔR | SmABCΔoriV derivative cured from Gm and Spec antibiotic resistance cassettes; C→T transition in the *cckA* sequence ($Tc^r$) | This study |
| SmABCΔoriVΔ$R_{Inv.}$ | SmABCΔoriV derivative cured from Gm and Spec antibiotic resistance cassettes; 3.2 Mbp inversion between *groEL*/*S1* and *groEL*/*S2* | This study |
| MWSm225 | SmCreΔhsdR derivative with *oriC* labeling, *parB::mcerulean* ($Tc^r$) | This study |
| MWSm226 | SmABΔR derivative with *oriC* labeling, *parB::mcerulean* ($Tc^r$) | This study |
| MWSm230 | SmABCΔR derivative with *oriC* labeling, *parB::mcerulean* ($Tc^r$) | This study |
| MWSm285 | SmABCΔoriVΔR derivative with *oriC* labeling, *parB::mcerulean* ($Tc^r$) | This study |

$Km^r$/$Spec^r$/$Tc^r$/$Hyg^r$/$Gm^r$: kanamycin/spectinomycin/tetracycline/hygromycin/gentamicin resistance. smc/a/b: sequence position on the chromosome, pSymA or pSymB, SMc/a/b: CDS annotation.

**S7 Table (2/3). Bacterial strains used in this study.**

| Strains | Description/ Genotype | References |
|---|---|---|
| ***S. meliloti*** | | |
| SmABCΔR-DnaN-mCh | MWSm230 with DnaN fluorophore fusion, *dnaN*::*mcherry* ($Tc^r$, $Gm^r$) | This study |
| SmABCΔoriVΔR-DnaN-mCh | MWSm285 with DnaN fluorophore fusion, *dnaN*::*mcherry* ($Tc^r$, $Gm^r$) | This study |
| SmCreΔhsdR-lacO-2 | MWSm225 with *lacO* array at 0.61 Mbp (60.0°, pAM7 integrated at SMc02244/SMc02245) on the chromosome and pFROS ($Tc^r$, $Gm^r$, $Sp^r$) | This study |
| SmCreΔhsdR-lacO-3 | MWSm225 with *lacO* array at 1.15 Mbp (113.8°, pAM13 integrated at SMc02404/SMc02405) on the chromosome and pFROS ($Tc^r$, $Gm^r$, $Sp^r$) | This study |
| SmCreΔhsdR-tetO-5 | MWSm225 with *tetO* array at 2.44 Mbp (240.2°, pAM32 integrated at SMc01644/SMc01645) on the chromosome and pFROS ($Tc^r$, $Km^r$, $Sp^r$) | This study |
| SmCreΔhsdR-lacO-6 | MWSm225 with *lacO* array at 0.61 Mbp (130.0°, pAM68 integrated at SMb20819/SMb20820) on pSymB and pFROS ($Tc^r$, $Gm^r$, $Sp^r$) | This study |
| SmCreΔhsdR-lacO-9 | MWSm225 with *lacO* array at 0.85 Mbp (182.1°, pAM72 integrated at SMb21278/SMb21279) on pSymB and pFROS ($Tc^r$, $Gm^r$, $Sp^r$) | This study |
| SmCreΔhsdR-lacO-10 | MWSm225 with *lacO* array at 0.43 Mbp (114.2°, pAM45 integrated at SMa0789) on pSymA and pFROS ($Tc^r$, $Gm^r$, $Sp^r$) | This study |
| SmCreΔhsdR-lacO-13 | MWSm225 with *lacO* array at 0.55 Mbp (145.7°, pAM47 integrated at SMa0985/SMa0988) on pSymA and pFROS ($Tc^r$, $Gm^r$, $Sp^r$) | This study |
| SmCreΔhsdR-lacO-14 | MWSm225 with *lacO* array at 0.73 Mbp (156.3°, pAM70 integrated at SMb21089/SMb21090) on pSymB and pFROS ($Tc^r$, $Gm^r$, $Sp^r$) | This study |
| SmCreΔhsdR-lacO-15 | MWSm225 with *lacO* array at 0.67 Mbp (143.2°, pAM69 integrated between SMb21662 and SMb21696) on pSymB and pFROS ($Tc^r$, $Gm^r$, $Sp^r$) | This study |
| SmCreΔhsdR-tetO16 | MWSm225 with *tetO* array at 2.49 Mbp (246.0°, pAM31 integrated at SMc01556/SMc01557) on the chromosome and pFROS ($Tc^r$, $Km^r$, $Sp^r$) | This study |
| SmCreΔhsdR-tetO17 | MWSm225 with *tetO* array at 2.62 Mbp (258.3°, pAM30 integrated at SMc01507) on the chromosome and pFROS ($Tc^r$, $Km^r$, $Sp^r$) | This study |
| SmCreΔhsdR-tetO18 | MWSm225 with *tetO* array at 3.05 Mbp (300.2°, pAM25 integrated at SMc04043/SMc04044) on the chromosome and pFROS ($Tc^r$, $Km^r$, $Sp^r$) | This study |
| SmABCΔR-lacO-2 | MWSm230 with *lacO* array at 0.61 Mbp (32.8°, pAM7 integrated at SMc02244/SMc02245) and pFROS ($Tc^r$, $Gm^r$, $Sp^r$) | This study |
| SmABCΔR-lacO-3 | MWSm230 with *lacO* array at 1.15 Mbp (62.2°, pAM13 integrated at SMc02404/SMc02405) and pFROS ($Tc^r$, $Gm^r$, $Sp^r$) | This study |
| SmABCΔR-tetO-5 | MWSm230 with *tetO* array at 2.44 Mbp (131.3°, pAM32 integrated at SMc01644/SMc01645) and pFROS ($Tc^r$, $Km^r$, $Sp^r$) | This study |
| SmABCΔR-lacO-6 | MWSm230 with *lacO* array at 2.52 Mbp (135.5°, pAM68 integrated at SMb20819/SMb20820) and pFROS ($Tc^r$, $Gm^r$, $Sp^r$) | This study |
| SmABCΔR-lacO-9 | MWSm230 with *lacO* array at 3.96 Mbp (213.2°, pAM72 integrated at SMb21278/SMb21279) and pFROS ($Tc^r$, $Gm^r$, $Sp^r$) | This study |
| SmABCΔR-lacO-10 | MWSm230 with *lacO* array at 4.06 Mbp (219.0°, pAM45 integrated at SMa0789) and pFROS ($Tc^r$, $Gm^r$, $Sp^r$) | This study |
| SmABCΔR-lacO-13 | MWSm230 with *lacO* array at 5.30 Mbp (285.5°, pAM47 integrated at SMa0985/SMa0988) and pFROS ($Tc^r$, $Gm^r$, $Sp^r$) | This study |
| SmABCΔR-lacO-14 | MWSm230 with *lacO* array at 5.43 Mbp (292.5°, pAM70 integrated at SMb21089/SMb21090) and pFROS ($Tc^r$, $Gm^r$, $Sp^r$) | This study |
| SmABCΔR-lacO-15 | MWSm230 with *lacO* array at 5.49 Mbp (295.8°, pAM69 integrated between SMb21662 and SMb21696) and pFROS ($Tc^r$, $Gm^r$, $Sp^r$) | This study |
| SmABCΔR-tetO-16 | MWSm230 with *tetO* array at 5.53 Mbp (297.7°, pAM31 integrated at SMc01556/SMc01557) and pFROS ($Tc^r$, $Km^r$, $Sp^r$) | This study |
| SmABCΔR-tetO-17 | MWSm230 with *tetO* array at 5.65 Mbp (304.4°, pAM30 integrated at SMc01507) and pFROS ($Tc^r$, $Km^r$, $Sp^r$) | This study |
| SmABCΔR-tetO-18 | MWSm230 with *tetO* array at 6.08 Mbp (327.3°, pAM25 integrated at SMc04043/SMc04044) and pFROS ($Tc^r$, $Km^r$, $Sp^r$) | This study |
| SmCreΔhsdR *oriC-terC* | MWSm225 with *tetO* array at 1.72 Mbp (169.7°, pJD169 at SMc01205) for *terC* labeling on the chromosome and pFROS ($Tc^r$, $Km^r$, $Sp^r$) | This study |
| SmCreΔhsdR *oriC/A-terA* | MWSm225 with *tetO* array at 1.35 Mbp (358.0°, sma1346524::$tetO_{120}$) for *oriA* labeling, *lacO* array at 0.65 Mbp (174.0°, pMW193 integrated at SMa1188) for *terA* labeling on pSymA and pFROS ($Tc^r$, $Km^r$, $Sp^r$) | This study |
| SmCreΔhsdR *oriC/B-terB* | MWSm225 with *lacO* array at 0.05 Mbp (11.2°, smb52247::$lacO_{120}$) for *oriB* labeling, *tetO* array at 1.10 Mbp (234.4°, pJD170 integrating at SMb21555) for *terB* labeling on pSymB and pFROS ($Tc^r$, $Km^r$, $Sp^r$) | This study |
| SmCreΔhsdR *oriC/A/B* | MWSm225 with *tetO* array at 1.35 Mbp (358.0°, sma1346524::$tetO_{120}$) for *oriA* labeling on pSymA, *lacO* array at 0.05 Mbp (11.2°, smb52247::$lacO_{120}$) for *oriB* labeling on pSymB and pFROS ($Tc^r$, $Sp^r$) | This study |
| SmABΔR *oriC-terC* | MWSm226 with *tetO* array at 1.72 Mbp (169.7°, pJD169 at SMc01205) for *terC* labeling on the chromosome and pFROS ($Tc^r$, $Km^r$, $Sp^r$) | This study |
| SmABΔR *oriC/A-terA* | MWSm226 with *tetO* array at 3.03 Mbp (359.6°, sma3033862::$tetO_{120}$) for *oriA* labeling, *lacO* array at 2.34 Mbp (277.6°, pMW193 integrated at SMa1188) for *terA* labeling on pSymAB and pFROS ($Tc^r$, $Km^r$, $Sp^r$) | This study |
| SmABΔR *oriC/B-terB* | MWSm226 with *lacO* array at 1.43 Mbp (170.0°, smb1425712::$lacO_{120}$) for *oriB* labeling, *tetO* array at 0.79 Mbp (93.0°, pJD170 integrating at SMb21555) for *terB* labeling on pSymAB and pFROS ($Tc^r$, $Km^r$, $Sp^r$) | This study |
| SmABΔR *oriC/A/B* | MWSm226 with *tetO* array 3.03 Mbp (359.6°, sma3033862::$tetO_{120}$) for *oriA* labeling, *lacO* array 1.43 Mbp (170.0°, smb1425712::$lacO_{120}$) for *oriB* labeling on pSymAB and pFROS ($Tc^r$, $Sp^r$) | This study |

$Km^r$/$Spec^r$/$Tc^r$/$Hyg^r$/$Gm^r$: kanamycin/spectinomycin/tetracycline/hygromycin/gentamicin resistance. smc/a/b: sequence position on the chromosome, pSymA or pSymB, SMc/a/b: CDS annotation.

**S7 Table (3/3). Bacterial strains used in this study.**

| Strains | Description/ Genotype | References |
|---|---|---|
| ***S. meliloti*** | | |
| SmABCΔR *oriC-terC* | MWSm230 with *tetO* array at 1.72 Mbp (92.7°, pJD169 at SMc01205) for *terC* labeling on ABC and pFROS ($Tc^r$, $Km^r$, $Sp^r$) | This study |
| SmABCΔR *oriC/A-terA* | MWSm230 with *tetO* array at 4.50 Mbp (242.5°, sma4501016::$tetO_{120}$) for *oriA* labeling, *lacO* array at 5.19 Mbp (279.8°, pMW193 integrated at SMa1188) for *terA* labeling on ABC and pFROS ($Tc^r$, $Km^r$, $Sp^r$) | This study |
| SmABCΔR *oriC/B-terB* | MWSm230 with *lacO* array at 3.07 Mbp (165.5°, smb3071828::$lacO_{120}$) for *oriB* labeling, *tetO* array at 3.71 Mbp (200.0°, pJD170 integrating at SMb21555) for *terB* labeling on ABC and pFROS ($Tc^r$, $Km^r$, $Sp^r$) | This study |
| SmABCΔR *oriC/A/B* | MWSm230 with *tetO* array 4.50 Mbp (242.5°, sma4501016::$tetO_{120}$) for *oriA* labeling, *lacO* array at 3.07 Mbp (165.5°, smb3071828::$lacO_{120}$) for *oriB* labeling on ABC and pFROS ($Tc^r$, $Sp^r$) | This study |
| SmABCΔoriVΔR *oriC-terC* | MWSm285 with *tetO* array at 1.72 Mbp (92.8°, pJD169 at SMc01205) for *terC* labeling on ABCΔoriV and pFROS ($Tc^r$, $Km^r$, $Sp^r$) | This study |
| SmABCΔoriVΔR *oriC/A-terA* | MWSm285 with *tetO* array at 4.50 Mbp (242.5°, sma4498131::$tetO_{120}$) for *oriA* labeling, *lacO* array at 5.19 Mbp (279.8°, pMW193 integrated at SMa1188) for *terA* labeling on ABCΔoriV and pFROS ($Tc^r$, $Km^r$, $Sp^r$) | This study |
| SmABCΔoriVΔR *oriC/B-terB* | MWSm285 with *lacO* array at 3.07 Mbp (165.5°, smb3070368::$lacO_{120}$) for *oriB* labeling, *tetO* array at 3.71 Mbp (200.1°, pJD170 integrating at SMb21555) for *terB* labeling on ABCΔoriV and pFROS ($Tc^r$, $Km^r$, $Sp^r$) | This study |
| SmABCΔoriVΔR *oriC/A/B* | MWSm285 with *tetO* array 4.50 Mbp (242.5°, sma4498131::$tetO_{120}$) for *oriA* labeling, *lacO* array at 3.07 Mbp (165.5°, smb3070368::$lacO_{120}$) for *oriB* labeling on ABCΔoriV and pFROS ($Tc^r$, $Sp^r$) | This study |
| SmCreΔhsdR $CckA^{R436H}$ | MWSm225 with mutagenized *cckA* sequence (C→T transition) ($Tc^r$) | This study |
| SmABCΔR $CckA^{R436H}$ | MWSm230 with mutagenized *cckA* sequence (C→T transition) ($Tc^r$) | This study |
| SmABC *oriC* | SmABC derivative with *oriC* labeling, *parB*::pMW261 ($Tc^r$, $Km^r$) | This study |
| SmABCΔoriV *oriC* | SmABCΔoriV derivative with *oriC* labeling, *parB*::pMW261 ($Tc^r$, $Km^r$) | This study |

$Km^r$/$Spec^r$/$Tc^r$/$Hyg^r$/$Gm^r$: kanamycin/spectinomycin/tetracycline/hygromycin/gentamicin resistance. smc/a/b: sequence position on the chromosome, pSymA or pSymB, SMc/a/b: CDS annotation.

**S8 Table (1/2). Plasmids used in this study.**

| Plasmids | Description | References |
|---|---|---|
| **pK18mob2** | Suicide vector carrying replication origin pMB1 | Tauch et al., 1998 |
| **pK18mobsacB** | Suicide vector carrying *sacB* for sucrose selection | Schäfer et al., 1994 |
| **pSRK-Gm/Km** | Broad-host-range expression vector with pBBR1 based *oriV* | Khan et al., 2008 |
| **pAGM8031** | broad-host-range expression vector with RK2 based *oriV* | Weber et al., 2011 |
| **pMS252** | pUC derivative carrying a gentamicin resistance cassette derived from RlO33 [Tn1696] | Becker et al., 1995 |
| **pLAU53** | FROS plasmid carrying *tetR* and *lacI* | Lau et al., 2003 |
| **pLAU43** | FROS plasmid carrying 2 arrays of 120 *lac* operators | Lau et al., 2003 |
| **pLAU44** | FROS plasmid carrying 2 arrays of 120 *tet* operators | Lau et al., 2003 |
| **pWBT5$^{cerulean}$** | Used for PCR amplification of *cerulean* | Schlüter et al., 2015 |
| **pLoriT** | pK18mob2 derivative used for cloning of FROS labelling constructs | Döhlemann et al., 2017 |
| **pK19ms DnaN-mCherry** | pK19mobsacB derivative with *dnaN-mCherry* translational fusion | Frage et al., 2016 |
| **pG18mob-mVenus** | Used for PCR amplification of *mVENUS* | Elizaveta Krol |
| **pSRKGm-CmChr** | Used for PCR amplification of *mCherry* | Elizaveta Krol |
| **pYEM1** | pAGM8031 derivative eqipped with a multiple cloning site | Yannick End |
| **pAM7** | Mobilizable pACYC177 derivative with Gm$^r$ cassette carrying a *lacO* array (120 copies) and *S. meliloti* genome sequence smc608072 – smc609077 | Anna Motnenko |
| **pAM13** | Mobilizable pACYC177 derivative with Gm$^r$ cassette carrying a *lacO* array (120 copies) and *S. meliloti* genome sequence smc1156615 – smc1157633 | Anna Motnenko |
| **pAM25** | pK18mob2 derivative carrying a *tetO* array (120 copies) and *S. meliloti* genome sequence smc3046746 – smc3047760 | Anna Motnenko |
| **pAM30** | pK18mob2 derivative carrying a *tetO* array (120 copies) and *S. meliloti* genome sequence smc2621880 – smc2622890 | Anna Motnenko |
| **pAM31** | pK18mob2 derivative carrying a *tetO* array (120 copies) and *S. meliloti* genome sequence smc2497039 – smc2498058 | Anna Motnenko |
| **pAM32** | pK18mob2 derivative carrying a *tetO* array (120 copies) and *S. meliloti* genome sequence smc2438176 – smc2439176 | Anna Motnenko |
| **pAM45** | Mobilizable pACYC177 derivative with Gm$^r$ cassette carrying a *lacO* array (120 copies) and *S. meliloti* genome sequence sma428900 – sma429665 | Anna Motnenko |
| **pAM47** | Mobilizable pACYC177 derivative with Gm$^r$ cassette carrying a *lacO* array (120 copies) and *S. meliloti* genome sequence sma547269 – sma548268 | Anna Motnenko |
| **pAM68** | Mobilizable pACYC177 derivative with Gm$^r$ cassette carrying a *lacO* array (120 copies) and *S. meliloti* genome sequence smb608355 – smb609352 | Anna Motnenko |
| **pAM69** | Mobilizable pACYC177 derivative with Gm$^r$ cassette carrying a *lacO* array (120 copies) and *S. meliloti* genome sequence smb670529 – smb671370 | Anna Motnenko |
| **pAM70** | Mobilizable pACYC177 derivative with Gm$^r$ cassette carrying a *lacO* array (120 copies) and *S. meliloti* genome sequence smb731852 – smb732852 | Anna Motnenko |
| **pAM72** | Mobilizable pACYC177 derivative with Gm$^r$ cassette carrying a *lacO* array (120 copies) and *S. meliloti* genome sequence smb852585 – smb853589 | Anna Motnenko |
| **pJD98** | pK18mobsacB carrying the *thrA-rpoC* terminator sequence, *loxR*, *aadA1* and the *S. meliloti* genome sequence smb789438 – smb789925. Used for construction of *S. meliloti* SmAB | This study |
| **pJD99** | pK18mobsacB carrying *PaacC1*, *loxL* and the *S. meliloti* genome sequence sma471764 – sma472263 and sma472264 – sma472736. Used for construction of *S. meliloti* SmAB | This study |
| **pJD126** | pK18mobsacB carrying the *thrA-rpoC* terminator sequence, *loxR*, *aacC1* and the *S. meliloti* genome sequence smc2493479 – smc2493978. Used for construction of *S. meliloti* SmABC | This study |
| **pJD130** | pK18mobsacB carrying *Pmin2*, *loxL* and the *S. meliloti* genome sequence smb637066 – smb637566 and smb637565 – smb638065. Used for construction of *S. meliloti* SmABC | This study |
| **pJD169** | pK18mob2 derivative with *tetO* array and *S. meliloti* genome sequence smc1721001-smc1721494 for integration at *terC* | This study |
| **pJD170** | pK18mob2 derivative with *tetO* array and *S. meliloti* genome sequence smb1097631-smb1098128 for integration at *terB* | This study |
| **pJD201** | pK18mobsacB derivative for deletion of *repC2* | This study |
| **pJD202** | pK18mobsacB derivative for deletion of *repC1* | This study |
| **pJD206** | pK18mob2 derivative for *repA2B2C2* kockout | This study |
| **pJD207** | pK18mob2 derivative for *repA1B1C1* kockout | This study |
| **pJD222** | pK18mobsacB derivative for deletion of a spectinomycin resistance cassette in *S. meliloti* replicon fusion strains | This study |
| **pJD225** | pK18mobsacB derivative for deletion of *repA3B3* on pSymB | This study |

smc/a/b: sequence position on the chromosome, pSymA or pSymB

**S8 Table (2/2). Plasmids used in this study.**

| Plasmids | Description | References |
|---|---|---|
| **pJD226** | pK18mobsacB derivative for deletion of the *hemE* promoter region. | This study |
| **pJD227** | pK18mobsacB derivative for deletion of the *hemE* promoter region and DnaA boxes 2-4. | This study |
| **pJD228** | pK18mobsacB derivative for deletion of DnaA boxes 2-4 in *oriC* | This study |
| **pJD229** | pK18mobsacB derivative for deletion of a gentamicin resistance cassette in *S. meliloti* replicon fusion strains | This study |
| **pJD266** | pK18mob2 derivative for *repA3B3* kockout | This study |
| **pMW186** | pK18mobsacB derivative with *tetO* array and *S. meliloti* genome sequence sma1345524-sma1346523 and sma1346524-sma1347522 for insertion at *oriA* | This study |
| **pMW188** | pK18mobsacB derivative with *lacO* array and *S. meliloti* genome sequence smb51601-smb52246 and smb52247-smb52897 for insertion at *oriB* | This study |
| **pMW189** | pK18mobsacB derivative for genomic integration of *tetR-mVenus* and *lacI-mChr* under control of P*tau* | This study |
| **pMW193** | pK18mob2 derivative with *lacO* array and *S. meliloti* genome sequence sma654101-sma654590 for integration at *terA* | This study |
| **pMW198** | pK18mobsacB derivative for markerless in-frame fusion of *parB* with *cerulean* | This study |
| **pMW210** | pK18mobsacB derivative for deletion of *repA1B1C1* including three related *parS* like sequences on pSymB | This study |
| **pMW211** | pK18mobsacB derivative for deletion of *repA2B2C2* including six related *parS* like sequences on pSymA | This study |
| **pMW212 (pK18mob3)** | pK18mob2 integration vector derivative without $P_{lac}$ promoter activity (cured from $P_{lac}$ and parts of the *lacZa* fragment) | This study |
| **pMW230** | pK18mobsacB derivative for deletion of six *parS* like sequences upstream of *repA2B2C2* on pSymA | |
| **pMW256** | pK18mobsacB derivative for replacement of native *cckA* sequence with the *cckA* C→T ( bp 1307) SNV version | This study |
| **pMW257** | pK18mobsacB derivative for repair of the *cckA* C→T (bp 1307) transition in SmABCΔoriVΔR | This study |
| **pMW261** | pK18mob3 derivative for in frame fusion of *parB* with *cerulean* (with backbone integration) | This study |
| **pFROS** | Mobilizable pAGM8031 derivative carrying *tetR-mVenus* and *lacI-mChr* under control of P*tau* | This study |

smc/a/b: sequence position on the chromosome, pSymA or pSymB

**S9 Table (1/3). Construction of *S. meliloti* replicon fusion strains and derivatives.**

| Strains | Construction description |
|---|---|
| **SmAB** | *S. melilo*ti **SmCreΔhsdR** (Döhlemann et al., 2016) was transformed with pJD99 thereby equipping pSymA with pMS252 derived gentamicin resistance cassette promoter $P_{aacC1}$ and *loxL*. Deletion of the plasmid backbone via sucrose selection allowed subsequent insertion of pJD98 thereby providing promoterless *aadA1* and *loxR* on pSymB in resulting strain **JDSm97**. Cre/*lox* recombination resulted in co-integration of pSymA and pSymB thereby translocate $P_{aacC1}$ in front of *aadA1.*This turned the newly strain **JDSm98** resistant to spectinomycin ($Spec^R$) which allowed for positive selection on medium supplemented with the same. Megaplasmid fusion brought the homologous *nodPQ* clusters into close proximity, allowing for subsequent deletion of the pJD98 derived plasmid backbone and active wild type *loxP* site (resulting from *loxL*/*R* mediated recombination) via sucrose selection and creating final strain **SmAB**. |
| **SmABC** | *S. meliloti* strain SmAB was transformed with pJD130 thereby equipping pSymAB with constitutive promoter $P_{min2}$ (Döhlemann et al., 2017) and *loxL*. The plasmid backbone was deleted via sucrose selection resulting in strain **JDSm111**. This strain was then transformed with pJD126 thereby equipping the chromosome with the promoterless gentamicin resistance gene *aacC1* and *loxR* site. Cre/*lox* recombination resulted in fusion of pSymAB and the chromosome in strain **JDSm118**. Since both *lox* sites were integrated next to the *algI* homologs of pSymB and the chromosome the site specific recombination enabled subsequent sucrose selection mediated deletion of the remaining pJD126 backbone including the *loxP site* via those now closely located homologous regions generating **SmABC**. |
| **SmABΔrepC1** | **SmAB** was transformed with pJD202, allowing for sucrose selection mediated deletion of *repC1* and the corresponding *repBC* intergenic region. |
| **SmABΔrepC2** | **SmAB** was transformed with pJD201, allowing for sucrose selection mediated deletion of *repC2* and the corresponding *repBC* intergenic region. |
| **SmABCΔrepC1** | **SmABC** was transformed with pJD202, allowing for sucrose selection mediated deletion of *repC1* and the corresponding *repBC* intergenic region. |
| **SmABCΔrepC2** | **SmABC** was transformed with pJD201, allowing for sucrose selection mediated deletion of *repC2* and the corresponding *repBC* intergenic region. |
| **SmABCΔoriV** | **SmABCΔrepC2** was transformed with pJD202, allowing for sucrose selection mediated deletion of *repC1* and the corresponding *repBC* intergenic region. |
| **SmCreΔhsdR-repA3B3KO** | **SmCreΔhsdR** was transformed with pJD266 integrating and interrupting the coding sequence of *repA3* on pSymB. |
| **SmCreΔhsdRΔrepA3B3** | **SmCreΔhsdR** was transformed with pJD225, allowing for sucrose selection mediated deletion of *repA3B3* on pSymB. |
| **SmAB-repA3B3KO** | **SmAB** was transformed with pJD266 integrating and interrupting the coding sequence of *repA3* on pSymAB. |
| **SmABΔrepA3B3** | **SmAB** was transformed with pJD225, allowing for sucrose selection mediated deletion of *repA3B3* on pSymAB. |
| **SmAB-repA1B1C1KO** | **SmAB** was transformed with pJD207 integrating and interrupting the coding sequence of *repA1* on pSymAB. |
| **SmABΔrepA1B1C1** | **SmAB** was transformed with pMW210, allowing for sucrose selection mediated deletion of *repA1B1C1* on pSymAB. |
| **SmABΔrepA1B1C1-repA3B3KO** | **SmABΔrepA1B1C1** was transformed with pJD266 integrating and interrupting the coding sequence of *repA3* on pSymAB. |
| **SmABC-repA3B3KO** | **SmABC** was transformed with pJD266 integrating and interrupting the coding sequence of *repA3*. |
| **SmABCΔrepA3B3** | **SmABC** was transformed with pJD225, allowing for sucrose selection mediated deletion of *repA3B3*. |
| **SmABC-repA1B1C1KO** | **SmABC** was transformed with pJD207 integrating and interrupting the coding sequence of *repA1*. |
| **SmABCΔrepA1B1C1** | **SmABC** was transformed with pMW210, allowing for sucrose selection mediated deletion of *repA1B1C1*. |
| **SmABCΔrepA1B1C1-repA3B3KO** | **SmABCΔrepA1B1C1** was transformed with pJD266 integrating and interrupting the coding sequence of *repA3*. |
| **SmABΔR** | **SmAB** was transformed with pJD222, allowing for sucrose selection mediated deletion of *aadA1*. |
| **SmABCΔR** | **SmABC** was transformed with pJD222, allowing for sucrose selection mediated deletion of $P_{aacC1}$, *loxLE* and *aadA1*. Subsequent, resulting strain was then transformed with pJD229 for deletion of $P_{min2}$, *loxLR* and *aacC1*. |
| **SmABCΔoriVΔR** | **SmABCΔoriV** was transformed with pJD229, allowing for sucrose selection mediated deletion of $P_{min2}$, *loxLR* and *aacC1*. Subsequent, resulting strain was then transformed with pJD222 for deletion of $P_{aacC1}$, *loxLE* and *aadA1*. |
| **SmABCΔoriVΔR$_{Inv.}$** | **SmABCΔoriV**was transformed with pJD222, allowing for sucrose selection mediated deletion of $P_{aacC1}$, *loxLE* and *aadA1*. Subsequent, resulting strain was then transformed with pJD229 for deletion of $P_{min2}$, *loxLR* and *aacC1*. |
| **MWSm225** | **SmCreΔhsdR** was transformed with pMW198, allowing sucrose selection mediated markerless translational fusion of native *parB* with coding sequence of Cerlulean. |
| **MWSm226** | **SmABΔR** was transformed with pMW198, allowing sucrose selection mediated markerless translational fusion of native *parB* with coding sequence of Cerlulean. |
| **MWSm230** | **SmABCΔR** was transformed with pMW198, allowing sucrose selection mediated markerless translational fusion of native *parB* with coding sequence of Cerlulean. |
| **MWSm285** | **SmABCΔoriVΔR** was transformed with pMW198, allowing sucrose selection mediated markerless translational fusion of native *parB* with coding sequence of Cerlulean. |
| **SmABCΔR-DnaN-mCh** | **MWSm230** was transformed with pK19ms DnaN-mCherry, allowing sucrose selection mediated translational fusion of native *dnaN* with coding sequence of mCherry. |
| **SmABCΔoriVΔR-DnaN-mCh** | **MWSm285** was transformed with pK19ms DnaN-mCherry, allowing sucrose selection mediated translational fusion of native *dnaN* with coding sequence of mCherry. |

$Km^r/Spec^r/Tc^r/Hyg^r/Gm^r$: kanamycin/spectinomycin/tetracycline/hygromycin/gentamicin resistance. smc/a/b: sequence position on the chromosome, pSymA or pSymB, SMc/a/b: CDS annotation.

**S9 Table (2/3). Construction of *S. meliloti* replicon fusion strains and derivatives.**

| Strains | Construction description |
|---|---|
| **SmCreΔhsdR-lacO-2** | **MWSm225** was transformed with pAM7 integrating at SMc02244/SMc02245 on the chromosome. Subsequent resulting strain was transformed with pFROS. |
| **SmCreΔhsdR-lacO-3** | **MWSm225** was transformed with pAM13 integrating at SMc02404/SMc02405 on the chromosome. Subsequent resulting strain was transformed with pFROS. |
| **SmCreΔhsdR-tetO-5** | **MWSm225** was transformed with pAM32 integrating at SMc01644/SMc01645 on the chromosome. Subsequent resulting strain was transformed with pFROS. |
| **SmCreΔhsdR-lacO-6** | **MWSm225** was transformed with pAM68 integrating at SMb20819/SMb20820 on pSymB. Subsequent resulting strain was transformed with pFROS. |
| **SmCreΔhsdR-lacO-9** | **MWSm225** was transformed with pAM72 integrating at SMb21278/SMb21279 on pSymB. Subsequent resulting strain was transformed with pFROS. |
| **SmCreΔhsdR-lacO-10** | **MWSm225** was transformed with pAM45 integrating at SMa0789 on pSymA. Subsequent resulting strain was transformed with pFROS. |
| **SmCreΔhsdR-lacO-13** | **MWSm225** was transformed with pAM47 integrating at SMa0985/SMa0988 on pSymA. Subsequent resulting strain was transformed with pFROS. |
| **SmCreΔhsdR-lacO-14** | **MWSm225** was transformed with pAM70 integrating at SMb21089/SMb21090 on pSymB. Subsequent resulting strain was transformed with pFROS. |
| **SmCreΔhsdR-lacO-15** | **MWSm225** was transformed with pAM69 integrating between SMb21662 and SMb21696 on pSymB. Subsequent resulting strain was transformed with pFROS. |
| **SmCreΔhsdR-tetO16** | **MWSm225** was transformed with pAM31 integrating at SMc01556/SMc01557 on the chromosome. Subsequent resulting strain was transformed with pFROS. |
| **SmCreΔhsdR-tetO17** | **MWSm225** was transformed with pAM30 integrating at SMc01507 on the chromosome. Subsequent resulting strain was transformed with pFROS. |
| **SmCreΔhsdR-tetO18** | **MWSm225** was transformed with pAM25 integrating at SMc04043/SMc04044 on the chromosome. Subsequent resulting strain was transformed with pFROS. |
| **SmABCΔR-lacO-2** | **MWSm230** was transformed with pAM7 integrating at SMc02244/SMc02245. Subsequent resulting strain was transformed with pFROS. |
| **SmABCΔR-lacO-3** | **MWSm230** was transformed with pAM13 integrating at SMc02404/SMc02405. Subsequent resulting strain was transformed with pFROS. |
| **SmABCΔR-tetO-5** | **MWSm230** was transformed with pAM32 integrating at SMc01644/SMc01645. Subsequent resulting strain was transformed with pFROS. |
| **SmABCΔR-lacO-6** | **MWSm230** was transformed with pAM68 integrating at SMb20819/SMb20820. Subsequent resulting strain was transformed with pFROS. |
| **SmABCΔR-lacO-9** | **MWSm230** was transformed with pAM72 integrating at SMb21278/SMb21279. Subsequent resulting strain was transformed with pFROS. |
| **SmABCΔR-lacO-10** | **MWSm230** was transformed with pAM45 integrating at SMa0789. Subsequent resulting strain was transformed with pFROS. |
| **SmABCΔR-lacO-13** | **MWSm230** was transformed with pAM47 integrating at SMa0985/SMa0988. Subsequent resulting strain was transformed with pFROS. |
| **SmABCΔR-lacO-14** | **MWSm230** was transformed with pAM70 integrating at SMb21089/SMb21090. Subsequent resulting strain was transformed with pFROS. |
| **SmABCΔR-lacO-15** | **MWSm230** was transformed with pAM69 integrating between SMb21662 and SMb21696. Subsequent resulting strain was transformed with pFROS. |
| **SmABCΔR-tetO-16** | **MWSm230** was transformed with pAM31 integrating at SMc01556/SMc01557. Subsequent resulting strain was transformed with pFROS. |
| **SmABCΔR-tetO-17** | **MWSm230** was transformed with pAM30 integrating at SMc01507. Subsequent resulting strain was transformed with pFROS. |
| **SmABCΔR-tetO-18** | **MWSm230** was transformed with pAM25 integrating at SMc04043/SMc04044. Subsequent resulting strain was transformed with pFROS. |
| **SmCreΔhsdR *oriC-terC*** | **MWSm225** was transformed with pJD169 integrating at SMc01205 on the chromosome. Subsequent resulting strain was transformed with pFROS. |
| **SmCreΔhsdR *oriC/A-terA*** | **MWSm225** was transformed with pMW186, allowing sucrose selection mediated markerless integration of $tetO_{120}$ in the SMa2383 – SMa2385 intergenic region. Resulting strain was transformed with pMW193 for $lacO_{120}$ integration at SMa1188 and subsequent equipped with pFROS. |
| **SmCreΔhsdR *oriC/B-terB*** | **MWSm225** was transformed with pMW188, allowing sucrose selection mediated markerless integration of $lacO_{120}$ in the SMb20041 – SMb20042 intergenic region. Resulting strain was transformed with pJD170 for $tetO_{120}$ integration at SMb21555 and subsequent equipped with pFROS. |
| **SmCreΔhsdR *oriC/A/B*** | **MWSm225** was successive transformed with pMW188 and pMW186, allowing sucrose selection mediated integration of $lacO_{120}$ in the SMb20041 – SMb20042 and $tetO_{120}$ in the SMa2383 – SMa2385 intergenic region. Subsequent, resulting strain was transformed with pFROS. |
| **SmABΔR *oriC-terC*** | **MWSm226** was transformed with pJD169 integrating at SMc01205 on the chromosome. Subsequent resulting strain was transformed with pFROS. |
| **SmABΔR *oriC/A-terA*** | **MWSm226** was transformed with pMW186, allowing sucrose selection mediated markerless integration of $tetO_{120}$ in the SMa2383 – SMa2385 intergenic region. Resulting strain was transformed with pMW193 for $lacO_{120}$ integration at SMa1188 and subsequent equipped with pFROS. |
| **SmABΔR *oriC/B-terB*** | **MWSm226** was transformed with pMW188, allowing sucrose selection mediated markerless integration of $lacO_{120}$ in the SMb20041 – SMb20042 intergenic region. Subsequent, resulting strain was transformed with pJD170 for $tetO_{120}$ integration at SMb21555 and subsequent equipped with pFROS. |
| **SmABΔR *oriC/A/B*** | **MWSm226** was successive transformed with pMW188 and pMW186, allowing sucrose selection mediated integration of $lacO_{120}$ in the SMb20041 – SMb20042 and $tetO_{120}$ in the SMa2383 – SMa2385 intergenic region. Subsequent, resulting strain was transformed with pFROS. |
| **SmABCΔR *oriC-terC*** | **MWSm230** was transformed with pJD169 integrating at SMc01205 on the chromosome. Subsequent resulting strain was transformed with pFROS. |
| **SmABCΔR *oriC/A-terA*** | **MWSm230** was transformed with pMW186, allowing sucrose selection mediated markerless integration of $tetO_{120}$ in the SMa2383 – SMa2385 intergenic region. Resulting strain was transformed with pMW193 for $lacO_{120}$ integration at SMa1188 and subsequent equipped with pFROS. |
| **SmABCΔR *oriC/B-terB*** | **MWSm230** was transformed with pMW188, allowing sucrose selection mediated markerless integration of $lacO_{120}$ in the SMb20041 – SMb20042 intergenic region. Resulting strain was transformed with pJD170 for $tetO_{120}$ integration at SMb21555 and subsequent equipped with pFROS. |
| **SmABCΔR *oriC/A/B*** | **MWSm230** was successive transformed with pMW188 and pMW186, allowing sucrose selection mediated integration of $lacO_{120}$ in the SMb20041 – SMb20042 and $tetO_{120}$ in the SMa2383 – SMa2385 intergenic region. Subsequent, resulting strain was transformed with pFROS. |

$Km^r$/$Spec^r$/$Tc^r$/$Hyg^r$/$Gm^r$: kanamycin/spectinomycin/tetracycline/hygromycin/gentamicin resistance. smc/a/b: sequence position on the chromosome, pSymA or pSymB, SMc/a/b: CDS annotation.

**S9 Table (3/3). Construction of *S. meliloti* replicon fusion strains and derivatives.**

| Strains | Construction description |
|---|---|
| **SmABCΔoriVΔR *oriC-terC*** | **MWSm285** was transformed with pJD169 integrating at SMc01205 on the chromosome. Subsequent resulting strain was transformed with pFROS. |
| **SmABCΔoriVΔR *oriC/A-terA*** | **MWSm285** was transformed with pMW186, allowing sucrose selection mediated markerless integration of $tetO_{120}$ in the SMa2383 – SMa2385 intergenic region. Resulting strain was transformed with pMW193 for $lacO_{120}$ integration at SMa1188 and subsequent equipped with pFROS. |
| **SmABCΔoriVΔR *oriC/B-terB*** | **MWSm285** was transformed with pMW188, allowing sucrose selection mediated markerless integration of $lacO_{120}$ in the SMb20041 – SMb20042 intergenic region. Resulting strain was transformed with pJD170 for $tetO_{120}$ integration at SMb21555 and subsequent equipped with pFROS. |
| **SmABCΔoriVΔR *oriC/A/B*** | **MWSm285** was successive transformed with pMW188 and pMW186, allowing sucrose selection mediated integration of $lacO_{120}$ in the SMb20041 – SMb20042 and $tetO_{120}$ in the SMa2383 – SMa2385 intergenic region. Subsequent, resulting strain was transformed with pFROS. |
| **SmCreΔhsdR CckA$^{R436H}$** | **MWSm225** was transformed with pMW256 allowing sucrose selection mediated C→T transition at position 1307bp in the *cckA* (SMc00471) sequence. |
| **SmABCΔR CckA$^{R436H}$** | **MWSm230** was transformed with pMW256 allowing sucrose selection mediated C→T transition at position 1307bp in the *cckA* (SMc00471) sequence. |
| **SmABC *oriC*** | **SmABC** was transformed with pMW261 allowing translational fusion of native *parB* with coding sequence of Cerlulean. |
| **SmABCΔoriV *oriC*** | **SmABCΔoriV** was transformed with pMW261 allowing translational fusion of native *parB* with coding sequence of Cerlulean. |

Km$^r$/Spec$^r$/Tc$^r$/Hyg$^r$/Gm$^r$: kanamycin/spectinomycin/tetracycline/hygromycin/gentamicin resistance. smc/a/b: sequence position on the chromosome, pSymA or pSymB, SMc/a/b: CDS annotation.

**S10 Table (1/2). Plasmid construction.**

| Construct | Construction |
|---|---|
| pJD98 | The *thrA-rpoC* terminator sequence was PCR amplified with primers 131 and 132 from *S. meliloti* B033 gDNA. A *loxR* site was generated via hybridization of primers 189 and 190. The spectinomycin resistance cassette *aadA1* was PCR amplified with primers 127 and 128 from *S. meliloti* B033 gDNA. A ~500bp DNA fragment for homologous integration downstream of *nodQ2* (HR_nodQ2) was PCR amplified from *S. meliloti* Rm1021 gDNA with primers 240 and 241. Then, the *thrA-rpoC* terminator sequence (digested with BamHI/NdeI), the *loxR* site (NdeI/SalI compatible 5‘ phosphorylated cohesive ends), and *aadA1* (digested with SalI/PstI) were ligated into BamHI/PstI-opened pK18mobsacB. The resulting precursor construct was digested with ScaI and SphI, and finally equipped with SphI-digested HR_nodQ2. |
| pJD99 | Two ~500bp DNA fragments for homologous integration upstream of *nodP1* (HR1_nodP1 and HR2_nodP1) were PCR amplified from *S. meliloti* gDNA with primer pairs 236/237 and 238/239, respectively. Gentamicin resistance cassette promoter P*aacC1* was PCR amplified with primers 6 and 7 from pMS252. A *loxL* site was generated via hybridzation of primers 187 and 188. Then, HR1_nodP1 (digested with EcoRI/BamHI), P*aacC1* (digested with BamHI/NdeI), *loxL* (compatible 5‘ phosphorylated cohesive ends) and HR2_nodP1 (SalI/SphI) were ligated into EcoRI/SphI-opened pK18mobsacB. |
| pJD126 | The *thrA-rpoC* terminator sequence was PCR amplified with primers 131 and 132 from *S. meliloti* B033 gDNA. A *loxR* site was generated via hybridization of primers 189 and 190. The gentamicin resistance cassette *aacC1* was PCR amplified with primers 125 and 126 from pMS252. A ~500bp DNA fragment for homologous integration downstream of the chromosomal *algI* homolog (HR_algI_Chr) was PCR amplified with primers 364 and 365 from *S. meliloti* Rm1021 gDNA. Then, the *thrA-rpoC* terminator sequence (digested with BamHI/NdeI), the *loxR* site (compatible 5‘ phosphorylated cohesive ends), and *aacC1* (digested with SalI/PstI) were ligated into BamHI/PstI-opened pK18mobsacB. The resulting precursor construct was digested with ScaI and finally equipped with HR_algI (5‘ phosphorylated blunt ends). |
| pJD130 | Two ~500bp DNA fragments for homologous integration upstream of *algI* on pSymB (HR1_algI_B and HR2_algI_B) were PCR amplified with primer pairs 360/361 and 362/363, respectively. Consitutive promoter P*min2* (Döhlemann *et a*l., 2017) was generated via hybridization of 388 and 389. A *loxL* site was generated via hybridzation of primers 187 and 188. Then, HR1_algI (digested with EcoRI/BamHI), P*min2* (BamHI/NdeI compatible 5‘ phosphorylated cohesive ends), *loxL* (NdeI/SalI compatible 5‘ phosphorylated cohesive ends) and HR2_algI (SalI/HindIII) were ligated into EcoRI/HindIII-opened pK18mobsacB. |
| pJD169 | A *tetO* array (120 copies) was released from EcoRV/HincII-digested pLAU44. In a next step a 500bp homologous region allowing for integration into the chromosomal terminus region was PCR amplified from *S. meliloti* gDNA with primers 510 and 511. Finally, the *tetO* array (SmaI) and the SalI/HindIII-digested PCR product (SalI/HindIII) were successive ligated into pK18mob2. |
| pJD170 | A *tetO* array (120 copies) was released from EcoRV/HincII-digested pLAU44. In a next step a 500bp homologous region allowing for integration into the pSymB terminus region was PCR amplified from *S. meliloti* gDNA with primers 514 and 515. Finally, the *tetO* array (SmaI) and the SalI/HindIII-digested PCR product (SalI/HindIII) were successive ligated into pK18mob2. |
| pJD201 | Two ~500bp homologous regions (HR1 and HR2) allowing for deletion of *repC2* were PCR amplified from *S. meliloti* gDNA using primer pairs 637/638 and 639/640, respectively. Then, HR1 (digested with EcoRI/NdeI) and HR2 (NdeI/HindIII) were ligated into EcoRI/HindIII-opened pK18mobsacB. |
| pJD202 | Two ~500bp homologous regions (HR1 and HR2) allowing for deletion of *repC1* were PCR amplified from *S. meliloti* gDNA using primer pairs 641/642 and 643/644, respectively. Then, HR1 (digested with EcoRI/NdeI) and HR2 (NdeI/HindIII) were ligated into EcoRI/HindIII-opened pK18mobsacB. |
| pJD206 | A ~500bp homologous region allowing for integration into *repA2* was PCR amplified from *S. meliloti* gDNA with primers 694 and 695. Then, the BamHI/HindIII-digested PCR product was ligated into BamHI/HindIII-opened pK18mob2. |
| pJD207 | A ~550bp homologous region allowing for integration into *repA1* was PCR amplified from *S. meliloti* gDNA with primers 696 and 697. Then, the BamHI/HindIII-digested PCR product was ligated into BamHI/HindIII-opened pK18mob2. |
| pJD222 | Two ~500bp homologous regions (HR1 and HR2) allowing for deletion of a spectinomycin resistance cassette in *S. meliloti* replicon fusion strains were PCR amplified from *S. meliloti* gDNA using primers 236/237 and 240/241, respectively. HR1 and HR2 were 5‘ phosphorylated and then digested with EcoRI and SphI, respectively, and ligated into EcoRI/SphI-opened pK18mobsacB. |
| pJD225 | Two ~500bp homologous regions (HR1 and HR2) allowing for deletion of *repA3B3* were PCR amplified from *S. meliloti* gDNA using primer pairs 787/788 and 789/790, respectively. Then, HR1 (digested with HindIII/NdeI) and HR2 (NdeI/EcoRI) were ligated into EcoRI/HindIII-opened pK18mobsacB. |
| pJD226 | Two ~500bp homologous regions (HR1 and HR2) allowing for deletion of the *hemE* promoter region were PCR amplified from *S. meliloti* gDNA using primer pairs 796/797 and 798/799, respectively. Then, HR1 (digested with EcoRI/NdeI) and HR2 (NdeI/HindIII) were ligated into EcoRI/HindIII-opened pK18mobsacB. |
| pJD227 | Two ~500bp homologous regions (HR1 and HR2) allowing for deletion of the entire *oriC* region were PCR amplified from *S. meliloti* gDNA using primer pairs 796/797 and 800/801, respectively. Then, HR1 (digested with EcoRI/NdeI) and HR2 (NdeI/HindIII) were ligated into EcoRI/HindIII-opened pK18mobsacB. |
| pJD228 | Two ~500bp homologous regions (HR1 and HR2) allowing for deletion of *oriC* derived DnaA boxes 2-4 were PCR amplified from *S. meliloti* gDNA using primer pairs 802/803 and 800/801, respectively. Then, HR1 (digested with EcoRI/NdeI) and HR2 (NdeI/HindIII) were ligated into EcoRI/HindIII-opened pK18mobsacB. |
| pJD229 | Two ~500bp homologous regions (HR1 and HR2) allowing for deletion of a gentamicin resistance cassette in *S. meliloti* replicon fusion strains were PCR amplified from *S. meliloti* gDNA using primers 360/361 and 364/791, respectively. HR1 and HR2 were 5‘ phosphorylated and then digested with EcoRI and PstI, respectively, and ligated into EcoRI/PstI-opened pK18mobsacB. |
| pJD266 | A ~500bp homologous region allowing for integration into *repA3* was PCR amplified from *S. meliloti* gDNA with primers 952 and 953. Then, the EcoRI/HindIII-digested PCR product was ligated into EcoRI/HindIII-opened pK18mob2. |

**S10 Table (2/2). Plasmid construction.**

| Construct | Construction |
|---|---|
| pMW186 | Homologous regions HR_SMa2383 and HR_SMa2385 (each ~1000bp) were PCR amplified from *S. meliloti* gDNA using primers 878/879 and 880/881, respectively. A *tetO* array (120 copies) was PCR amplified from pLAU44 using primers 837/838. A mini-MCS was generated by hybridization and 5‘-phosphorylation of primers 719/720 and inserted into BamHI/SphI-opened pK18mobsacB. The resulting construct was then succesive equipped with the *tetO* array (SphI/XbaI), HR_SMa2383 (NotI/SphI) and HR_SMa2385 (HindIII/XbaI). |
| pMW188 | Homologous regions HR_SMb20041 and HR_SMb20042 (each ~650bp) were PCR amplified from *S. meliloti* gDNA using primers 839/840 and 841/842, respectively. A *lacO* array (120 copies) was PCR amplified from pLAU43 using primers 835/836. A mini-MCS was generated by hybridization and 5‘-phosphorylation of primers 719/720. pK18mobsacB was then successive equipped with the Mini-MCS (BamHI/SphI), the *lacO* array (SphI/XbaI), HR_SMb20042 (XbaI/XhoI) and HR_SMb20041 (NotI/SphI). The HR_SMb20041-$lacO_{120}$-HR_SMb20042 fragment was released from resulting construct using NotI and PmeI, treated with Klenow enzyme in order to generate blunt ends and finally ligated (reverse orientation) into a new SmaI-opened pK18mobsacB backbone. |
| pMW189 | PCR amplification of ~500bp homologous regions HR1 and HR2 from *S. meliloti* gDNA with primers 266/267 and 330/269, respectively. HR1 and HR2 were then digested with EcoRI/SalI and SalI/SphI, respectively, and ligated into SalI/SphI-opened pK18mobsacB (pK18mobsacB/HR1-HR2). The coding region of *tetR* was PCR amplified from pLAU53 with primers tetR fwd and tetR rev. *mVenus* was PCR amplified from pG18mob-mVenus with primers mVenus fwd/ mVenus rev. Both PCR products were digested with MluI and ligated with each other. The ligation product was then PCR amplified with primers tetR fwd and mVenus rev, resulting in *tetR-mVENUS*. *lacI* was PCR amplified from pLAU53 with primers lacI fwd and lacI rev. *mCherry* was PCR amplified from pSRKGm-CmChr with primers mCherry fwd and mCherry rev. Both PCR products were digested with NotI and ligated with each other. The ligation product was then PCR amplified with primers lacI fwd and mCherry rev, resulting in *lacI-mCherry*. *tetR-mVENUS* (digested with SalI and SpeI) and *lacI-mCherry* (digested with SpeI and ScaI) were ligated into SalI/ScaI-opened pK18mobsacB/HR1-HR2. The resulting construct was digested with NotI and subsequently ligated with hybridized and 5’ phosphorylated primers 892/893. |
| pMW193 | A ~500bp homologous region (HR_terA) was PCR amplified from *S. meliloti* gDNA with primers 512/894. A *lacO* (120 copies) array was PCR amplified from pLAU43 with primers 835/836, 5‘ phosphorylated and inserted into ScaI-opened and dephosphorylated pLoriT. The resulting construct was ligated with HR_terA via digestion of insert and vector with EcoRI and KpnI. |
| pMW198 | A 3´ portion of *parB* was PCR amplified from *S. meliloti* gDNA with primers parB+1-Nde and parB+888-r-Xba. *cerulean* was PCR amplified from $pWBT5^{cerulean}$ with primers EGFP/mCh-f-Xba and mCh/egfp-Hind-r. Then *parB* (digested with NdeI/XbaI) and *cerulean* (digested with XbaI/HindIII) were ligated into NdeI/HindIII-opened pSRKKm. The resulting construct was digested with HindIII, treated with Klenow fragment for generation of blunt ends, and finally ligated with a ~500bp homologous downstream region of *parB* (HR_parB_down) which was PCR amplified from *S. meliloti* gDNA with primers ParB-down fwd and ParB-down rev. The ligation product served as template for PCR amplification of *parB-cerulean-HR_parB_down* with primers ParBHR1-cerulean fwd and ParB-down rev , and the 5‘ phosphorlylated PCR product was finally inserted into SmaI-opened pK18mobsacB. |
| pMW210 | Two ~500bp homologous regions (HR1 and HR2) allowing for deletion of *repA1B1C1* in *S. meliloti* were PCR amplified from *S. meliloti* gDNA using primers 822/823 and 828/829, respectively. Then, HR1 (digested with HindIII/PstI) and HR2 (PstI) were ligated into HindIII/SmaI-opened pK18mobsacB. |
| pMW211 | Two ~500bp homologous regions (HR1 and HR2) allowing for deletion of *repA2B2C2* in *S. meliloti* were PCR amplified from *S. meliloti* gDNA using primers 814/815 and 820/821, respectively. Then, HR1 (digested with HindIII/PstI) and HR2 (PstI) were ligated into HindIII/SmaI-opened pK18mobsacB. |
| pMW212 (pK18mob3) | A 408 bp region containing the promoter sequence of $P_{lac}$ and parts of the *lacZα* fragment was released from AseI/NheI-digested pK18mob2 and replaced by a smaller MCS fragment (amplified from pK18mob2 using primer -48/-67 after treatment of the vector backbone with Klenow fragment. |
| pMW230 | Two ~500bp homologous regions (HR1 and HR2) allowing for deletion of the *parS* sequences upstrem of *repA2* on pSymA were PCR amplified from *S. meliloti* gDNA using primers 1039/1040 and 820/821, respectively. Then, HR1 (digested with EcoRI/PstI) and HR2 (PstI/HindIII) were ligated into EcoRI/HindIII-opened pK18mobsacB. |
| pMW256 | Two fragments representing the N- and C-terminal part of *cckA* were PCR amplified from *S. meliloti* gDNA using modified primer 1064/1066 and 1068/1069, respectively enabling introduction of the C→T transition at bp 1307 in the *cckA* sequence. Subsequent, overlapping fragments were then used in a overlap extension PCR to generate a full length sequence of mutagenized *cckA*. This fragment was introduced via EcoRI and HindIII into pK18mobsacB. |
| pMW257 | Sequence of *cckA* was PCR amplified from *S. meliloti* gDNA using primer 1064/1069. Resulting fragment was digested with EcoRI and HindIII and introduced into EcoRI/HindIII opened pK18mobsacB. |
| pMW261 | A fragment containing *parB-cerulean* was PCR amplified from pMW198 using primer ParBHR1-cerulean fwd and ParBHR1-cerulean rev. Resulting amplicon was digested with PstI and introduced into SmaI/PstI opened pMW212. |
| pFROS | P*tau-tetR-mVenus-lacI-mCherry* was PCR amplified from pMW189 with primers Ptau Fwd + KpnI and 890. A *mob* site was PCR amplified from pK18mob2 with primers 331 and 332. pYEM1 was digested with SpeI, treated with Klenow enzyme in order to generate blunt ends, and finally ligated with the 5‘ phosphorylated *mob* fragment. The resulting construct was digested with KpnI and PacI and ligated with P*tau-tetR-mVenus-lacI-mCherry* (digested with KpnI/PacI). |

**S11 Table (1/5). Oligonucleotides used in this study.**

| Number/ Name | Sequence (5‘-3‘) | Used for |
|---|---|---|
| EGFP/mCh-f-Xba | GAGTCTAGAATGGTGAGCAAGGGCGAGGAG | PCR amplification of *cerulean* |
| mCh/egfp-Hind-r | GTACAAGCTTTTACTTGTACAGCTCGTCCATG | |
| parB+1-Nde | GAACCATATGAACGACGACAGCTCCAAG | PCR amplification of *parB* |
| parB+888-r-Xba | GTACAAGCTTTTACTTGTACAGCTCGTCCATG | |
| tetR fwd | ATATGTCGACTCTAGCAGGAGGAATTCACCAT | PCR amplification of *tetR* |
| tetR rev | ATATACGCGTGTATGTCAGACCCACTTTCACATT | |
| mVenus fwd | ATATACGCGTTGGTGAGCAAGGGCGAGGA | PCR amplification of *mVenus* |
| mVenus rev | ATATACTAGTTTACTTGTACAGCTCGTCCATGC | |
| lacI fwd | ATATACTAGTTTTGGGCTAGCAGGAGGAATT | PCR amplification of *lacI* |
| lacI rev | ATATGCGGCCGCCAGCTGCATTAATGAATCGG | |
| mCherry fwd | ATATGCGGCCGCTGGTGAGCAAGGGCGAGGA | PCR amplification of *mCherry* |
| mCherry rev | ATATAGTACTTTACTTGTACAGCTCGTCCATGC | |
| ParBHR1-cerulean fwd | ATATCTGCAGATCGTCGAGAATGTTCAGCG | PCR amplification of *parB-cerulean* |
| ParBHR1-cerulean rev | ATATACGCGTTTACTTGTACAGCTCGTCCATGC | |
| ParB-down fwd | CGCACGCGTTACAAACTCAACGCCTCGC | PCR amplification of HR_parB_down |
| ParB-down rev | GCGAAGCTTGGCTTCGCGCAACGAGCTT | |
| Ptau fwd + KpnI | CGCGGTACCAATTCGAAGGTCGAGCGCAA | PCR amplification of P*tau-tetR-mVenus-lacI-mCherry* |
| -48 | AGCGGATAACAATTTCACACAGGA | PCR amplification of the pK18mob2 MCS/ Sequencing of pK18mobsacB/2 derivatives |
| -67 | AAGGCGATTAAGTTGGGTAACG | PCR amplification of the pK18mob2 MCS/ Sequencing of pK18mobsacB/2 derivatives |
| 6 | CATCGGCAATGTTGAATGG | PCR amplification of $P_{aacC1}$ |
| 7 | CGCGGATCCCCTGACGATGCGTGGAGACC | |
| 125 | GCGGTCGACGTTATGGAGCAGCAACGATGTT | PCR amplification of gentamicin resistance cassette *aacC1* |
| 126 | GCGCTGCAGAGTACT TTAGGTGGCGGTACTTGGGTCG | |
| 127 | CGCGTCGACGGACTCAATACACCATGAGGG | PCR amplification of spectinomycin resistance cassette *aadA1* |
| 128 | GCGCTGCAGAGTACT TTATTTGCCGACTACCTTGGTG | |

**S11 Table (2/5). Oligonucleotides used in this study.**

| Number/ Name | Sequence (5'-3') | Used for |
|---|---|---|
| 131 | CGCGGATCCCTGACGCGTACAGGAAACAC | PCR amplification of the *thrA-rpoC* terminator sequence, Transition zone validation (131) |
| 132 | GCGCATATGCGGGCTCTTTCCCTAAACTCC | |
| 187 | TATGTACCGTTCGTATAGCATACATTATACGAAGTTATG | Generation of *loxL* |
| 188 | TCGACATAACTTCGTATAATGTATGCTATACGAACGGTACA | |
| 189 | TATGATAACTTCGTATAGCATACATTATACGAACGGTAG | Generation of *loxR* |
| 190 | TCGACTACCGTTCGTATAATGTATGCTATACGAAGTTATCA | |
| 236 | CGCGAATTCTGGAGACCGGGTCAAGCTG | PCR amplification of a ~500bp region for homologous integration upstream of *nodP1* (HR1_nodP1), Transition zone validation (236) |
| 237 | CGCGGATCCCGACTTCGGTACCGGTGCC | |
| 238 | GCGGTCGACCGTCCGCCGTTGCGTATCTC | PCR amplification of a ~500bp region for homologous integration upstream of *nodP1* (HR2_nodP1), Transition zone validation (239) |
| 239 | CGCGCATGCCTATCCGGGTTACGCCTCGG | |
| 240 | CGCCCCTTTGCGAAATGCT | PCR amplification of a ~500bp region for homologous integration downstream of *nodQ2* (HR_nodQ2) |
| 241 | CGCGCATGCTGGCAACCGCGGTCAGTTC | |
| 247 | CGGGATCCTTAGGTGGCGGTACTTGGGTC | Transition zone validation |
| 249 | CGGGATCCTTATTTGCCGACTACCTTGGTGATCT | |
| 266 | GCGGAATTCGAAGGTCGAGCGCAACTTG | PCR amplification of a ~500bp region (HR1) used for integration into the *tau* promoter region |
| 267 | GCGGTCGACCGTTACCCTCTTTGGTTATGTC | |
| 269 | GCGGCATGCGTTGAGGATGTTCACTTCC | PCR amplification of a ~500bp region (HR2) used for integration into the *tau* promoter region |
| 330 | GCGAGTACTACGGTCGAGCCCGCCATTGTTC | |
| 331 | CTCGCGGACGTGCTCATAG | PCR amplification of a *mob* site |
| 332 | GCTTTTCCGCTGCATAACCC | |
| 360 | CGCGAATTCTGAAAATAGCGGTATAGCGATGTGGC | PCR amplification of a ~500bp region for homologous integration upstream of *algI* (HR1_algI_B), Transition zone validation (360) |
| 361 | GCGGGATCCCCGCTTGCGCGTCTTCTCG | |
| 362 | CGCCTGCAGGTGCTGGTCGCCGGCCAGG | PCR amplification of a ~500bp region for homologous integration upstream of *algI* (HR2_algI_B), Transition zone validation (363) |
| 363 | GCGAAGCTTGCGGATCGAAAGTGATAACTGCCATAG | |
| 364 | GAAAGTGGATTCCGAGTCTTC | PCR amplification of a ~500bp region for homologous integration downstream of the chromosomal *algI* homolog (HR_algI_Chr) |
| 365 | TCCAATCTCCGTTTGTCGTTG | |

**S11 Table (3/5). Oligonucleotides used in this study.**

| Number/ Name | Sequence (5'-3') | Used for |
|---|---|---|
| 388 | GATCCTGTTGACAATTAATCATCGAACTAGTTAACTAGTACGCAAGTAGACGCGTTACGTCA | Generation of P*min2* |
| 389 | TATGACGTAACGCGTCTACTTGCGTACTAGTTAACTAGTTCGATGATTAATTGTCAACAG | |
| 510 | CGCGTCGACGATTATCGGCGTCGAAACAGGC | PCR amplification of a ~500bp region for homologous insertion at *terC* |
| 511 | GCGAAGCTTAAGAACCAGGGCACCGGTTC | |
| 512 | CGCGAATTCGGATTGTCGGCCTGCTCAAG | PCR amplification of a ~500bp region (HR_terA) for homologous insertion at *terA* |
| 514 | CGCGTCGACGATGATCAACACCGCGCGC | PCR amplification of a ~500bp region for homologous insertion at *terB* |
| 515 | GCGAAGCTTGAGGATCGCGGCTTCGACG | |
| 637 | CGCGAATTCCGAAACCTCCTTGGATCACG | PCR amplification of a ~500bp region (HR1) for *repC2* deletion |
| 638 | GCGCATATGGATCCCACTCAAGCGGAATC | |
| 639 | CGCCATATGTTAGTCTCCTTGTTTCGAACTCTG | PCR amplification of a ~500bp region (HR2) for *repC2* deletion |
| 640 | GCGAAGCTTCGGGAACTTAGCGACGAAGA | |
| 641 | CGCGAATTCGCGTTTCGTTCTGCATTTCC | PCR amplification of a ~500bp region (HR1) for *repC1* deletion |
| 642 | GCGCATATGTGTCTCCGCTGAACAGTCTTTG | |
| 643 | CGCCATATGTTAGTCTCCTGTTTCCCGTTGATA | PCR amplification of a ~500bp region (HR2) for *repC1* deletion |
| 644 | GCGAAGCTTACGCAATCGTCGAACTGGAC | |
| 694 | GCGGGATCCGAGAAAAGTGTATCGCCGCCG | PCR amplification of a ~500bp region for homologous inseration into *repA2* |
| 695 | CGCAAGCTTGACGCGACACCTAAGCAGC | |
| 696 | GCGGGATCCCTCGTAGCGTATGGCGCCAT | PCR amplification of a ~500bp region for homologous inseration into *repA1* |
| 697 | CGCAAGCTTGTTGCAGCAGAGAATCCAGAGTC | |
| 719 | GATCCCGCGCGGCCGCCGCGCATG | Generation of a mini-MCS |
| 720 | CGCGGCGGCCGCGCGG | |
| 787 | CGCAAGCTTCGCCATGTTTATTCTTTCCGG | PCR amplification of a ~500bp region (HR1) for *repA3B3* deletion |
| 788 | GCGCATATGATCCATGGTCAAATCTCCTCGTG | |
| 789 | CGCCATATGCGTGGAGAATCAGAAGCCTTAGAG | PCR amplification of a ~500bp region (HR2) for *repA3B3* deletion |
| 790 | GCGGAATTCATGACGAATTTCCTGCCCGC | |
| 791 | GCGCTGCAGTCCAATCTCCGTTTGTCGTTG | PCR amplification of a ~500bp region used for deletion of a gentamicin resistance cassette in *S. meliloti* replicon fusion strains |

**S11 Table (4/5). Oligonucleotides used in this study.**

| Number/ Name | Sequence (5'-3') | Used for |
|---|---|---|
| 796 | CGCGAATTCACCAGATAGTCGGCGGAAATG | PCR amplification of a ~500bp region (HR1) used for deletion of *oriC* and subregions |
| 797 | CGCCATATGACAAGCGGGACGCTATCTC | |
| 798 | CGCCATATGTTTGCGGTTCCTGTGGATG | PCR amplification of a ~500bp region (HR2) used for deletion of the *hemE* promoter region |
| 799 | CGCAAGCTTATTCGGCATCCATGACATGC | |
| 800 | CGCCATATGGTGGAGAACCGCAAAAACTAC | PCR amplification of a ~500bp region (HR2) used for deletion of the entire *oriC* region |
| 801 | GCGAAGCTTGCGAGATAGATGCTCGTCG | |
| 802 | CGCGAATTCTCGATCGGATCCATCCGTG | PCR amplification of a ~500bp region (HR1) used for deletion of *oriC* derived DnaA boxes 2-4 |
| 803 | CGCCATATGGAACCGCAAACTCCCGATC | |
| 814 | CGCAAGCTTCCGCGTCGATGCCGGGCAG | PCR amplification of a ~500bp region (HR1) used for deletion of *repA2B2C2* |
| 815 | GCGCTGCAGCAAATTCGTCGGTCGCCGCACTTCGTAATCAGCG | |
| 820 | CGCCTGCAGCTGGATGCCCGCCTCCGC | PCR amplification of a ~500bp region (HR2) used for deletion of *repA2B2C2* or *parS* sequences |
| 821 | GCGAAGCTTATACGCTCGAACTTATCCGAGCCCGGC | |
| 822 | CGCAAGCTTGTCCCGGAGAGCCGAAGT | PCR amplification of a ~500bp region (HR1) used for deletion of *repA1B1C1* |
| 823 | GCGCTGCAGGAACAGATGTGAGCGAAGCACAATAGG | |
| 828 | CGCCTGCAGCCCCCAGACGTTGCAACAGA | PCR amplification of a ~500bp region (HR2) used for deletion of *repA1B1C1* |
| 829 | GCGAAGCTTAGAATTCTCTGATGCAGCGACAGG | |
| 835 | CGCGCATGCAAAAAGCAATGCGGACCTTG | PCR amplification of a *lacO* array |
| 836 | GCGTCTAGAGTAACATCAGCTAGAGCAGGT | |
| 837 | CGCGCATGCCACAGGAAACAGCTATGACCATGA | PCR amplification of the *tetO* array |
| 838 | GCGTCTAGATTTACGAACCGAACAGGCGC | |
| 839 | AAATATGCGGCCGCGCAGCGATCCGCCGTTTCAT | PCR amplification of a ~650bp region (HR_SMb20041) for homologous insertion at *oriB* |
| 840 | GCGGCATGCGGGAACGACCGCTTCAGCT | |
| 841 | CGCTCTAGACTCGCTCTTTCAGAGCACGCC | PCR amplification of a ~650bp region (HR_SMb20042) for homologous insertion at *oriB* |
| 842 | GCGCTCGAGGAAGCGCGGTCATGGCAGAAC | |
| 850 | GATACAAGGCGCTGCAACC | PCR amplification of the *repC2* deletion site |
| 851 | TCAGATCGTATTTGGCCGG | |

**S11 Table (5/5). Oligonucleotides used in this study.**

| Number/ Name | Sequence (5‘-3‘) | Used for |
|---|---|---|
| 852 | CAAAAGGTGGCCGACGAATA | PCR amplification of the *repC1* deletion site |
| 853 | CCTCTCGAACATCACCCAG | |
| 878 | AAATATGCGGCCGCATGAAAGCCGTTGTGATGAAAGAGG | PCR amplification of a ~1000bp region (HR_SMa2383) for homologous insertion at *oriA* |
| 879 | GCGGCATGCCGCTCCGAGATATAGTTCATGCG | |
| 880 | CGCAAGCTTCGCGTCTGGTACCCGAAAAGC | PCR amplification of a ~1000bp region (HR_SMa2385) for homologous insertion at *oriA* |
| 881 | GCGTCTAGACCAACCGACACGGGCATGA | |
| 890 | CGCTTAATTAAGCTTGAAACTTCTAAGGTGGATCATGTC | PCR amplification of P*tau-tetR-mVenus-lacI-mCherry* |
| 892 | GGCCTGTTTAAACT | Linker for in frame fusion of *lacI* with *mCherry* |
| 893 | GGCCAGTTTAAACA | |
| 894 | GCGGGTACCGCTGGGCTGAGGTTCGAATTTC | PCR amplification of a ~500bp region (HR_terA) for homologous insertion at *terA* |
| 952 | CGCAAGCTTGACCTATTTCGACGGGCTCG | PCR amplification of a ~500bp region for homologous inseration into *repA3* |
| 953 | GCGGAATTCGCTTGGTGAGACCCGCATC | |
| 1039 | CGCCTGCAGAGAGTTCGGCAGATCTCCATTG | PCR amplification of a ~500bp region (HR1) used for deletion of *parS* sequences upstream of *repA2B2C2* |
| 1040 | GCGGAATTCAACCTGGCGGATATAGGCCT | |
| 1064 | ATATGAATTCGCGCCGCATAGTTCAGAGGAG | PCR amplification of wt *cckA* and *cckA* with C→T (bp 1307) transition |
| 1066 | AGGATGAGGGGCGTCACTTCCACTTCTATGTCAATGCAGTCA | PCR amplification of *cckA* with C→T (bp 1307) transition |
| 1068 | TGGAAGTGACGCCCCTCATCC | PCR amplification of *cckA* with C→T (bp 1307) transition |
| 1069 | ATATAAGCTTTCGGGATCGCGTTCGATGCGA | PCR amplification of wt *cckA* and *cckA* with C→T (bp 1307) transition |

**S1 Data. Validation of antibiotic marker deletion and genome configuration in *S. meliloti* fusion strains used for microscopy (A)** Deletion of spectinomycin resistance cassette ($P_{min2}$-*aadA1*) and gentamicin resistance cassette ($P_{min2}$-*aacC1*) together with the remaining *loxLR* sites from SmAB, SmABC and SmABCΔoriV leads to fusion strain variants SmABΔR, SmABCΔR and SmABCΔoriVΔR**. i)** Proper removal of SpecR in the respective strains was realized using construct pJD222 and verified by PCR with primer 792/793 (precursor strains: 2.32 kb, ΔSpecR: 1.16 kb). **ii)** Construct pJD229 mediated deletion of GmR was confirmed using primer 794/795 (precursor strains: 1.89 kb, ΔGmR: 1.24 kb). Pulsed-field gel electrophoresis banding pattern of PacI-digested gDNA of **(B)** SmABΔR, SmABCΔR and SmABCΔoriVΔR subsequent after antibiotic marker deletion, **(C)** SmCreΔhsdR (wt) and SmABCΔR with *parB::cerulean* after integration of $tetO_{120}$ array at loci 18, 17, 16, 5 or a $lacO_{120}$ array at loci 2, 3, 10, 13, 6, 15, 14, 9. (S11A Fig) and **(D)** SmCreΔhsdR (wt), SmABΔR, SmABCΔR and SmABCΔoriVΔR with simultaneously tagged *oriC*/*terC* (1), *oriC*/*oriA*/*oriB* (2), *oriC*/*oriA*/*terA* (3) and *oriC*/*oriB*/*terB* (4) regions via *parB::cerulean*, *tetO* or *lacO* array. Expected fragment sizes for SmCreΔhsdR (wt) derivatives (3.65, 1.35, 1.15, 0.53 Mbp), SmABΔR derivatives (3.65, 1.67, 0.83, 0.53 Mbp), SmABCΔR derivatives (2.53, 1.94, 1.67, 0.53 Mbp), and SmABCΔoriVΔR (2.53, 1.94, 1.67, 0.53 Mbp). PFGE with 0.5 x TBE in a 0.7 % agarose gel, separation 72 h. M: PFGE marker *S. cerevisiae*, Bio-Rad.

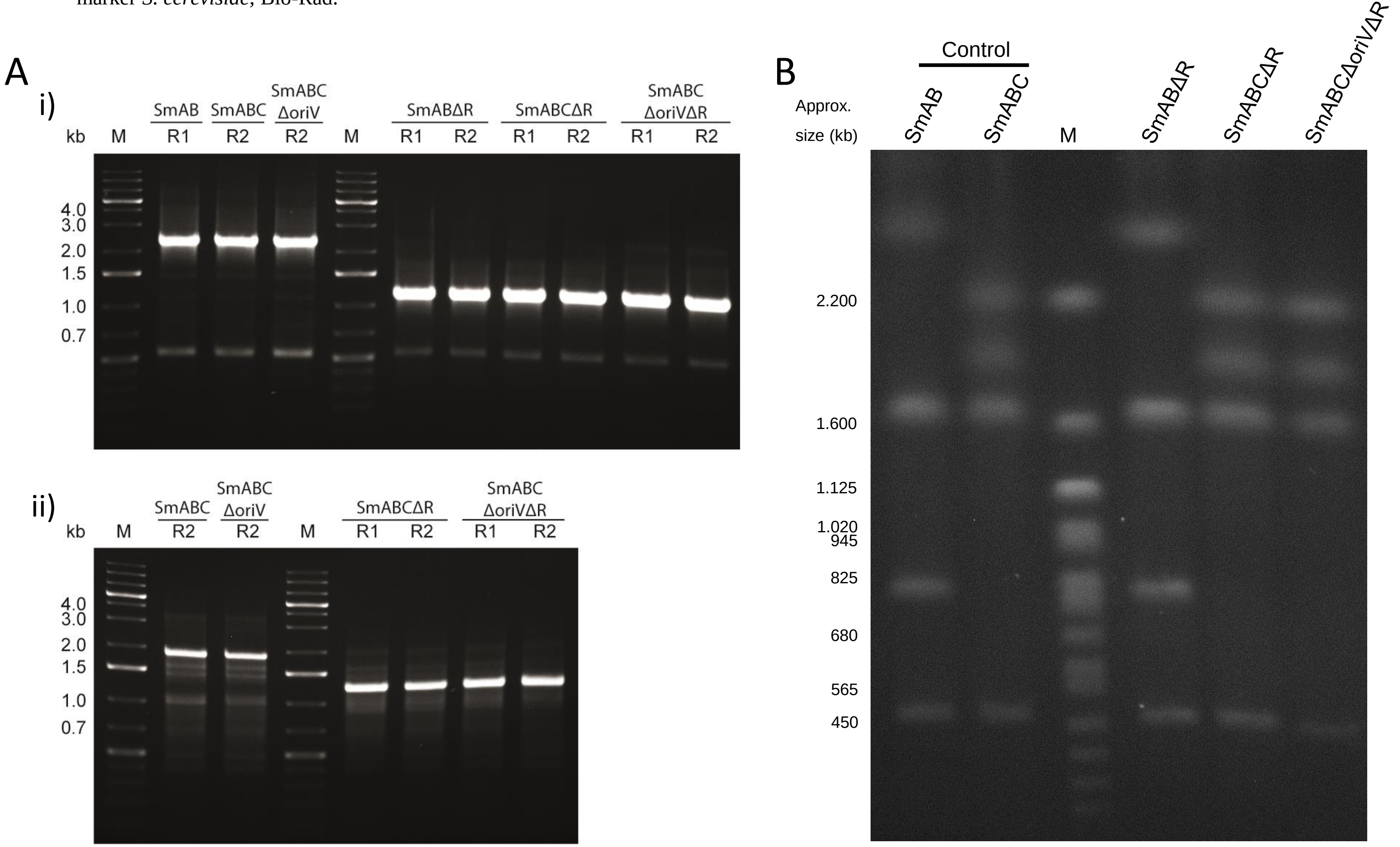

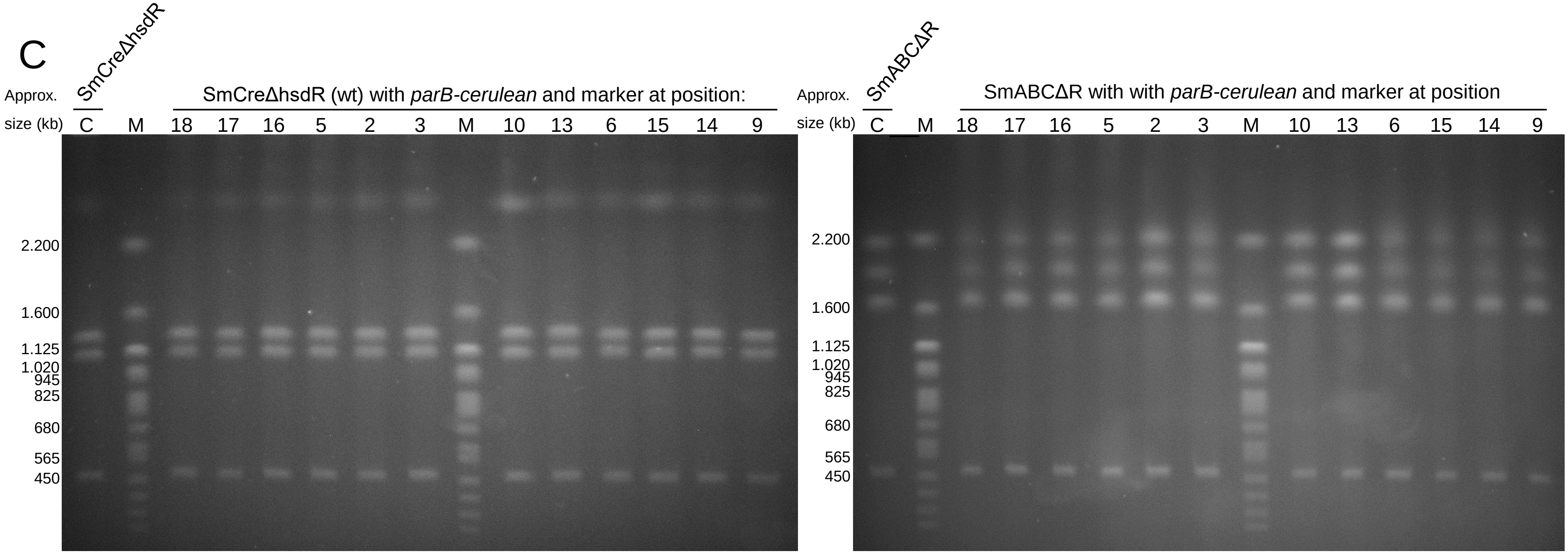
C
SmCreΔhsdR
Approx.
size (kb)
SmCreΔhsdR (wt) with parB-cerulean and marker at position:
C M 18 17 16 5 2 3 M 10 13 6 15 14 9
2.200
1.600
1.125
1.020
945
825
680
565
450
SmABCΔR
SmABCΔR with with parB-cerulean and marker at position
Approx.
size (kb)
C M 18 17 16 5 2 3 M 10 13 6 15 14 9
2.200
1.600
1.125
1.020
945
825
680
565
450

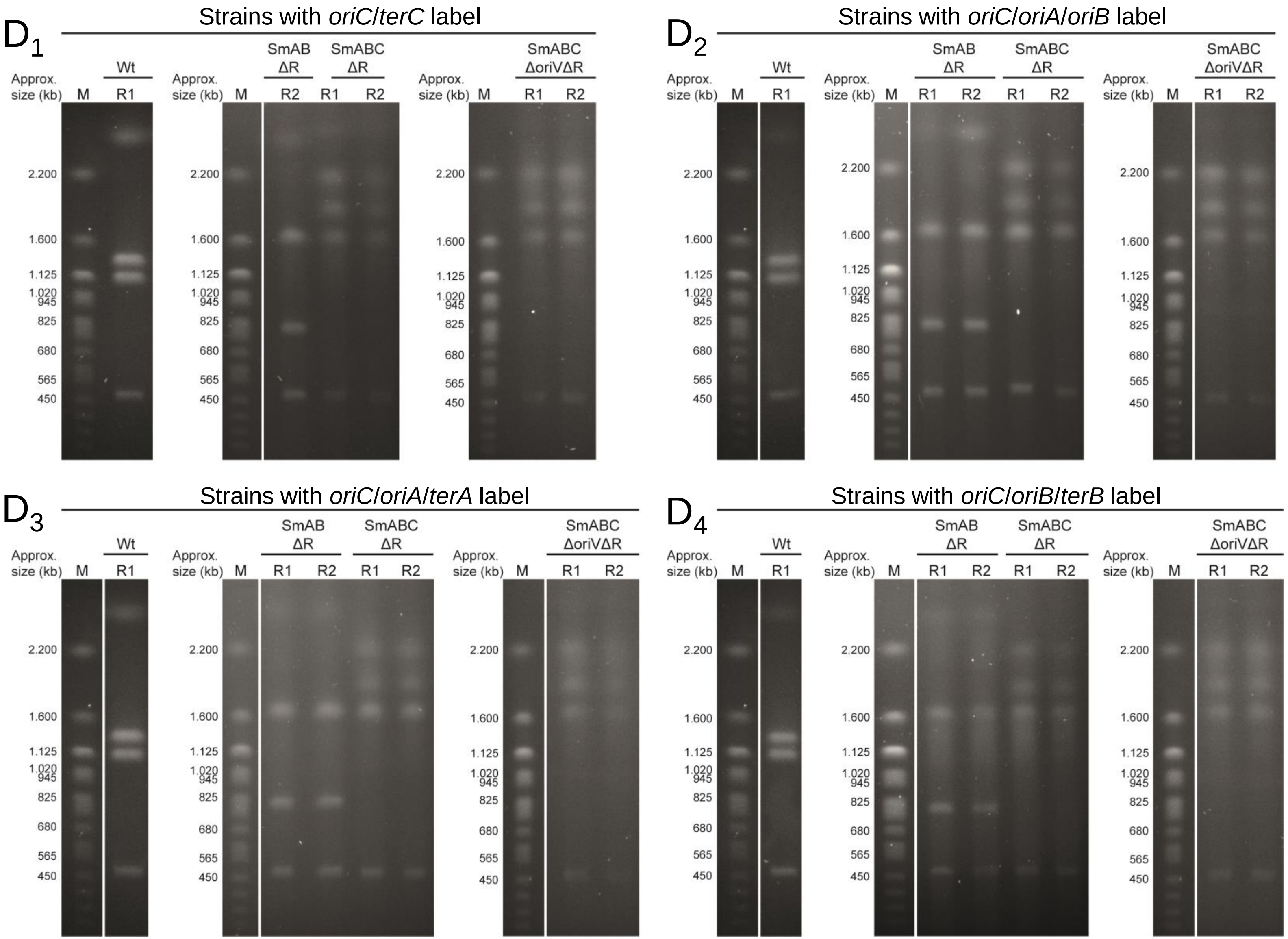
D1
Strains with oriC/terC label
Wt
SmAB ΔR
SmABC ΔR
SmABC ΔoriVΔR
Approx. size (kb)
M
R1
R2
2.200
1.600
1.125
1.020
945
825
680
565
450
D2
Strains with oriC/oriA/oriB label
D3
Strains with oriC/oriA/terA label
D4
Strains with oriC/oriB/terB label

**S2 Data. Visualization of single cell time-lapse data in SmCreΔhsdR (A), SmABΔR (B), SmABCΔR (C) and SmABCΔoriVΔR (D) during the cell cycle.** Displayed are the individual trajectories of simultaneously tagged *oriC*/*oriA*/*oriB* **(1)**, *oriC*/*terC* **(2)**, *oriC*/*oriA*/*terA* **(3)** and *oriC*/*oriB*/*terB* **(4)** regions in mother cells (M, upper plots) and daughter cells (D, lower plots). Shown are the relative positions within the cell with old pole at 0 and new pole at 1 over a time course of 200 min. Color code: chromosome (grey), pSymA (green), pSymB (blue). Origins are represented as circles and terminus regions as diamonds.

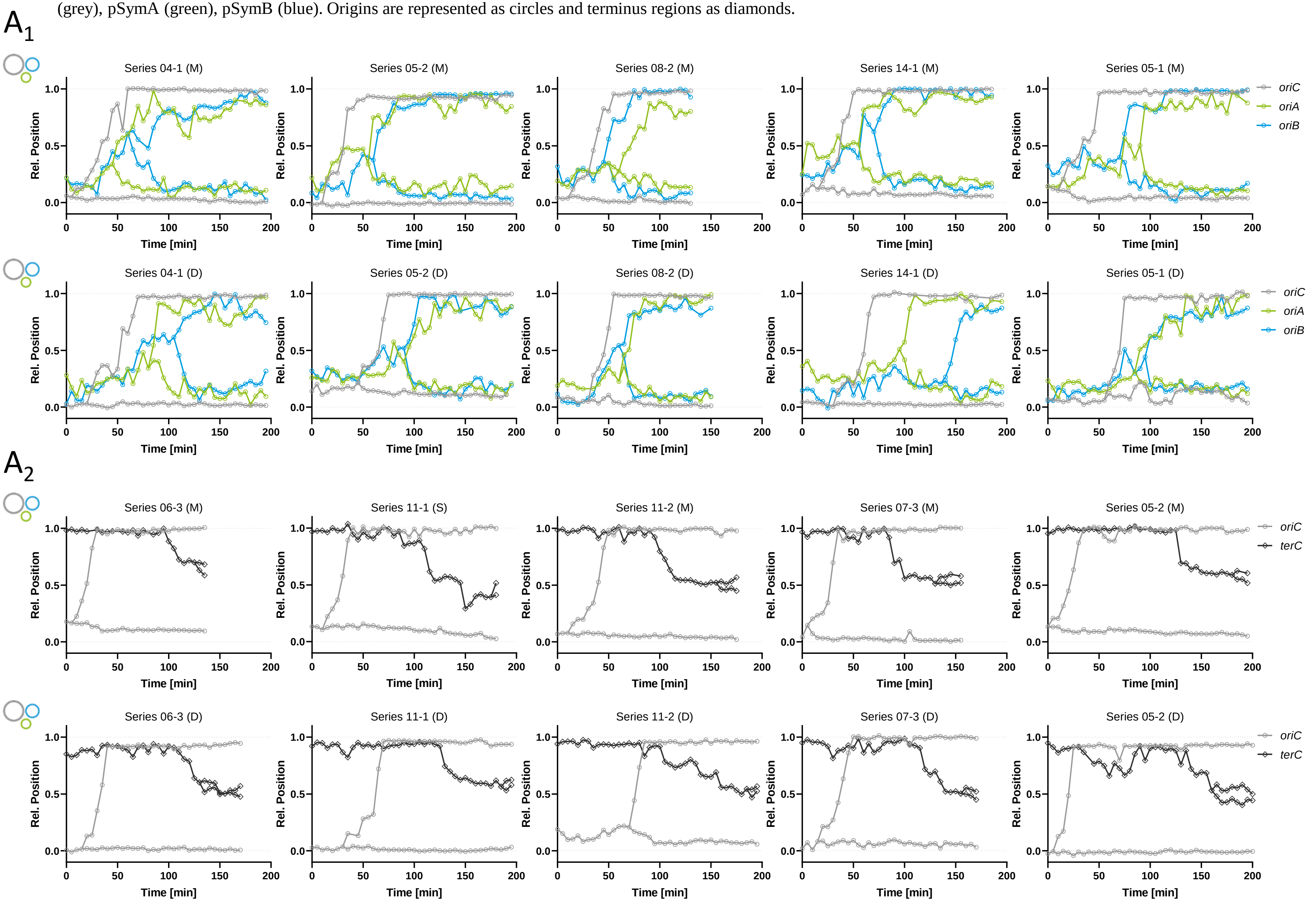

$A_3$

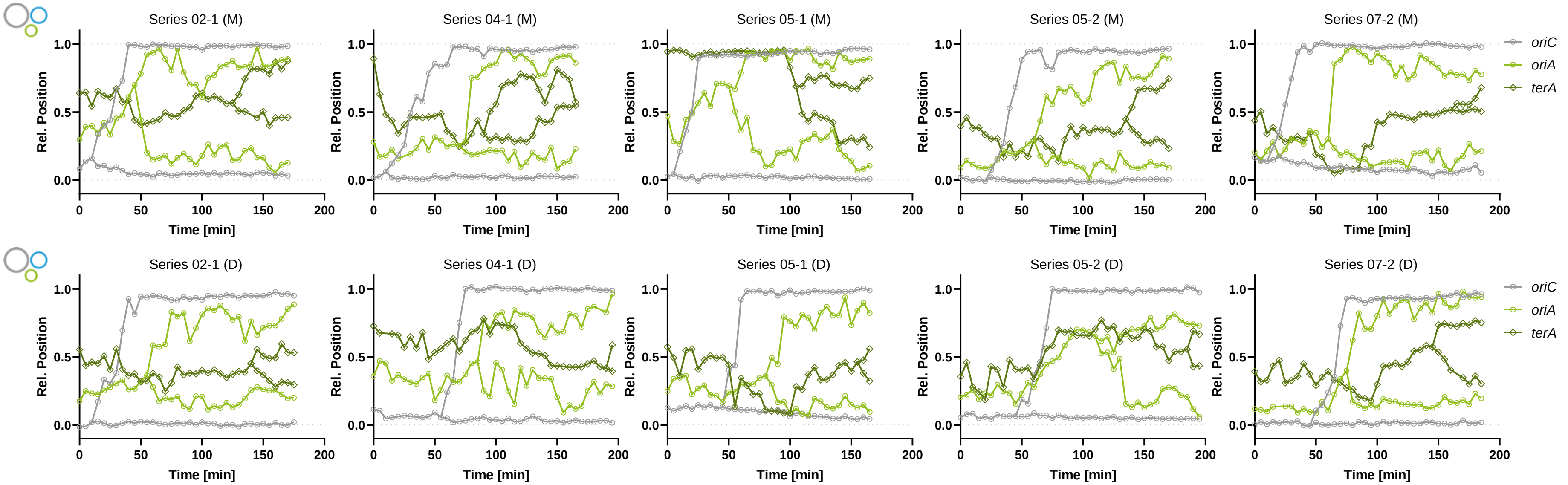
Series 02-1 (M)
Series 04-1 (M)
Series 05-1 (M)
Series 05-2 (M)
Series 07-2 (M)
Series 02-1 (D)
Series 04-1 (D)
Series 05-1 (D)
Series 05-2 (D)
Series 07-2 (D)
Rel. Position
Time [min]
0.0
0.5
1.0
0
50
100
150
200
oriC
oriA
terA

$A_4$

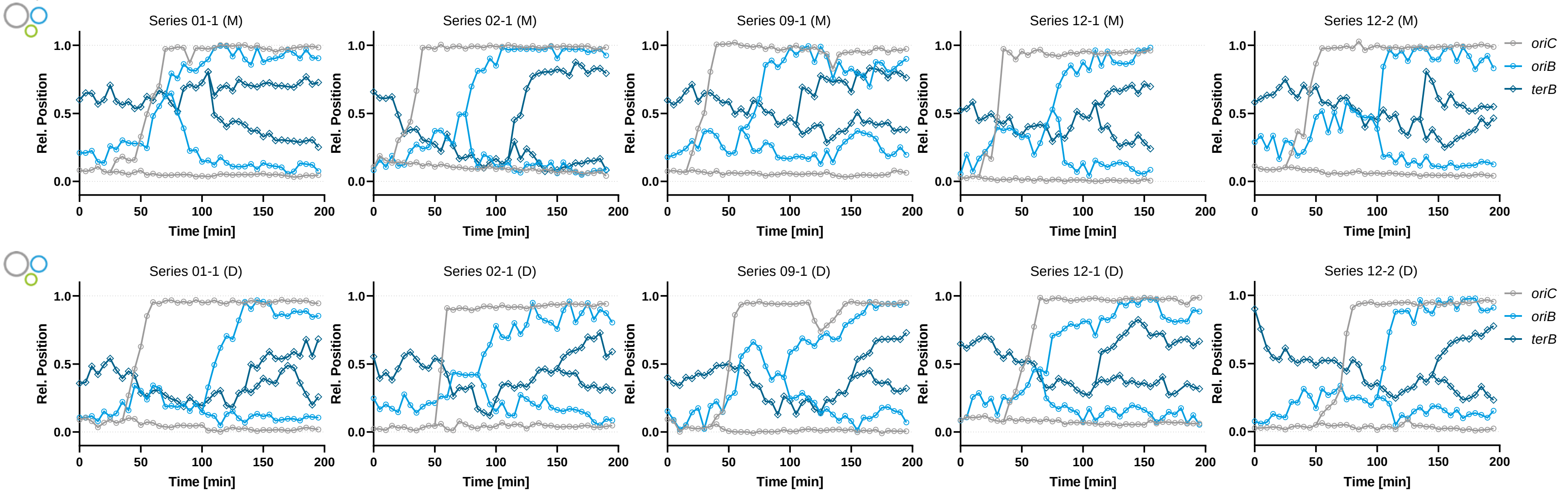
Series 01-1 (M)
Series 02-1 (M)
Series 09-1 (M)
Series 12-1 (M)
Series 12-2 (M)
Series 01-1 (D)
Series 02-1 (D)
Series 09-1 (D)
Series 12-1 (D)
Series 12-2 (D)
Rel. Position
Time [min]
0.0
0.5
1.0
0
50
100
150
200
oriC
oriB
terB

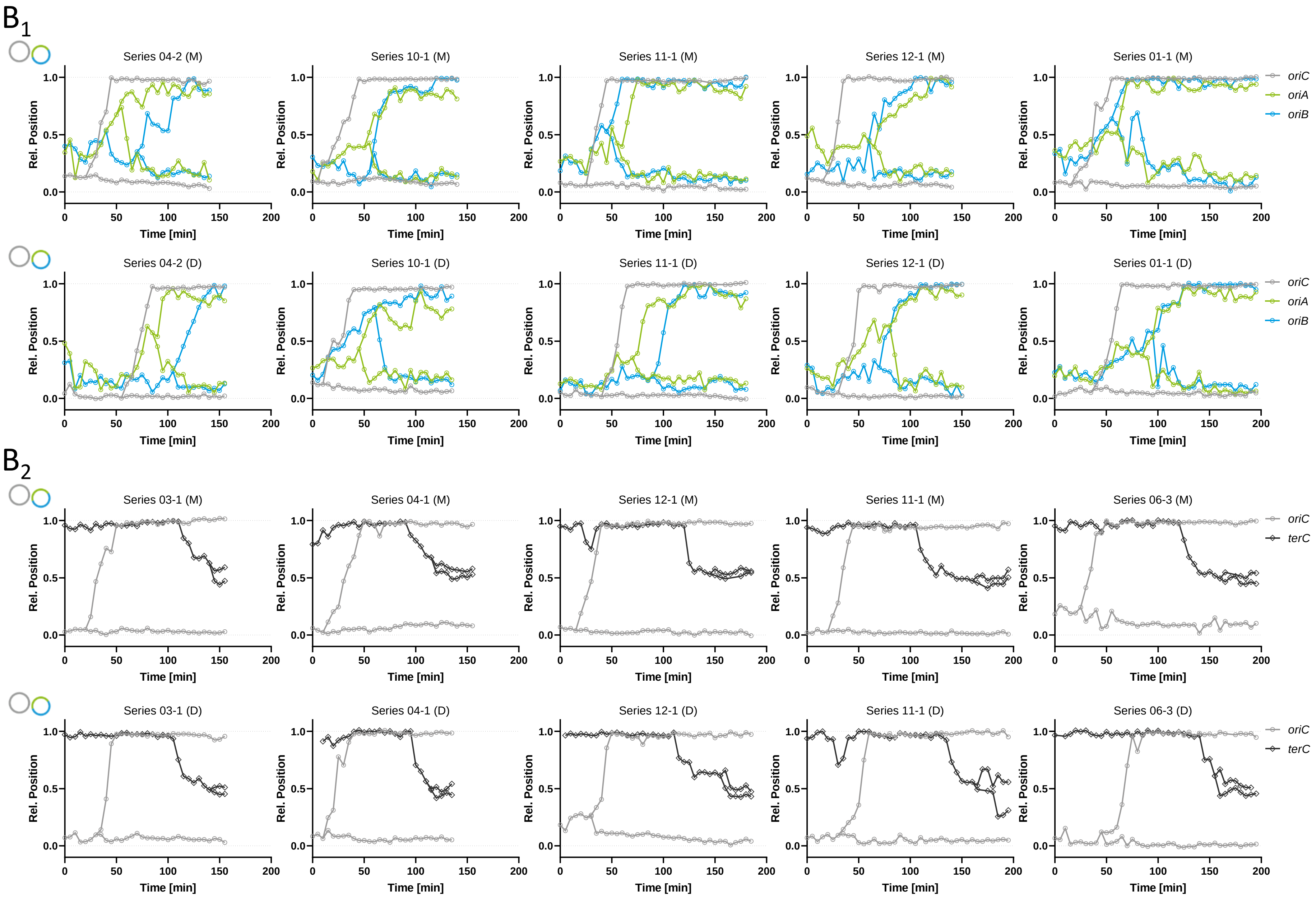
B1
Series 04-2 (M)
Series 10-1 (M)
Series 11-1 (M)
Series 12-1 (M)
Series 01-1 (M)
Series 04-2 (D)
Series 10-1 (D)
Series 11-1 (D)
Series 12-1 (D)
Series 01-1 (D)
oriC
oriA
oriB
B2
Series 03-1 (M)
Series 04-1 (M)
Series 12-1 (M)
Series 11-1 (M)
Series 06-3 (M)
Series 03-1 (D)
Series 04-1 (D)
Series 12-1 (D)
Series 11-1 (D)
Series 06-3 (D)
oriC
terC
Rel. Position
Time [min]

$B_3$

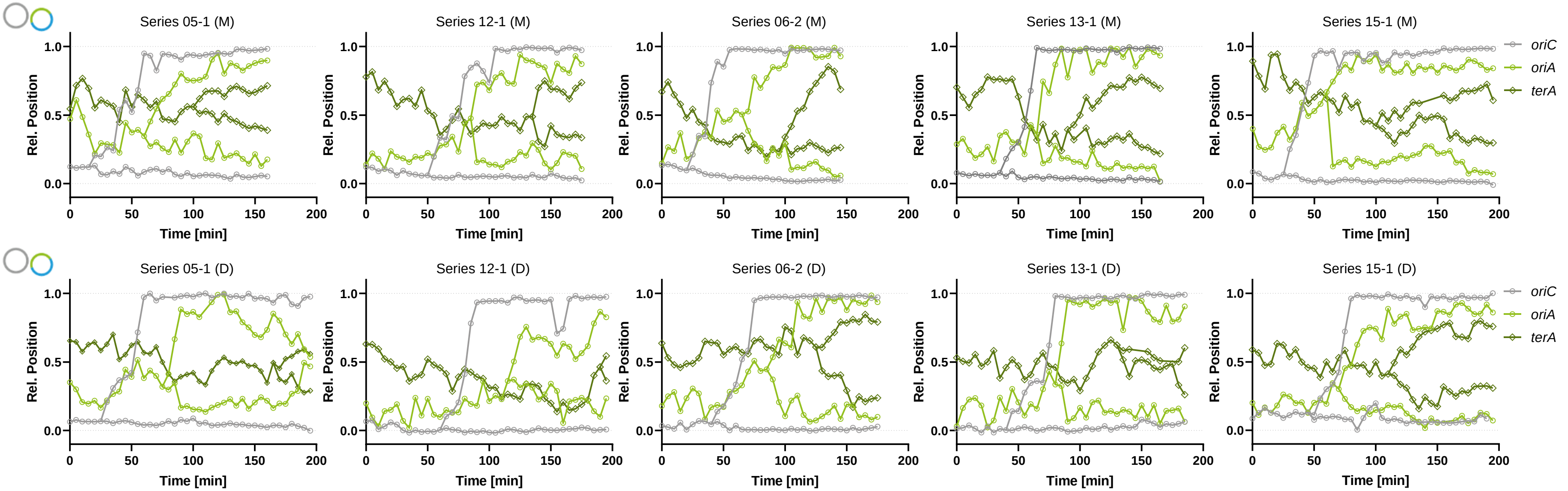
Series 05-1 (M)
Series 12-1 (M)
Series 06-2 (M)
Series 13-1 (M)
Series 15-1 (M)
Series 05-1 (D)
Series 12-1 (D)
Series 06-2 (D)
Series 13-1 (D)
Series 15-1 (D)
Rel. Position
Time [min]
oriC
oriA
terA

$B_4$

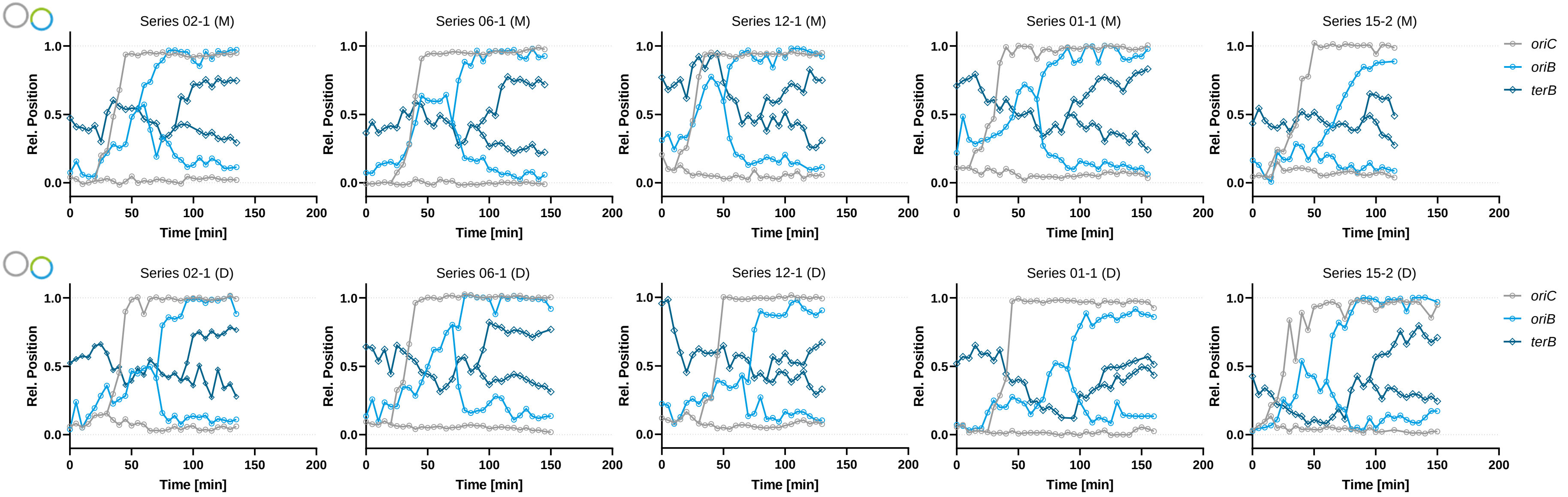
Series 02-1 (M)
Series 06-1 (M)
Series 12-1 (M)
Series 01-1 (M)
Series 15-2 (M)
Series 02-1 (D)
Series 06-1 (D)
Series 12-1 (D)
Series 01-1 (D)
Series 15-2 (D)
Rel. Position
Time [min]
oriC
oriB
terB

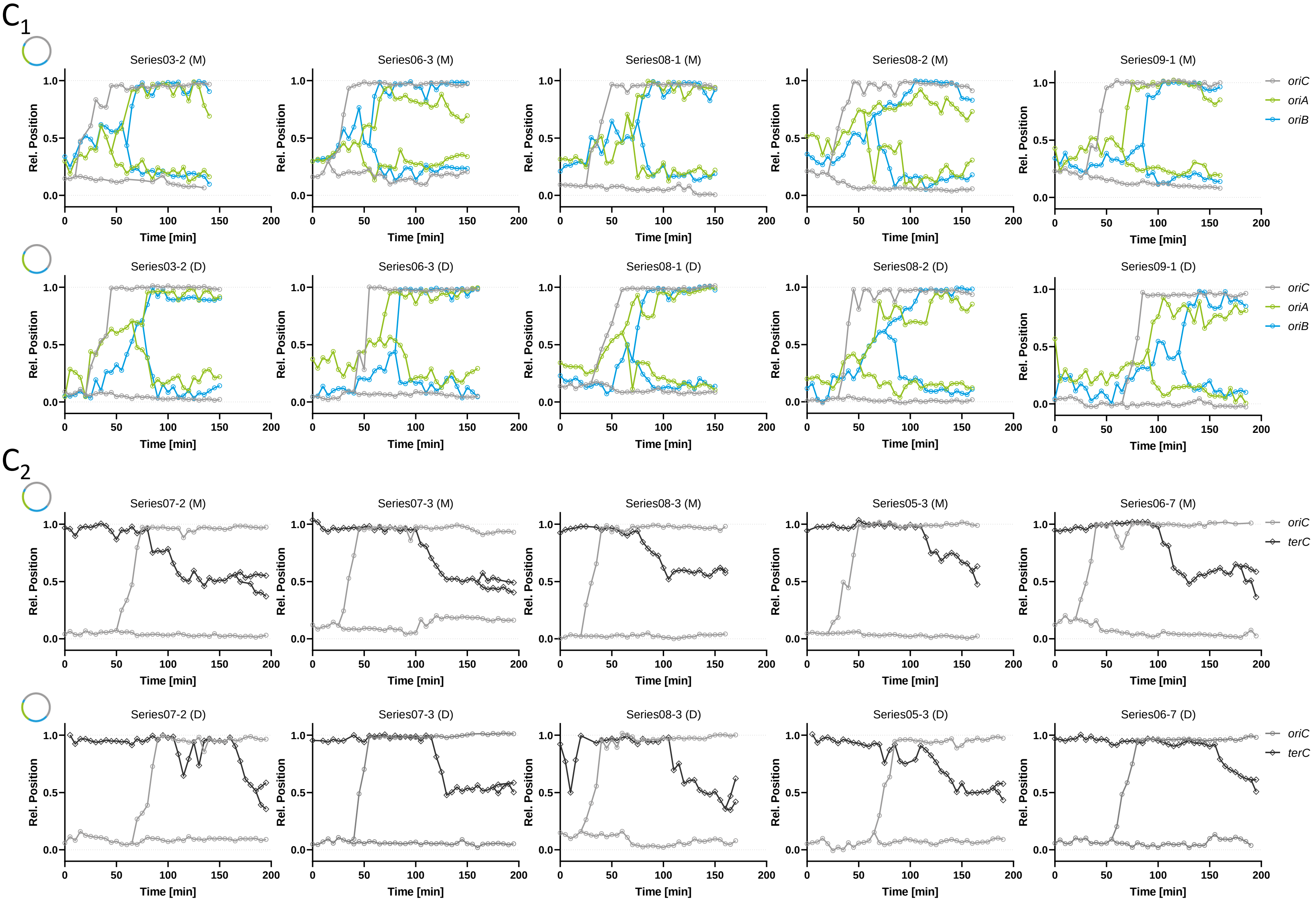
C1
Series03-2 (M)
Series06-3 (M)
Series08-1 (M)
Series08-2 (M)
Series09-1 (M)
Series03-2 (D)
Series06-3 (D)
Series08-1 (D)
Series08-2 (D)
Series09-1 (D)
oriC
oriA
oriB
C2
Series07-2 (M)
Series07-3 (M)
Series08-3 (M)
Series05-3 (M)
Series06-7 (M)
Series07-2 (D)
Series07-3 (D)
Series08-3 (D)
Series05-3 (D)
Series06-7 (D)
oriC
terC
Rel. Position
Time [min]

$C_3$

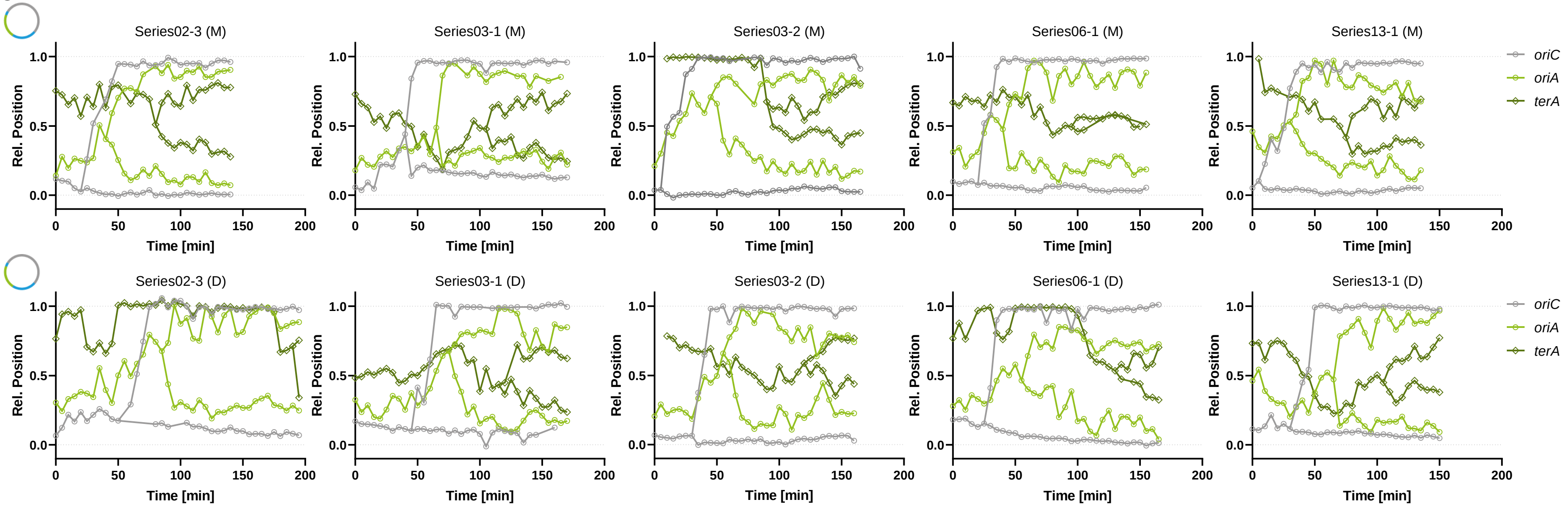
Series02-3 (M)
Series03-1 (M)
Series03-2 (M)
Series06-1 (M)
Series13-1 (M)
Series02-3 (D)
Series03-1 (D)
Series03-2 (D)
Series06-1 (D)
Series13-1 (D)
Rel. Position
Time [min]
oriC
oriA
terA

$C_4$

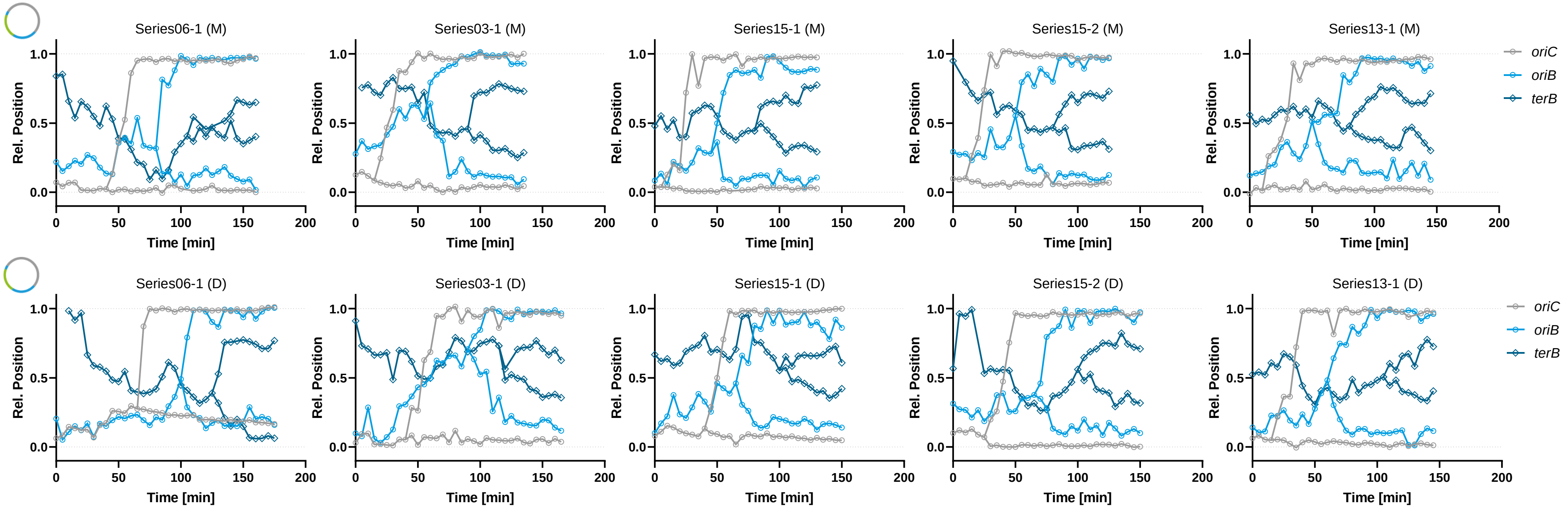
Series06-1 (M)
Series03-1 (M)
Series15-1 (M)
Series15-2 (M)
Series13-1 (M)
Series06-1 (D)
Series03-1 (D)
Series15-1 (D)
Series15-2 (D)
Series13-1 (D)
Rel. Position
Time [min]
oriC
oriB
terB

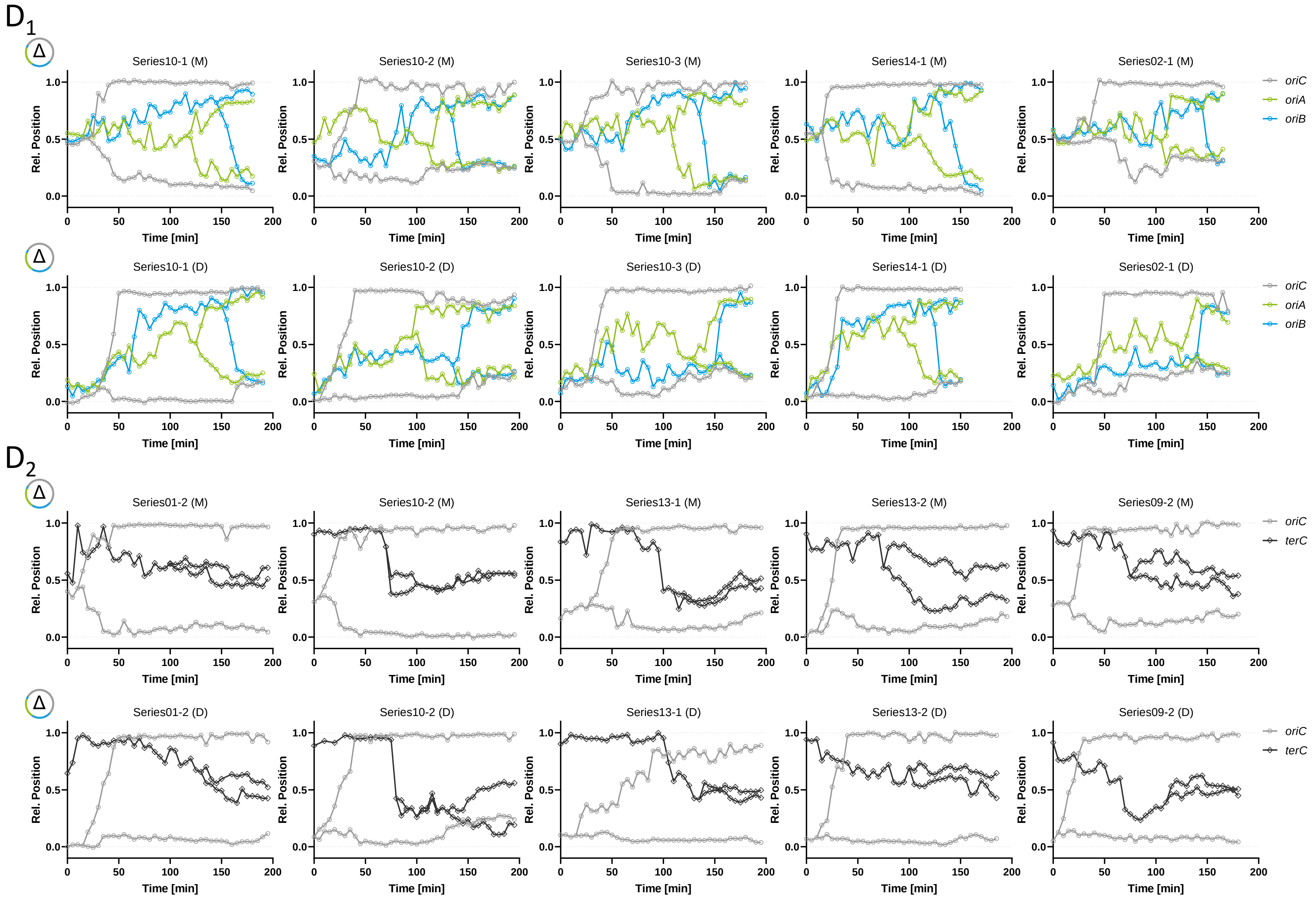
D1
Series10-1 (M)
Series10-2 (M)
Series10-3 (M)
Series14-1 (M)
Series02-1 (M)
oriC
oriA
oriB
Series10-1 (D)
Series10-2 (D)
Series10-3 (D)
Series14-1 (D)
Series02-1 (D)
D2
Series01-2 (M)
Series10-2 (M)
Series13-1 (M)
Series13-2 (M)
Series09-2 (M)
oriC
terC
Series01-2 (D)
Series10-2 (D)
Series13-1 (D)
Series13-2 (D)
Series09-2 (D)
Rel. Position
Time [min]
Δ

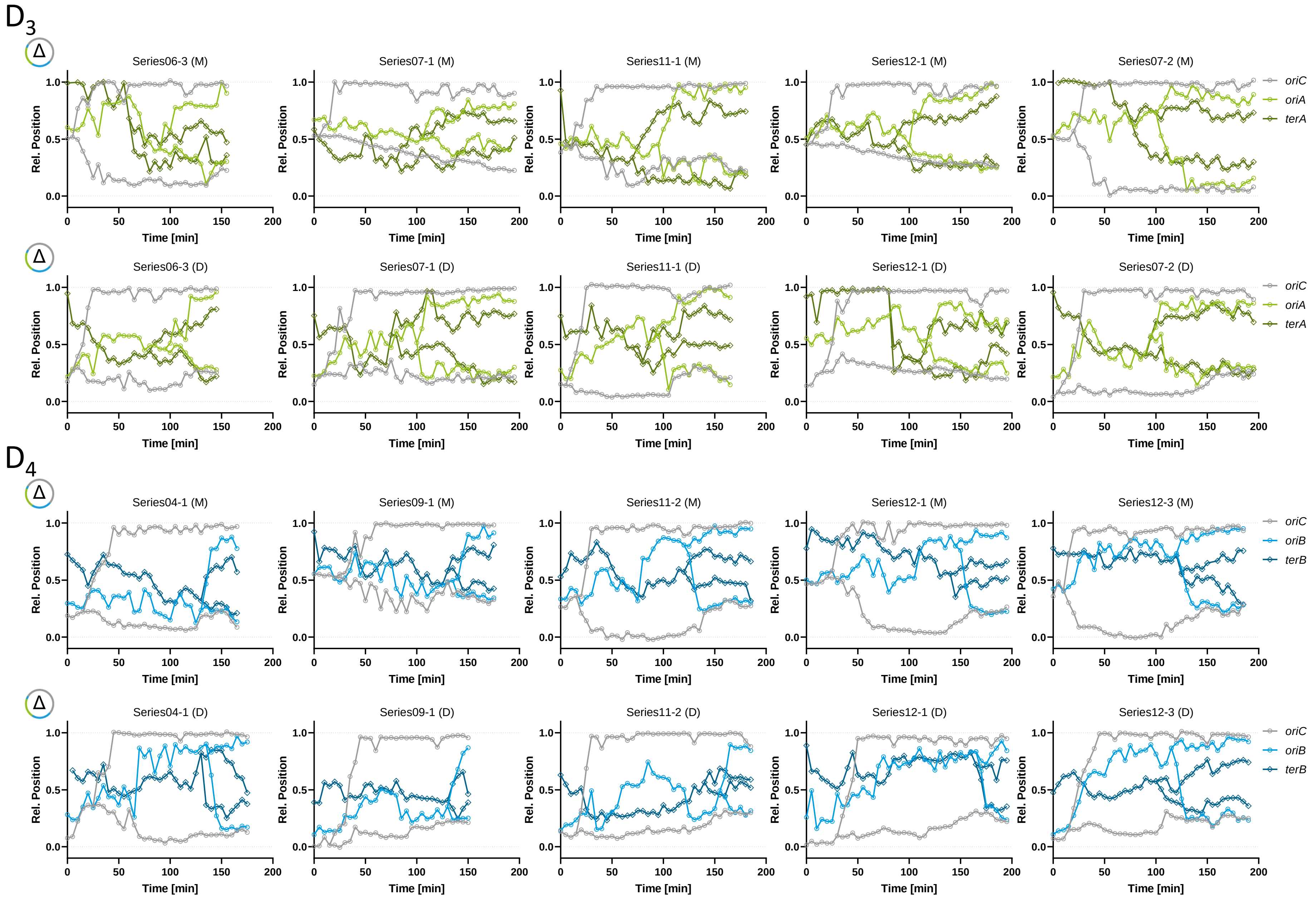
D3
Series06-3 (M)
Series07-1 (M)
Series11-1 (M)
Series12-1 (M)
Series07-2 (M)
Series06-3 (D)
Series07-1 (D)
Series11-1 (D)
Series12-1 (D)
Series07-2 (D)
oriC
oriA
terA
D4
Series04-1 (M)
Series09-1 (M)
Series11-2 (M)
Series12-1 (M)
Series12-3 (M)
Series04-1 (D)
Series09-1 (D)
Series11-2 (D)
Series12-1 (D)
Series12-3 (D)
oriC
oriB
terB
Rel. Position
Time [min]
Δ

**S1 Text. Model of compacted DNA.**

For the simulations used in this study we employed stochastic Monte Carlo computer simulations to generate ensembles of DNA configurations in the bacterial cell. In order to do so, various important biological aspects of the bacterial chromosome and its organization had to be considered.

The first important fact is that the length of bacterial chromosomes is about three orders of magnitude larger than the cell (Wang et al., 2013; Badrinarayanan et al., 2015). Thus, cells have to massively compact their DNA in a manner that is compatible with DNA replication, DNA repair and further cellular processes (Badrinarayanan et al., 2015). While eukaryotes use histone proteins around which DNA is wrapped to form nucleosomes, bacteria deploy a combination of different mechanisms to achieve this task (Stavans and Oppenheim, 2006). The most prominent ones include entropic forces as a result of macromolecular crowding (Heermann, 2011) (Jun, 2015), the partition of long DNA molecules into so-called supercoiled domains (Wang et al., 2013), and finally, the association of the DNA with nucleoid-associated proteins (NAPs) (Stavans and Oppenheim, 2006). The combination of these mechanisms leads to an organization of the chromosome into various domains on different length scales (Wang et al., 2013; Badrinarayanan et al., 2015; Marbouty et al., 2015; Wu et al., 2019).

For our simulations we used the model for DNA from Buenemann and Lenz, 2010. Here, the compaction of DNA is implemented by modeling the DNA as a semi-flexible polymer which locally has the shape of a sphere (called “blob”), due to the interactions with compaction proteins and negative DNA supercoiling. Thus, we can view the chromosome as an entropic spring of blobs with each blob representing a structural unit of the chromosome consisting of supercoiled DNA stabilized by DNA-binding proteins (Jun and Wright, 2010). Examples for such compaction proteins are the NAPs IHF and HU which are identified as key architectural proteins in prokaryotes (Swinger and Rice, 2004). IHF binds without cooperativity to ~35 bp under various experimental conditions and HU has been reported to bind to binding sites of between 9 and 42 bp (Stavans and Oppenheim, 2006). While there is no direct evidence for the involvement of specific architectural proteins in chromosomal interaction domain (CID) boundary formation in bacteria, the involvement of the NAPs FIS and H-NS has been suggested for microdomain formation in *E. coli* (Brocken et al., 2018).

Therefore, blobs define topological domains which are insulated from each other as described in Postow et al., 2004 and Wang et al., 2013. It was suggested that the supercoiled domains vary in with an average in the order of magnitude of 10 kb (Postow et al., 2004; Wang et al., 2013).

In our model we assume, that a blob typically contains one DNA-loop of a given loop size $l$. With the length of one bp of DNA $b$ = 0.34 nm we can calculate the blob radius as the radius of gyration

$$r_b = \frac{\sqrt{l} * b}{\sqrt{6}}$$

If we insert the above given loop size of approximately 10 kbp we receive a blob diameter of $d_b$ ~30 nm. Within this study we found the best agreement with the experimental data for a blob diameter of 10 nm which corresponds to a loop size of 1298 bp. Hence, the compacted chromosome can be represented as a chain of such blobs.

An important fact to notice here is that the organization of the chromosome into such domains is dynamic and hence the overall conformation of a chromosome is dynamic (Heermann, 2011; Wu et al., 2019). Thus, any model has to find a way to sample the conformation space ergodically in order to calculate meaningful ensemble averages for chromosome configurations.

**S2 Text. Monte Carlo sampling of configuration space.**

The goal of our model is to obtain an ergodic ensemble of possible configurations of the fused plasmid within the following geometric constraints.

(i) The first obvious restriction is the cell itself, within which a DNA molecule must be located. Confinement within the cell is an important aspect both for segregation and organization of the bacterial chromosome (Jun et al., 2007; Wiggins et al., 2010). It was shown that confinement manifests itself in the shape of the position distribution function (Wiggins et al., 2010). In our simulations the cellular volume is discretized and represented by a three-dimensional cubic lattice of dimensions *H* x ¼ *H* x ¼ *H* with *H* being the length of the cell. In our simulations we chose H to be between 1.3 µm - 2 µm which corresponds to the experimentally measured cell lengths.

(ii) The second constraint that we introduce into our model is self-avoidance of the chromosome. As described in Text S1, we represent the compacted chromosome as a chain of blobs. In our model such a chromosome is represented as a random walk on a three-dimensional lattice with the lattice spacing being exactly the diameter of a blob, $d_b$. Since the blobs contain extended pieces of DNA it is important to implement self-avoidance because two compact structures cannot occupy the same spatial position (Heermann, 2011). To take self-avoidance into account, each lattice point can only be occupied by one blob (Buenemann and Lenz, 2010).

(iii) A third constraint imposed on the chromosome is the spatial fixation of DNA loci within the cell. This results in the task of assembling the DNA molecule from a series of random walks that connect the individual fix-points.

(iv) The fourth and last constraint implemented in the model is given by the enrichment-region that we introduce for *terB*. Here, we do not want to strictly fixate the DNA locus to a distinct spatial position but to select configurations from our ensemble in which *terB* is located within the defined region.

We use the following protocol to compute an ensemble of configurations fulfilling the constraints:

1. Construct a single random walk connecting the selected fixpoints and fulfilling the constraints (i)-(iii).

2. Use the MOS-algorithm (Madras et al., 1990) to compute an ergodic ensemble of configurations fulfilling the constraints (i)-(iii).

3. Post-process the ensemble from 2. and select only those configurations which also fulfill constraint (iv).

For the construction of one initial random walk we require a robust method to connect given fix-points by a random walk of fixed length. Therefore, we implemented the A*-search algorithm. This is a path search algorithm often used because of its completeness, optimality and optimal efficiency (Russell and Norvig, 2018). It enables us to select any number of desired fix-points on the lattice and connect them with a random walk. Thereafter, we have to construct an ergodic ensemble of self-avoiding walks within the constraints (i)-(iii) from our initial walk. To address this task a simulation scheme based on the MOS method was used by which the configuration space of random walks of fixed length and fixed endpoints can be sampled. (Madras et al., 1990). Finally, we can use this ergodic ensemble and select the configurations which also fulfill constraint (iv). Thereby, we are able to produce an ensemble of DNA configurations with of our coarse-grained model that also takes molecular details (i.e. the local action of the compaction proteins) into account and is able to reflect cell-to-cell variance.

### S3 Text. SI References (1/2)

Badrinarayanan A, Le TB, Laub MT. Bacterial chromosome organization and segregation. *Annu Rev Cell Dev Biol*. 2015;31:171-99. doi: 10.1146/annurev-cellbio-100814-125211.

Becker A, Schmidt M, Jäger W, Pühler A. New gentamicin-resistance and lacZ promoter-probe cassettes suitable for insertion mutagenesis and generation of transcriptional fusions. *Gene*. 1995 Aug 30;162(1):37-9. doi: 10.1016/0378-1119(95)00313-u.

Bigot S, Saleh OA, Lesterlin C, Pages C, El Karoui M, Dennis C, Grigoriev M, Allemand JF, Barre FX, Cornet F. KOPS: DNA motifs that control *E. coli* chromosome segregation by orienting the FtsK translocase. *EMBO J*. 2005 Nov 2;24(21):3770-80. doi: 10.1038/sj.emboj.7600835.

Brocken JW, Tark-Dame M, Dame RT. The organization of bacterial genomes: Towards understanding the interplay between structure and function. *Curr Opin Syst Biol*. 2018 Apr,8:137-143, ISSN 2452-3100, https://doi.org/10.1016/j.coisb.2018.02.007.

Buenemann M, Lenz P. A geometrical model for DNA organization in bacteria. *PLoS One*. 2010 Nov 3;5(11):e13806. doi: 10.1371/journal.pone.0013806.

Cervantes-Rivera R, Pedraza-López F, Pérez-Segura G, Cevallos MA. The replication origin of a *repABC* plasmid. *BMC Microbiol*. 2011 Jun 30;11:158. doi: 10.1186/1471-2180-11-158.

Döhlemann J, Brennecke M, Becker A. Cloning-free genome engineering in *Sinorhizobium meliloti* advances applications of Cre/*loxP* site-specific recombination. *J Biotechnol*. 2016 Sep 10;233:160-70. doi: 10.1016/j.jbiotec.2016.06.033.

Döhlemann J, Wagner M, Happel C, Carrillo M, Sobetzko P, Erb TJ, Thanbichler M, Becker A. A family of single copy *repABC*-type shuttle vectors stably maintained in the Alpha-proteobacterium *Sinorhizobium meliloti*. *ACS Synth Biol*. 2017 Jun 16;6(6):968-984. doi: 10.1021/acssynbio.6b00320.

Frage B, Döhlemann J, Robledo M, Lucena D, Sobetzko P, Graumann PL, Becker A. Spatiotemporal choreography of chromosome and megaplasmids in the *Sinorhizobium meliloti* cell cycle. *Mol Microbiol*. 2016 Jun;100(5):808-23. doi: 10.1111/mmi.13351.

Hanahan D. Studies on transformation of *Escherichia coli* with plasmids. *J Mol Biol*. 1983 Jun 5;166(4):557-80. doi: 10.1016/s0022-2836(83)80284-8.

Harrison CL, Crook MB, Peco G, Long SR, Griffitts JS. Employing site-specific recombination for conditional genetic analysis in *Sinorhizobium meliloti*. *Appl Environ Microbiol*. 2011 Jun;77(12):3916-22. doi: 10.1128/AEM.00544-11.

Heermann DW. Physical nuclear organization: loops and entropy. *Curr Opin Cell Biol*. 2011 Jun;23(3):332-7. doi: 10.1016/j.ceb.2011.03.010.

Hendrickson H, Lawrence JG. Selection for chromosome architecture in bacteria. *J Mol Evol*. 2006 May;62(5):615-29. doi: 10.1007/s00239-005-0192-2.

Jun S. Chromosome, cell cycle, and entropy. *Biophys J*. 2015 Feb 17;108(4):785-786. doi: 10.1016/j.bpj.2014.12.032.

Jun S, Arnold A, Ha BY. Confined space and effective interactions of multiple self-avoiding chains. *Phys Rev Lett*. 2007 Mar 23;98(12):128303. doi: 10.1103/PhysRevLett.98.128303.

Jun S, Wright A. Entropy as the driver of chromosome segregation. *Nat Rev Microbiol*. 2010 Aug;8(8):600-7. doi: 10.1038/nrmicro2391.

Khan SR, Gaines J, Roop RM 2nd, Farrand SK. Broad-host-range expression vectors with tightly regulated promoters and their use to examine the influence of TraR and TraM expression on Ti plasmid quorum sensing. *Appl Environ Microbiol*. 2008 Aug;74(16):5053-62. doi: 10.1128/AEM.01098-08.

Lau IF, Filipe SR, Søballe B, Økstad OA, Barre FX, Sherratt DJ. Spatial and temporal organization of replicating *Escherichia coli* chromosomes. *Mol Microbiol*. 2003 Aug;49(3):731-43. doi: 10.1046/j.1365-2958.2003.03640.x.

Madras N, Orlitsky A & Shepp LA. Monte Carlo generation of self-avoiding walks with fixed endpoints and fixed length. *J Stat Phys*. 1990 Jan; 58:159-183. https://doi.org/10.1007/BF01020290

Marbouty M, Le Gall A, Cattoni DI, Cournac A, Koh A, Fiche JB, Mozziconacci J, Murray H, Koszul R, Nollmann M. Condensin- and replication-mediated bacterial chromosome folding and origin condensation revealed by Hi-C and super-resolution imaging. *Mol Cell*. 2015 Aug 20;59(4):588-602. doi: 10.1016/j.molcel.2015.07.020.

### S3 Text. SI References (2/2)

Marchler-Bauer A, Bo Y, Han L, He J, Lanczycki CJ, Lu S, Chitsaz F, Derbyshire MK, Geer RC, Gonzales NR, Gwadz M, Hurwitz DI, Lu F, Marchler GH, Song JS, Thanki N, Wang Z, Yamashita RA, Zhang D, Zheng C, Geer LY, Bryant SH. CDD/SPARCLE: functional classification of proteins via subfamily domain architectures. *Nucleic Acids Res*. 2017 Jan 4;45(D1):D200-D203. doi: 10.1093/nar/gkw1129.

Marchler-Bauer A, Derbyshire MK, Gonzales NR, Lu S, Chitsaz F, Geer LY, Geer RC, He J, Gwadz M, Hurwitz DI, Lanczycki CJ, Lu F, Marchler GH, Song JS, Thanki N, Wang Z, Yamashita RA, Zhang D, Zheng C, Bryant SH. CDD: NCBI's conserved domain database. *Nucleic Acids Res*. 2015 Jan;43(Database issue):D222-6. doi: 10.1093/nar/gku1221.

Marchler-Bauer A, Lu S, Anderson JB, Chitsaz F, Derbyshire MK, DeWeese-Scott C, Fong JH, Geer LY, Geer RC, Gonzales NR, Gwadz M, Hurwitz DI, Jackson JD, Ke Z, Lanczycki CJ, Lu F, Marchler GH, Mullokandov M, Omelchenko MV, Robertson CL, Song JS, Thanki N, Yamashita RA, Zhang D, Zhang N, Zheng C, Bryant SH. CDD: a Conserved Domain Database for the functional annotation of proteins. *Nucleic Acids Res*. 2011 Jan;39(Database issue):D225-9. doi: 10.1093/nar/gkq1189.

Postow L, Hardy CD, Arsuaga J, Cozzarelli NR. Topological domain structure of the *Escherichia coli* chromosome. *Genes Dev*. 2004 Jul 15;18(14):1766-79. doi: 10.1101/gad.1207504.

Russell SJ and Norvig P. Artificial intelligence: a modern approach. Pearson education limited (2018).

Schäfer A, Tauch A, Jäger W, Kalinowski J, Thierbach G, Pühler A. Small mobilizable multi-purpose cloning vectors derived from the *Escherichia coli* plasmids pK18 and pK19: selection of defined deletions in the chromosome of *Corynebacterium glutamicum*. *Gene*. 1994 Jul 22;145(1):69-73. doi: 10.1016/0378-1119(94)90324-7.

Schlüter JP, Czuppon P, Schauer O, Pfaffelhuber P, McIntosh M, Becker A. Classification of phenotypic subpopulations in isogenic bacterial cultures by triple promoter probing at single cell level. *J Biotechnol*. 2015 Mar 20;198:3-14. doi: 10.1016/j.jbiotec.2015.01.021.

Schlüter JP, Reinkensmeier J, Barnett MJ, Lang C, Krol E, Giegerich R, Long SR, Becker A. Global mapping of transcription start sites and promoter motifs in the symbiotic α-proteobacterium *Sinorhizobium meliloti* 1021. *BMC Genomics*. 2013 Mar 7;14:156. doi: 10.1186/1471-2164-14-156.

Sibley CD, MacLellan SR, Finan T. The *Sinorhizobium meliloti* chromosomal origin of replication. *Microbiology* (Reading). 2006 Feb;152(Pt 2):443-455. doi: 10.1099/mic.0.28455-0.

Simon R, Priefer U, Pühler A. A broad host range mobilization system for *in vivo* genetic engineering: transposon mutagenesis in Gram negative bacteria. *Nat Biotechnol*. 1983 Nov;1(9):784-791. doi: 10.1038/nbt1183-784

Stavans J, Oppenheim A. DNA-protein interactions and bacterial chromosome architecture. *Phys Biol*. 2006 Dec 22;3(4):R1-10. doi: 10.1088/1478-3975/3/4/R01. PMID: 17200598.

Swinger KK, Rice PA. IHF and HU: flexible architects of bent DNA. *Curr Opin Struct Biol*. 2004 Feb;14(1):28-35. doi: 10.1016/j.sbi.2003.12.003.

Tauch A, Zheng Z, Pühler A, Kalinowski J. *Corynebacterium striatum* chloramphenicol resistance transposon Tn5564: genetic organization and transposition in *Corynebacterium glutamicum*. *Plasmid*. 1998 Sep;40(2):126-39. doi: 10.1006/plas.1998.1362.

Wang X, Montero Llopis P, Rudner DZ. Organization and segregation of bacterial chromosomes. *Nat Rev Genet*. 2013 Mar;14(3):191-203. doi: 10.1038/nrg3375.

Weber E, Engler C, Gruetzner R, Werner S, Marillonnet S. A modular cloning system for standardized assembly of multigene constructs. *PLoS One*. 2011 Feb 18;6(2):e16765. doi: 10.1371/journal.pone.0016765.

Wiggins PA, Cheveralls KC, Martin JS, Lintner R, Kondev J. Strong intranucleoid interactions organize the *Escherichia coli* chromosome into a nucleoid filament. *Proc Natl Acad Sci U S A*. 2010 Mar 16;107(11):4991-5. doi: 10.1073/pnas.0912062107.

Wu F, Japaridze A, Zheng X, Wiktor J, Kerssemakers JWJ, Dekker C. Direct imaging of the circular chromosome in a live bacterium. *Nat Commun*. 2019 May 16;10(1):2194. doi: 10.1038/s41467-019-10221-0.
